# Redefining the topology of the human bone marrow using augmented spatial transcriptomic analysis

**DOI:** 10.64898/2025.12.22.694222

**Authors:** Rosalin A Cooper, Emily Thomas, Muhammad Dawood, Hosuk Ryou, Anna Sozanska, Carlo Pescia, Oliver McCallion, Muskaan Gupta, Renuka Teague, Joanna Hester, Fadi Issa, Dan J Woodcock, Bethan Psaila, Adam Mead, Jens Rittscher, Daniel Royston

**Affiliations:** Nuffield Division of Clinical Laboratory Sciences, University of Oxford, Oxford, UK; Oxford University Hospitals NHS Foundation Trust, Oxford, UK; Institute of Biomedical Engineering (IBME), Department of Engineering Science, University of Oxford, UK; Big Data Institute / Li Ka Shing Centre for Health Information and Discovery, University of Oxford, UK; Unit of Pathology, ASST Santi Paolo e Carlo, Milan, Italy; Nuffield Department of Surgical Sciences, University of Oxford, Oxford, UK; Chinese Academy of Medical Sciences Oxford Institute, University of Oxford, Oxford, UK; Oxford Centre for Histopathological Research, University of Oxford, Oxford, UK; MRC Molecular Haematology Unit, MRC Weatherall Institute for Molecular Medicine, Radcliffe Department of Medicine and National Institute of Health Research Centre, University of Oxford, Oxford, UK; Ludwig Institute for Cancer Research (Oxford Branch), University of Oxford, Oxford, UK

**Author notes:** Contributed equally (Rosalin A Cooper, Emily Thomas, Muhammad Dawood, Jens Rittscher, Daniel Royston).

## Abstract

The bone marrow (BM) is the main site of haematopoiesis in adult life. Our understanding of the pathogenesis of BM-derived blood cancers is limited by lack of spatial contextualisation. While emerging spatial transcriptomic (ST) platforms offer unprecedented opportunities for spatially-resolved cellular phenotyping, we recognise and incorporate the power of AI-based tissue feature detection to enhance ST workflows. We perform ST analysis to define the topology of the normal bone marrow (BM) and BM in myeloproliferative neoplasms (MPNs), profiling 5,104,452 cells across 30 human BM samples. Following rigorous histology-based QC, we identify spatially-restricted trajectories of haematopoiesis and extend our understanding of the haematopoietic stem cell (HSC) niche. We find that BM fibrosis in MPN is associated with expansion of distinct immune and stromal co-enriched cell neighbourhoods. We then present a machine learning (ML)-based model trained on ST data that quantifies BM microenvironmental deviation, identifying heretofore unrecognised inter- and intra-individual sample heterogeneity in MPN. Our study demonstrates the potential for AI-based augmented ST analysis, and redefines our understanding of human BM topology.

## INTRODUCTION

The bone marrow is the main site of haematopoiesis in adults and comprises a microenvironment rich in haematopoietic, immune and supporting stromal populations. Single-cell RNA sequencing (scRNA-seq)-based approaches have been instrumental in furthering our understanding of haematopoiesis and the perturbation in BM-derived blood cancers^1–3^. However, our understanding of disease pathogenesis is limited by the lack of spatially-resolved descriptions of the human BM. Emerging spatial transcriptomic (ST) approaches such as Xenium offer up to sub-cellular resolution providing comprehensive spatially-resolved topological mapping^4^. However, ST-based approaches do not detect features such as bone, adipocytes and fibrosis that influence haematopoiesis in health and disease. To address this, we have developed an augmented BM ST data analysis workflow. Unlike prior BM spatial studies^5–9^, our approach combines expert morphology-driven QC with multiple AI-based feature detectors and high-resolution Xenium ST across full tissue sections. We demonstrate the importance of histological annotation for quality control (QC) and employ multiple AI-based algorithms for feature detection (bone, megakaryocytes, adipocytes) and quantitation (fibrosis), enabling architectural features invisible to ST alone to be incorporated into spatial modelling. We then use supervised and unsupervised spatial analyses to exhaustively map the normal human BM. We identify distinct cellular topological niches, spatially-restricted patterns of haematopoiesis and provide novel insights into the haematopoietic stem cell (HSC) niche.

We then demonstrate how our augmented ST-analysis can be applied to BM-derived blood cancers to capture microenvironmental perturbation, using myeloproliferative neoplasms (MPNs) as an exemplar disease model. MPN is a highly tractable model of disease evolution; a myeloid blood cancer characterised by well-defined driver mutations (in *JAK2, CALR* or *MPL*), activation in JAK-STAT signalling and aberrant proliferation of myeloid, erythroid or megakaryocyte populations^10^. However, clinically MPNs are phenotypically highly variable, with essential thrombocythaemia (ET) and polythaemia vera (PV) characterised by thrombocytosis and elevated haemoglobin respectively, and an increased risk of thrombo-embolic events; while myelofibrosis (MF) is characterised by BM fibrosis and a poor prognosis^11^. Single cell RNA-seq (scRNA-seq)-based studies implicate interactions between the neoplastic haematopoietic clone, stromal and immune cells in the development of BM fibrosis^12–14^. However, spatial contextualisation of these interactions is lacking. Unlike other myeloid blood cancers for which molecular markers are utilised for prognostication and to monitor disease evolution, in MPN disparate clinical phenotypes and trajectories are associated with molecular commonality^3^. The extent to which this clinical variability is underpinned by heterogeneous microenvironmental topology is not yet fully defined. To address this, by applying our data-driven augmented ST analysis workflow we quantitatively map the perturbation seen in MPN. We employ our recently developed ‘band descriptor’ to the BM for the first time to identify novel spatial microenvironmental signatures, or ‘spatiotypes’, that define the BM in health and in MPN. Finally, we present a newly-developed ranking-based machine learning (ML) model that quantitatively captures previously undescribed cohort and intra-individual BM microenvironmetal variability in MPN. Together, these approaches constitute a new analytical paradigm for human BM spatial biology and redefine our understanding of human BM topology.

## RESULTS

### Comprehensive quantitative descriptions of the human bone marrow microenvironment using spatial transcriptomics

We developed a workflow for the analysis of ST data from human archival diagnostic formalin-fixed paraffin-embedded (FFPE) bone marrow trephine (BMT) material profiled with the 10x Xenium platform (10x Genomics) (Fig. 1). We analysed a total of 30 BMT FFPE samples from 30 individuals, profiling 244,714,040 transcripts within 5,104,452 cells. These included five BMTs with no evidence of haematological malignancy, three cases of PV, eight ET cases, three Pre-PMF cases (a non-fibrotic but ‘high-risk’ MPN subtype) and 11 MF BMTs (Supplementary Fig. 1&2).

**Figure 1:**
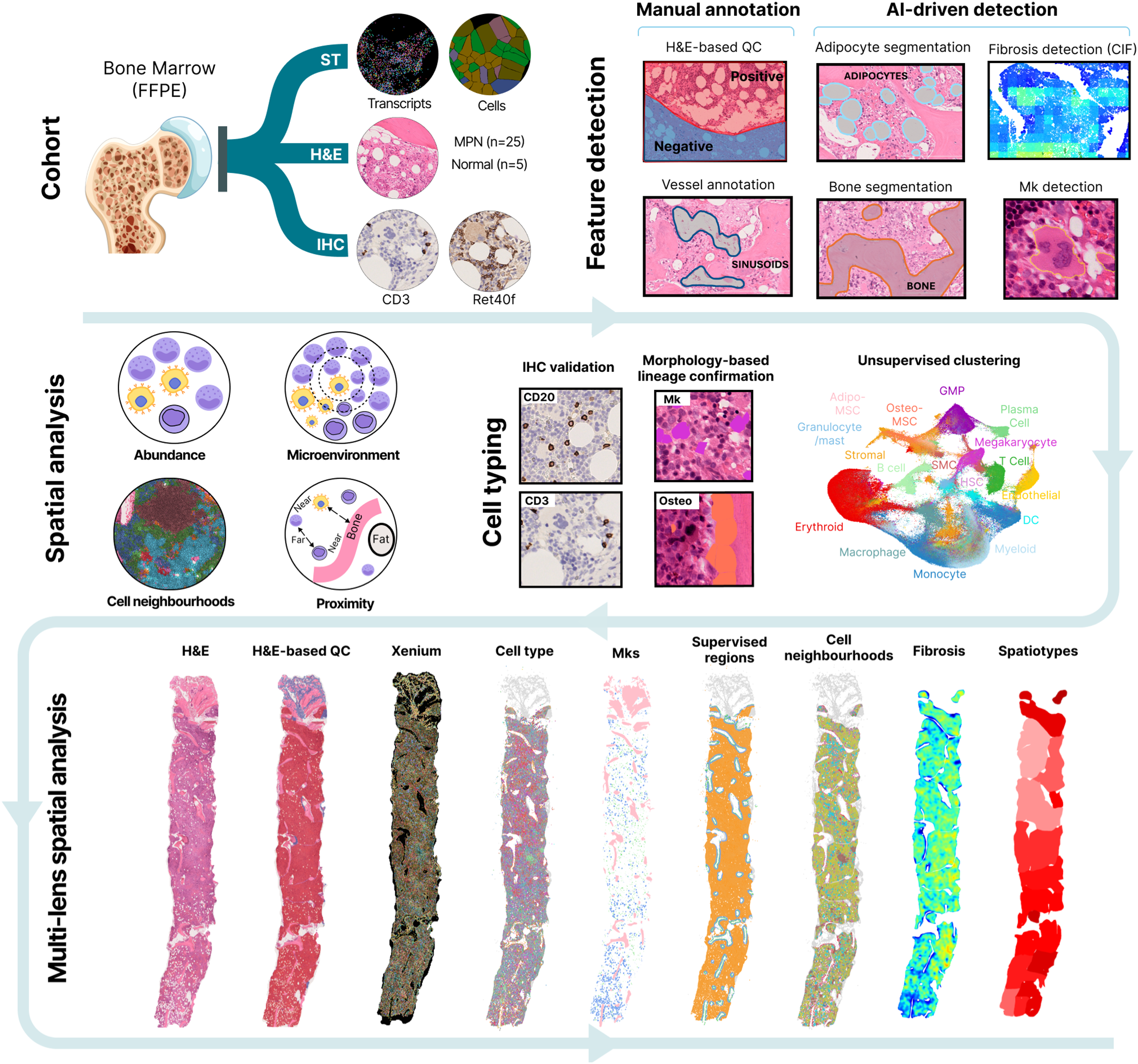
AI-augmented ST bone marrow analysis workflow. Study workflow: Spatial transcriptomic (ST) data, immunohistochemical (IHC) and H&E images from FFPE bone marrow trephine (BMT) material, including five normal samples and 25 from patients with MPN. Feature detection using both manual annotation and AI-based detection/segmentation of bone, adipocytes, megakaryocytes and quantitation of fibrosis was performed. Cell typing/annotation was performed with unsupervised clustering with subsequent validation via correlation between ST-identified cells and those detected on immunohistochemistry-based quantification. Spatial analyses included differential abundance analysis, unsupervised cell neighbourhood (CN) analysis, proximity (point pattern) analysis and interrogation of microenvironmental variability using a ‘band descriptor’ and development of a novel machine learning (ML)-based model. The workflow provides a novel multi-lens analytical paradigm for human BM spatial biology.

The Xenium platform is tissue preserving and a post-hoc H&E - on the same section - allows for direct integration with ST data for morphology-driven H&E-based quality control (QC). As we recently describe^15^, poorly preserved tissue regions (e.g. comprising crushed cells or folded tissue) were not detected on inspection of Xenium ST data, and were only evident on the H&E-stained histological image (Extended Data Fig. 1A&B). Despite this, histology-based annotation is not a routine part of most ST analysis workflows and its importance has largely gone unrecognised. We removed ‘cytomorphologically unpreserved’ regions from downstream analysis (n = 464,793 cells, 8.1%) (Extended Data Fig. 2, Supplementary Fig. 3, Methods). Some regions were cytomorphologically but not architecturally preserved (Extended Data Fig. 1B). Spatial analysis within such regions is not biologically meaningful. Regions of architectural preservation (broadly corresponding to an ‘intertrabecular space’, ITS) were therefore carefully annotated for downstream spatial analyses. We removed remaining cells with a low transcript number (<4 transcripts/cell, 180,784 cells, 3.1%). Notably, 83% of cells in regions of poor cytomorphological preservation were not identified by filtering according to transcript number (Extended Data Fig. 3A&B). This is particularly pertinent in BM material, in which tissue artefact and disruption is not uncommon. Notably, transcript density did not consistently correspond to regions of cellular cytomorphological or architectural preservation (Extended Data Fig. 1C), again demonstrating the importance of H&E-based spatial QC. We used the Xenium Onboard facility for cellular segmentation, but noted that megakaryocytes with large, polylobated nuclei, were ‘oversegmented’ (Extended Data Fig. 3C). To enhance the accuracy of megakaryocyte segmentation we therefore applied our previously developed AI-based megakaryocyte detection algorithm^16^ to the corresponding H&E image, inserting AI-segmented megakaryocytes (Extended Data Fig. 3C, Methods).

Together, through rigorous histology-based annotation and QC, transcript thresholding and AI-enhanced megakaryocyte segmentation, we removed 645,577 cells (11.2% of all nucleated cells) from downstream analysis. A total of 5,104,452 cells were taken forward for analysis with a median transcript number of 37 (Extended Data Fig. 3D).

To identify BM cell types, we used unsupervised expression-based clustering, annotation with scRNA-seq reference datasets, followed by H&E-based histological review (Supplementary Fig. 4, Methods). First, we identified hematopoietic stem cells (HSCs) as those co-expressing *CD34* and *KIT*. With remaining cells we performed unsupervised clustering (agnostic to spatial localisation), using differential expression analysis to identify cluster-defining markers (Fig. 2A&B, Table 1). To support lineage assignment, we annotated these clusters with human BM reference scRNA-seq datasets (Bandyopadhyay et al.^9^, Azimuth^17^) (Extended Data Fig. 3E). We identified an erythroid group expressing lineage-defining markers *GYPA* and *GYPB,* and a megakaryocyte group expressing *VWF* and *MMRN1* (Fig. 2B, Table 1). We identified dendritic cells (*IRF8, CD1C* and *CLEC10A*), monocytes (*CD14, SELL, CCR2, FCGR3A)* and macrophages (*CD163, CD68*). Typically lost in scRNA-seq-based studies due to conventional mononuclear cell isolation methods and challenging to capture with immunofluorescent (IF) imaging techniques, we identified a granulocyte/mast cell group expressing *KIT, CTSG, CPA3* and *FCER1A.* We also identified a group with an immature myeloid signature (expressing *FCN1, S100A12* and *MNDA*) we termed ‘myeloid’ and a population of immature myeloid progenitors best aligned to a granulocyte-monocyte progenitor (GMP) group (*KIT, CTSG)*. We identified plasma cells (*MZB1)* and lymphocytes, including T-cells (*CD3D, CD3E,* and *CD8A)* and B-cells (*CD19, MS4A1* and *CD79A)* (Fig. 2A-B, Table 1). Nearly a fifth of nucleated BM cells (18.1%) were non-haematological. These included endothelial cells (*PECAM1, CLEC14A* and *CD34);* smooth muscle cells (‘SMCs’) (*ACTA2, CNN1*) and a mesenchymal stromal cell (MSC) group (*PDGFRA, PDGFRB*) (Fig. 2A&B). Using the stromal-enriched scRNA-seq dataset generated by Bandyopadhyay et al.^9^, we performed supervised annotation of MSCs, identifying a group with an adipogenic gene signature (*LPL, ADIPOQ)*, ‘adipo-MSCs’, and a group with an osteogenic signature (*OGN, ASPN)* ‘osteo-MSCs’. All remaining MSC populations were grouped together (termed ‘stromal’) (Table 1). Haematopathologist-led visual inspection was then employed to confirm lineage assignment, overlaying the ST data on the H&E image to directly compare cell annotation with cellular morphology (Fig. 2C, Extended Data Fig. 4).

**Figure 2:**
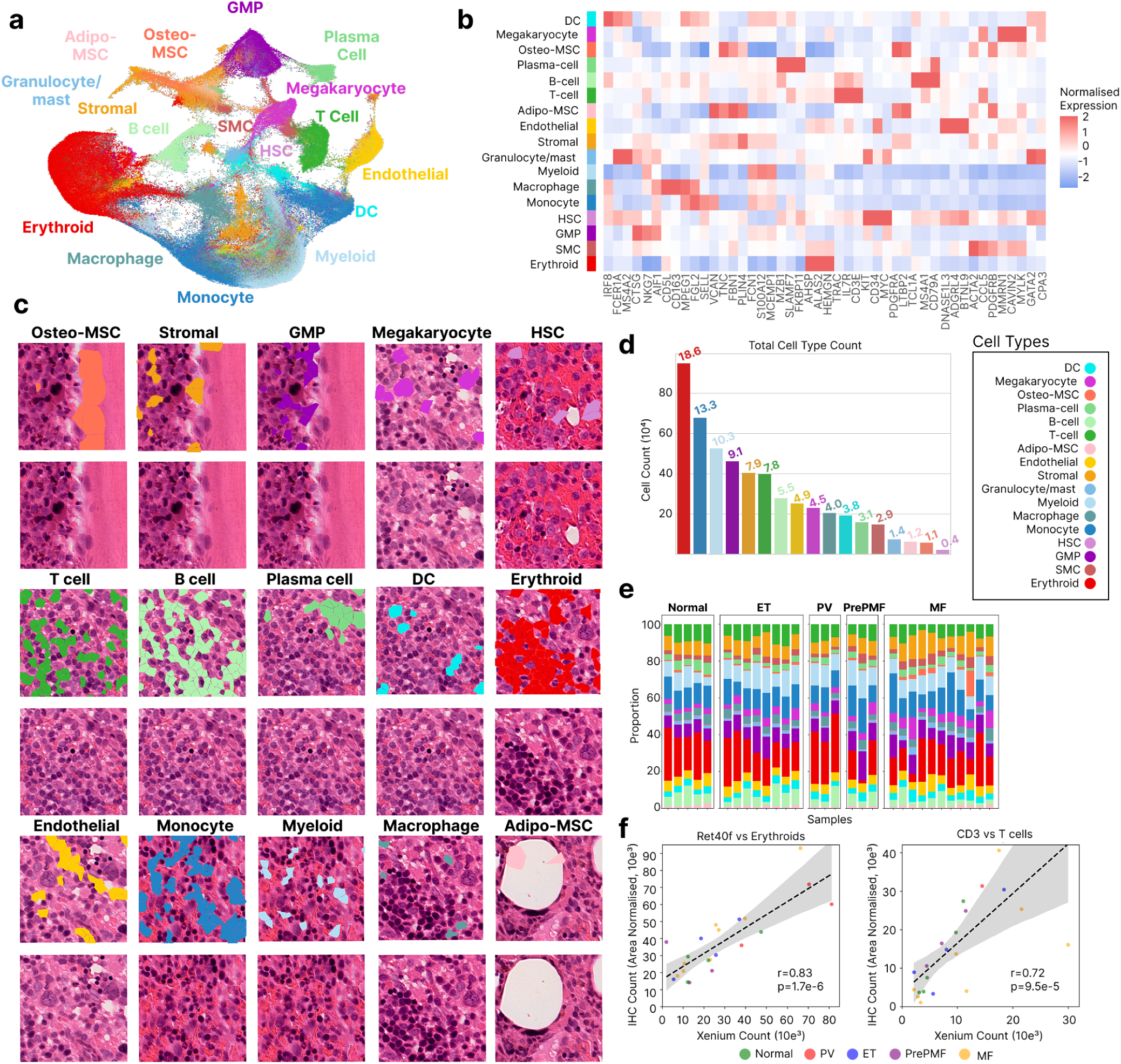
Cell type annotation. **A:** UMAP plot to show bone marrow (BM) cells coloured by cell type (randomly downsampled to 640,000 cells). **B:** Heatmap to show expression of key cluster-defining genes with the top log_2_ fold change (top 10 per cluster) across cell types (P_adj_ < 0.05, log_2_FC>1). The colour scale indicates normalised gene expression (red = high, blue = low). **C:** Panel to show cell types identified by the ST data overlying the H&E image. Each cell type is highlighted in a different colour (top image) and the corresponding H&E image is seen below. **D:** Bar plot to show the number of each BM cell type across all samples (n = 30 patients, n = 5,104,452 cells). The proportion of each cell type is indicated above each bar. **E:** Stacked bar plot to show the proportion of cell types per sample grouped by condition (Normal n = 5; PV n = 3; ET n = 8; PrePMF n = 3 and MF n = 11). **F:** Scatterplots to show correlation between the cell-type counts from ST data and area normalised counts from immunohistochemistry (IHC) quantification on sections from the same sample. Plots show correlation between erythroid cells and Ret40f (glycophorin C) IHC (n=22 samples) and T-cells and CD3 IHC (n=23 samples) (Pearson’s correlation).

Whole-section spatial profiling at single cell resolution with robust morphology-driven QC allows for accurate quantitation of cellular composition. Reflecting function, haematopoietic cells comprised 81.9% of all marrow cellular constituents (Fig. 2D&E). Indicative of optimal tissue preservation, we found that MSCs comprised a much greater proportion of cells compared to another recently published IF-based spatial map^9^, with osteo-MSCs comprising 1.1%, adipo-MSCs 1.2% and other stromal cells 7.9% (Fig. 2D&E). Unlike other spatial descriptions of the BM^9^, our sampling handling protocol^15^ minimised on-slide bone detachment, with osteo-MSCs seen to line the bone trabeculae (Fig. 2C). Likely reflecting lack of biases typically associated with liquid sampling and disaggregation in scRNA-seq-based studies^9^, the differential cellular abundance of both hematopoietic lineages and non-haematopoietic cells was largely consistent across the cohort (Fig. 2E). There was no association evident between cell type abundance and sex, age or driver mutation status (Supplementary Fig. 5A-D).

To further validate ST cell type assignment and quantitation, we correlated the number of cells assigned to a given cell type to the number of cells expressing a corresponding lineage-defining marker on immunohistochemistry (IHC) from serial FFPE sections from the same sample (see Methods). We performed automated quantification of IHC-detected cells on whole slide images (WSI) which revealed significant correlation between area normalised IHC and ST counts for CD3 with ST T-cells; CD20 with ST B-cells; and Ret40f (nucleated) with ST erythroids (Fig. 2F, Extended Data Fig. 3G&H).

### Quantitative spatial mapping of the normal human bone marrow identifies distinct topological niches

We generated a map of the normal BM across a total of 509,333 cells from five BMTs. Recognising the limitations of previously published studies regarding capture of BM-based architectural features such as bone, our previously developed AI-based image-analysis algorithm^18^ was used to segment bony trabeculae (Fig. 3A, Supplementary Fig. 6, Methods), and the computational tool *MarrowQuant 2.0* to segment adipocytes^19^ (Fig. 3A). Large vascular structures (arterioles and sinusoids) were manually annotated by a haematopathologist (Fig. 3A). We then used unsupervised cell neighbourhood (CN) analysis with *CellCharter*^20^ to identify distinct patterns of haematopoieis and associated microenvironmental support. Following spatial graph construction (see Methods, Supplementary Fig. 7&8), we identified ten spatial cellular neighbourhoods (CN0-9), characterised by mixed cell populations (Fig. 3B-D, Extended Data Fig. 5), many of which represented novel topological niches. Reflecting the role of megakaryocytes in maintaining BM vascular integrity^21^ we found that megakaryocytes and endothelial cells were co-enriched in CN9 (Fig. 3C). CN1 was enriched for macrophages and erythroid cells, in keeping with the key role of macrophages in supporting erythroid maturation^12,13^. We also identified a distinct immune niche, with CN5 characterised by co-enrichment of T-cells, B-cells and DCs. We identified CNs demonstrating divergent trajectories of myelopoiesis, with CN6 comprising a granulopoietic trajectory (GMPs, myeloid cells, granulocytes/mast cells), while CN8 represented an antigen presenting cell trajectory (GMPs, monocytes, macrophages and dendritic cells). Although granulocytes and antigen presenting cells share a common progenitor (GMPs), this demonstrates that these divergent patterns of differentiation are spatially discrete.

**Figure 3:**
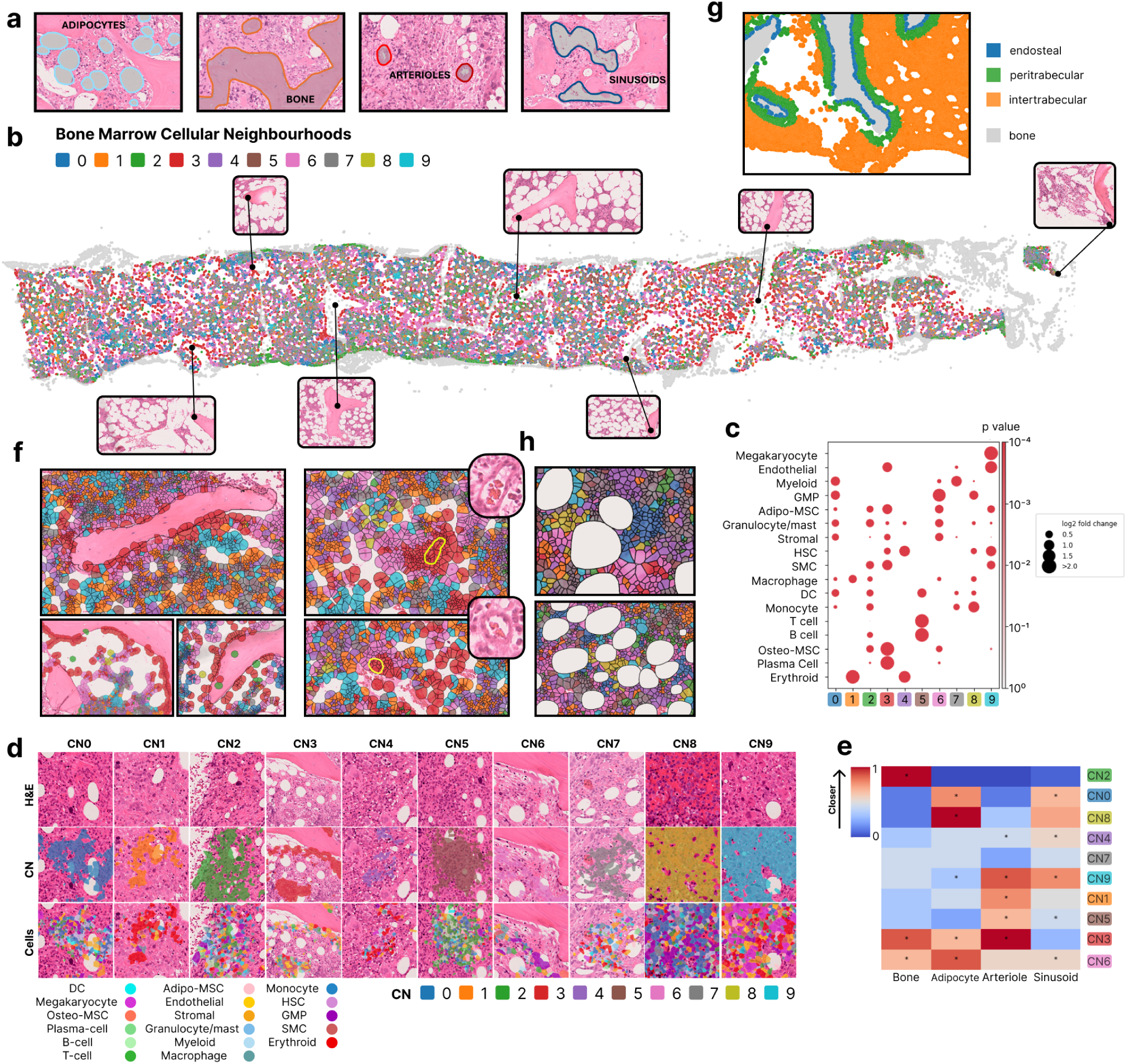
Mapping the normal human bone marrow using unsupervised cell neighbourhood analysis. **A:** Figure to show segmentation of adipocytes, bone, arteriolar and sinusoidal vascular structures. **B:** Bone marrow trephine (BMT) showing cell neighbourhoods (CNs) in a normal sample. CNs were identified using unsupervised clustering with *CellCharter* (k=3 hops, n = 30 samples) and are coloured by CN assignment (CN0-9). Inset images show the corresponding H&E images, highlighting peritrabecular regions. Cells highlighted in grey indicate those outside of regions of architectural preservation. **C:** Dot plot to show cell type enrichment in spatial cell neighbourhood (CN) clusters derived from *CellCharter* (1000 permutations, run across all samples/conditions, n = 30 samples). Dot size indicates fold enrichment. **D:** Panel to show H&E BM morphology (top), CN cell type assignment (middle) and the cell type composition of the same region (bottom) for each cell neighbourhood (CN0-9). **E:** Heatmap showing normalised rank proximity of CNs to key bone marrow topological structures (bone, adipocytes, arterioles, sinusoids). A rank towards 1 (red) infers proximity, while 0 (blue) indicates more distal (* denotes p<0.05, 100 permutations). **F:** Images to show: CNs (CN3, red; CN6, pink) proximal to bone (left), and the spatial location of arterioles within regions of CN3 (arterioles are denoted as a yellow polygon and inset shows the arteriole at higher power) (right). **G:** Figure to demonstrate BM architecture and topological regions. Those cells touching bone are highlighted in blue (‘endosteal’), cells within 1-300 pixels (aligned to the first ‘fat space’) of the bone surface are coloured green (‘peritrabecular’), and those within the ‘intertrabecular’ (or central) space are coloured orange. The bone is highlighted in grey. **H:** Images to show CN assignment of cells proximal to adipocytes. Cells are coloured by CN, and the white spaces are adipocytes.

To explore the distribution of CNs in relation to key topological BM features (bone, adipocytes, arterioles and sinusoids) ranking-based proximity analysis was performed^9,22^ (see Methods). We identified a previously unrecognised yet distinct endosteal/peritrabecular microenvironment, with CN2, CN3 and CN6 proximal to bone (Fig. 3D-G). CN2 was characterised by enrichment of MSCs (most prominently osteo-MSCs, but also adipo-MSCs and stromal cells) with a mixed immune cell population (B-cells, monocytes, macrophages, DCs); CN3 by co-enrichment of osteo-MSCs, SMCs, stromal cells, plasma cells, endothelial cells and adipo-MSCs; and CN6 by GMPs (Fig. 3C). CNs proximal to adipocytes included two endosteal/peritrabecular CNs (CN3 and CN6), likely reflecting the peritrabecular ‘fat space’ typically seen at the bone surface in the normal BM (Fig. 3H). In addition, CNs proximal to adipocytes were enriched for myeloid-lineage cells, including CN8 (GMP and monocyte co-enrichment); and CN0 (GMP, myeloid and DC enrichment) (Fig. 3C&E). The peri-arteriolar and sinusoidal microenvironments appeared distinct, with CN9 (megakaryocyte-enriched) most proximal to sinusoids and CN3 most proximal to arterioles (Fig. 3E). On visualisation, arterioles were consistently found within regions of CN3 (Fig 3F, Supplementary Fig. 9), however were not found to be proximal to bone with point pattern analysis (Supplementary Fig. 10).

To confirm and extend these observations, we performed global ranking-based proximity analysis of cell types to key architectural features (Methods). In keeping with the CN analysis, osteo-MSCs were most proximal to bone (lining the bone trabeculae), together with SMCs, GMPs, stromal cells, adipo-MSCs, and endothelial cells (Fig. 4A-D). The intertrabecular neighbourhood (defined as not proximal to bone) comprised monocytes, macrophages, erythroid cells and lymphocytes (Fig. 4A-D). Cell-cell proximity analysis demonstrated a ‘module’ of cellular proximity representing the intertrabecular cell neighbourhood, and another comprising endosteal niche-enriched cells (Fig. 4E). When we explored the cellular abundance within BM regions in accordance with conventional histological descriptions of the BM (as per Fig. 3G – across endosteal, peritrabecular and intertrabecular regions) our findings were concordant with proximity analysis (Supplementary Fig. 11). Erythroid cells showed greatest spatial autocorrelation, in keeping with a greater spatial clustering tendency (Supplementary Fig. 12). Megakaryocytes, contrary to conventional histological descriptions of the normal BM, also showed a tendency to cluster (Supplementary Fig. 12).

**Figure 4:**
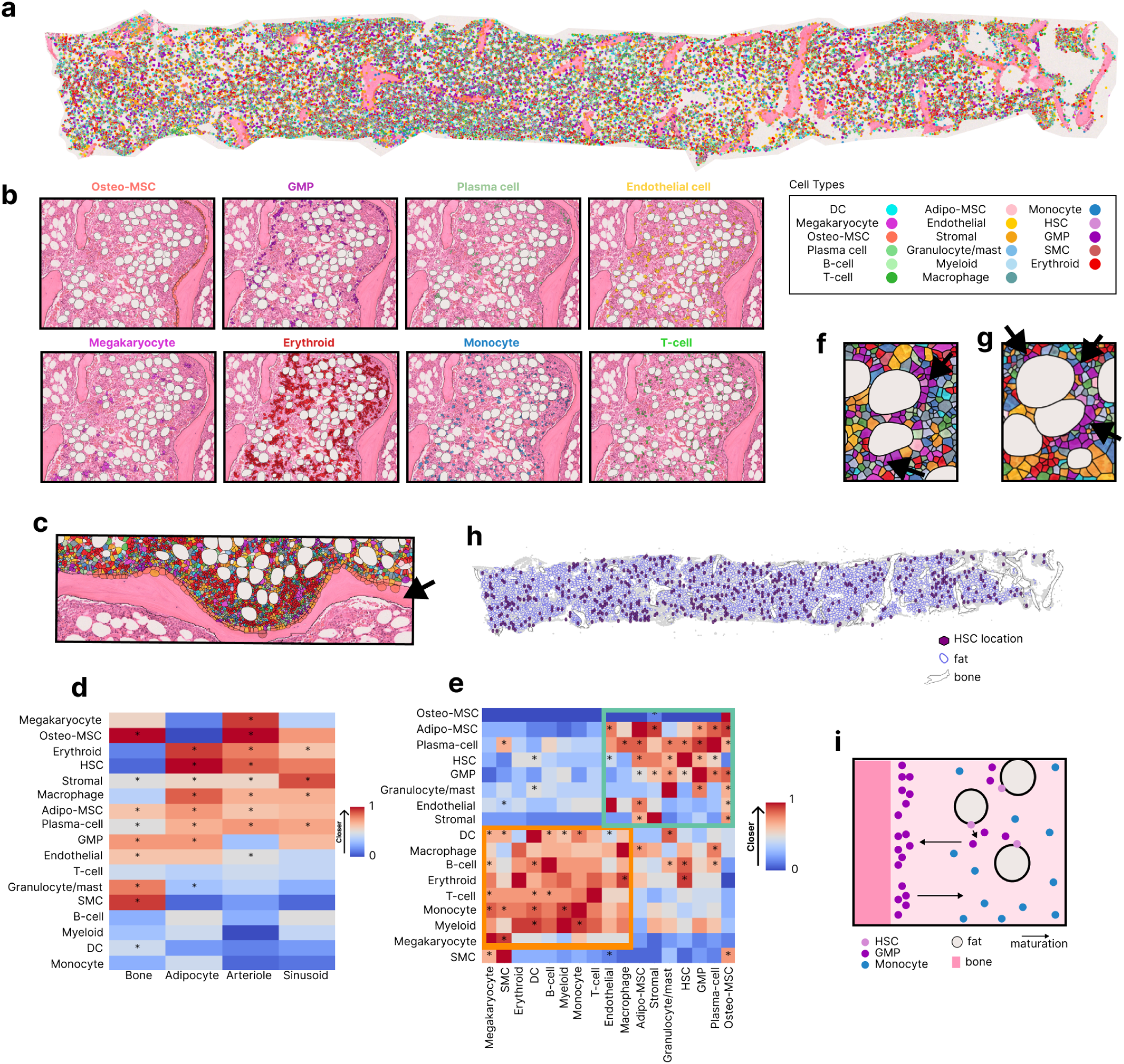
Characterising bone marrow topology with supervised spatial analysis. **A:** Normal BMT with cells coloured by type. **B:** Tile images to show cell types overlying the BMT H&E image in relation to the bone surface and distribution within the intertrabecular/central space. **C:** Image to show the BM microenvironment proximal to bone. Cells are coloured by type above the bone, and the H&E image is seen below. The osteo-MSCs (dark orange) and stromal cells (orange) are clearly seen lining the bony trabeculae. The arrow indicates the bone trabecula. **D:** Heatmaps showing normalised rank proximity of BM cell types in normal samples to key topological structures (bone, adipocytes, arterioles, sinusoids) and in **E:** other cell types. A rank towards 1 (red) infers proximity, while 0 (blue) indicates more distal (*denotes p<0.05, 100 permutations) (n = 5 normal samples, n = 509,333 cells). **F:** Tile images to show the peri-adipocytic location of HSCs (indicated by arrow). **G:** Tile images to show the peri-adipocytic location of GMPs (indicated by arrow). **H:** BMT image to show the distribution of HSCs and adipocytes. Purple hexagons (500 pixels in diameter) demonstrate HSC location. The blue circles bound the fat spaces. **I:** Schematic to demonstrate our inferred pattern of spatially-restricted myelopoiesis. HSCs (pink) are found proximal to adipocytes scattered across the BM. As they mature to GMPs (purple) they move to the bone surface. As they mature further along a mononuclear phagocytic trajectory (see monocytes, blue), they localise to the intertrabecular space.

We used this map to characterise the spatial distribution of haematopoiesis. Conventional descriptions of the HSC niche outline a spatially-restricted region local to a HSC, with constituent cells regulating HSC quiescence, differentiation and activation via stem cell factors^23^. Moreover, previous descriptions of the BM outline a predominantly endosteal HSC niche^24,25^. By contrast, our quantitative mapping approach revealed that haematopoietic progenitor populations (HSCs and GMPs) were proximal (as measured by point pattern analysis) to adipocytes (Fig. 4D, F&G), consistent with the role that adipocytes have in supporting haematopoiesis via cytokine-mediated HSC maintenance and activation^26^. Notably, we did not find a cell neighbourhood enriched solely for HSCs – rather, HSCs were distributed across several mixed cellular neighbourhoods, most prominently with erythroid cells (CN4) and megakaryocytes/endothelial cells (CN9) (Fig. 3C). The scattered spatial distribution of HSCs was confirmed on global review (Fig. 4H), and quantitated with Moran’s I, demonstrating low spatial autocorrelation and tendency towards random dispersion (Supplementary Fig. 12). GMPs were enriched in the endosteal/peritrabecular space, proximal to bone (Fig 4D & Supplementary Fig 11) and enriched within CN6, an endosteal CN (Fig. 3C&E). Mature mononuclear phagocytic populations (monocyte and macrophages) were depleted in the endosteal space and enriched within the peritrabecular and intertrabecular space (Supplementary Fig. 11). Together, this suggests that contrary to conventional descriptions of BM topology, the HSC compartment is peri-adipocytic, yet spatially dispersed without endosteal preponderance, with myelopoiesis characterised by migration of maturing progenitors to the bone surface, with localisation into the central, intertrabecular space with further maturation to mature myeloid populations (monocytes, macrophages) (Fig. 4I). The bone marrow vasculature regulates both haematopoiesis and osteogenesis via production of angiocrine factors^27,28^. This is further supported by our observation that HSCs were spatially proximal to arterioles (Fig. 4D), suggesting that progenitors have a proclivity for a well-oxygenated microenvironment.

Taken together, quantitative mapping of individual cell lineages, their respective cellular neighbourhoods and proximity analysis to histological tissue landmarks revealed highly conserved topological order within the normal human BM.

### Quantitative characterisation of bone marrow microenvironmental perturbation in MPN

Having demonstrated the utility and potential of ST augmented with AI-based feature detection to systematically analyse the normal BM ecosystem, we next sought to map the BM across the spectrum of MPN. We performed Xenium analysis of BMT FFPE material from patients with MPN, including ET (n = 8), PV (n = 3) and MF (n = 11) across 4,144,682 cells (Supplementary Fig. 1&2). Following expert haematopathological review, samples previously identified as pre-PMF were removed from downstream analysis due to lack of diagnostic concordance. We found no difference between the overall cellular composition of the BM in non-fibrotic MPN (PV and ET) and the normal BM. By contrast, MF was characterised by an increase in megakaryocytes and osteo-MSCs, and depletion of adipocytes (Extended Data Fig. 6).

To explore the spatial organisation of MPNs we performed ranking-based proximity analysis. While this revealed variation in the cellular microenvironments proximal to arterioles and sinusoids in MPN and the normal BM, notably the distinction between peritrabecular versus central microenvironmental niche spaces and periadipocytic localisation of HSCs observed in normal samples were conserved in MPN (Fig. 5, Supplementary Fig. 13). To interrogate microenvironmental heterogeneity in MPNs we used cell neighbourhood analysis. Strikingly, although globally we found no difference in CN abundance between conditions (Supplementary Fig. 14), MF (but not ET or PV) was characterised by expansion of a small number of large CN components (Fig. 6A-D, Extended Data Fig. 7) and greater inter-individual variation in the BM CN composition than in the normal BM, ET or PV (Fig. 6A, Extended Data Fig. 7). Notably, only some CNs were expanded in MF; namely, CN2 (stromal/mixed immune co-enriched), CN9 (megakaryocytes/endothelial) and CN0 (myeloid, GMP, stromal, DCs) (Fig. 6E, Supplementary Fig. 15). While there was no evidence of overall CN component expansion in ET or PV, CN2 and CN9 also represented the largest CN type components in ET (Fig. 6E), but not in PV or normal samples (Supplementary Fig. 16), suggesting a possible shared axis of perturbation in ET and MF. Taken together, we hypothesise that this pattern of microenvironmental remodelling and CN expansion might reflect regions of clonal expansion. Using CN rank proximity analysis, we also found that the endosteal/peritrabecular and central niche spaces were preserved, with CN3 and CN6 consistently proximal to bone in MPN and normal samples (Fig. 6F&G), confirming our cell-wise rank proximity analysis (Figure 5) and suggesting that this pattern of CN expansion occurs within the context of a highly preserved endosteal/peritrabecular niche.

**Figure 5:**
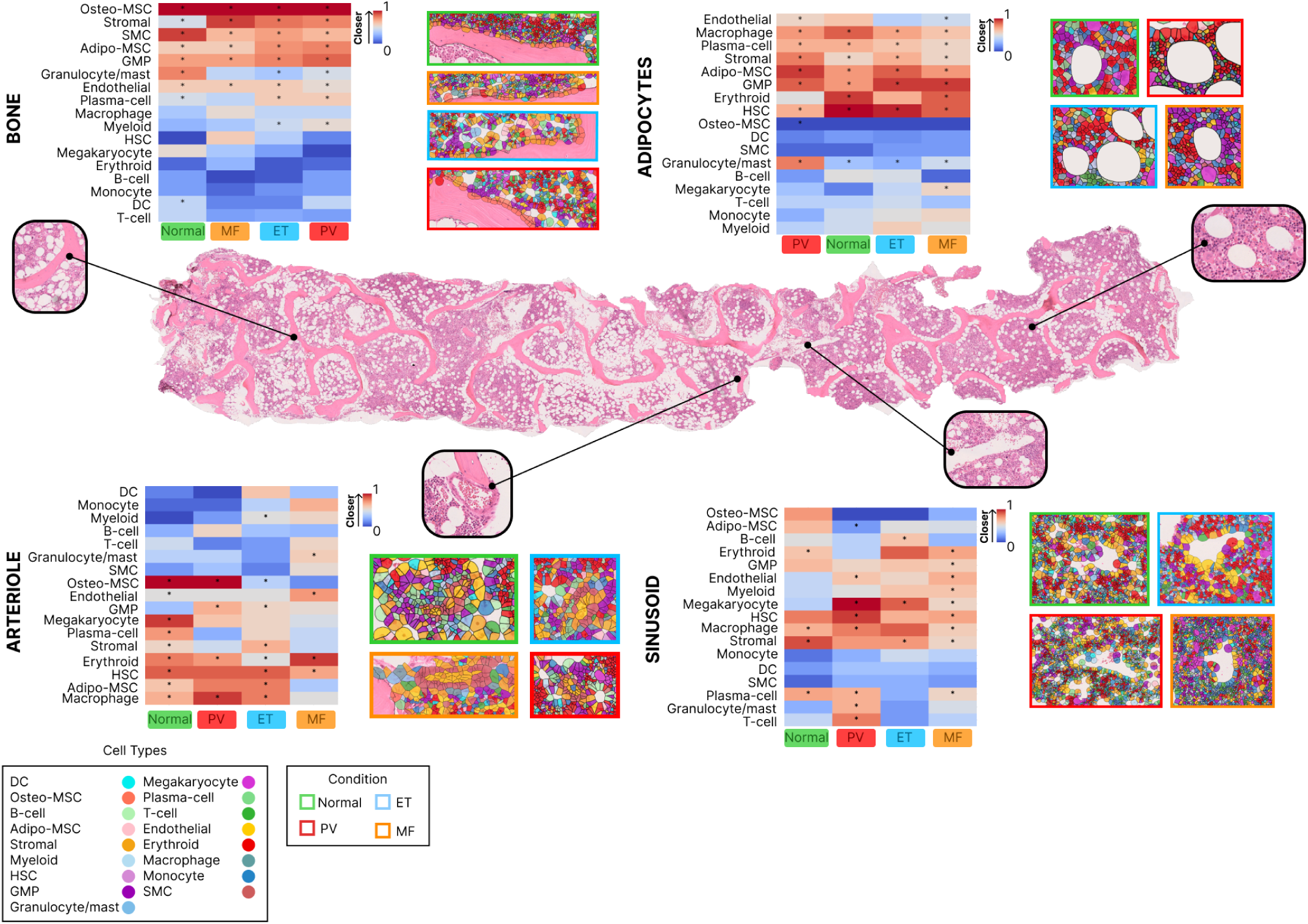
Quantitatively mapping the bone marrow landscape in myeloproliferative neoplasms. Bone marrow trephine H&E image (PV) with inset images to show key architectural features (left to right: bone, arterioles, sinusoids, adipocytes). Heatmaps are hierarchically clustered and show normalised rank proximity of BM cell types to key topological structures labelled to the left of each plot (bone, adipocytes, arterioles, sinusoids) across MPN subtypes (PV, ET, MF) and normal samples. A rank towards red infers proximity, while blue indicates more distal location (*denotes p <0.05 across 100 permutation runs). To the right of each heatmap the tile images show representative corresponding regions with cells coloured by type. The border of each tile image indicates the condition (normal = green, ET = blue, PV = red, MF = orange).

**Figure 6:**
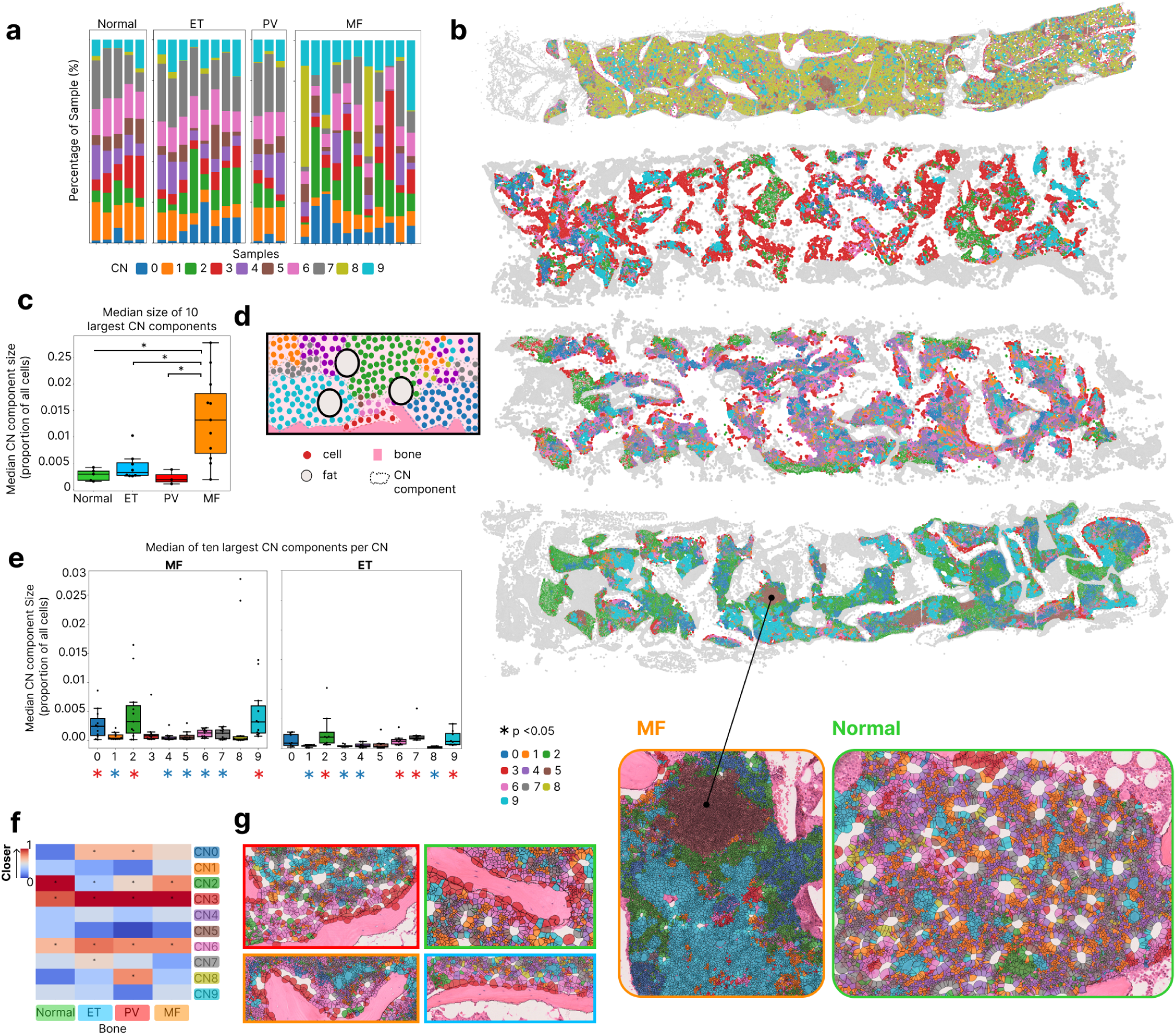
Characterising cell neighbourhoods in myeloproliferative neoplasms. **A:** Stacked bar plot to show the proportion of cells per sample assigned to each cell neighbourhood (CN 0-9) across MPN and normal samples (n = 27 samples, n = 4,654,015 cells). **B:** BMTs from four patients with MF to show the distribution, size and heterogeneity of CN distribution. Each cell is coloured by CN assignment (0-9). The inset images at the bottom demonstrate cells coloured by CN in MF (left) versus a normal sample (right). There are larger, expanded CN components in MF compared to normal samples. **C:** Boxplot to show the median size of the ten largest cell neighbourhood components as a proportion of all cells across conditions (Mann Whitney U test, *p<0.05). **D:** Schematic to demonstrate a cell neighbourhood (CN) component. Cells (dots) are coloured by cell neighbourhood assignment, discrete cell neighbourhood components are indicated via dashed lines. CN components are regions comprising at least five cells of the same CN connected in the spatial graph. Bone is indicated in pink and white circles represent adipocytes. **E:** Boxplot to show the median size of the ten largest cell neighbourhood components as a proportion of all cells by condition (Wilcoxon signed-rank test, *indicates p<0.05, red star (*) indicates that the component size of that CN is significantly increased compared to at least one other CN, and blue star (*) indicates significantly reduced compared to at least one other CN.) **F:** Heatmap to show normalized rank proximity of CNs to bone across conditions (normal, PV, ET, MF). A rank towards red infers proximity; blue infers a more distal location (*denotes p <0.05, permutation test with 100 runs). **G:** Images to show BM cells coloured by CN in relation to bone trabeculae. Cells assigned to CN3 (red) and CN6 (pink) are clearly seen lining the bony trabeculae across normal (top right, green border), ET (bottom right, orange border), PV (top left, red border) and MF (bottom left, orange border).

### Integration of ST data with AI-based fibrosis detection identifies microenvironmental features associated with bone marrow fibrosis

A defining feature of MF employed both diagnostically and prognostically is the presence of BM fibrosis, with fibrotic progression in MPN associated with a worsening clinical outcome^29^. To identify microenvironmental correlates of fibrosis in the human BM, we used our reticulin-free AI algorithm which outputs a quantitative metric of fibrosis, or ‘Continuous Index of Fibrosis’ (CIF) score^30,31^ (Fig. 7A, Extended Data Fig.8, Supplementary Fig. 17A). Through integrating ST data with CIF quantitation - *on the same tissue section* (Fig. 7A) - we identified cell-wise enrichment associated with regions of fibrosis (Fig. 7B, Extended Data Fig. 8B). There was a positive correlation between CIF and monocyte and macrophage abundance in MF and ET, but not in PV (Fig. 7B, Extended Data Fig. 8B). Although MSCs induce a pro-inflammatory environment, with production of factors such as TGF-β driving the development of fibrosis in MF^32,33^, unexpectedly, stromal cells and osteo-MSCs were depleted with increasing fibrosis (Fig. 7B). To explore this further, we then assigned the CIF score to the corresponding WHO fibrosis grade^34,35^, a framework used to categorise the severity of reticulin fibrosis in clinical pactice (Fig. 7A, Supplementary Fig. 17B). In MF, even in regions with no significant increase in fibrosis (WHO grade MF-0), we found enrichment of stromal cells, osteo-MSCs and megakaryocytes; and depletion of B-cells and T-cells when compared to normal samples (Extended Data Fig. 8C). This suggests that in MF the non-fibrotic microenvironment is characterised by a lymphocyte-deplete but stromal-enriched state, with stromal depletion and immune (both lymphocyte and mononuclear phagocytic (MNP) cell) enrichment associated with increasing fibrosis. By implication, this also suggests that the contributory role of stromal cells in mediating marrow fibrosis is either via local expansion of a subset not captured in our dataset, or is spatially or temporally distant, with stromal cells mediating fibrotic deposition with subsequent local ‘drop-out’ as more severe fibrosis accumulates.

**Figure 7:**
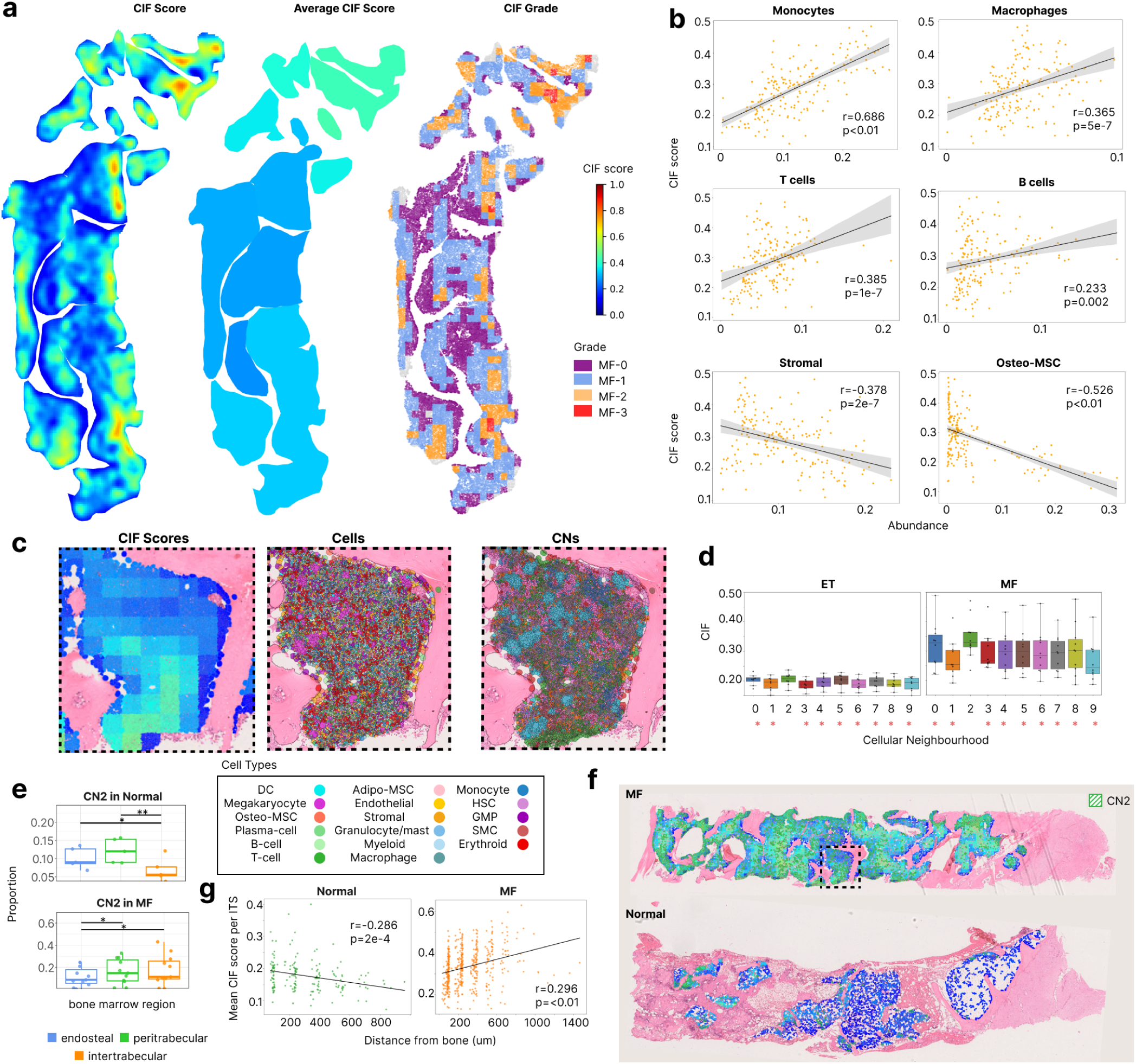
Identifying microenvironmental correlates of fibrosis. **A:** BMT heatmaps for an MF sample to show: (Left BMT) Continuous Indexing of Fibrosis (CIF) score. A higher CIF score (fibrosis) is indicated by yellow/orange/red, lower by blue and green (see scale right); (Middle BMT) Mean CIF score per intertrabecular space (ITS) (scale right); (Right BMT) WHO fibrosis grade (WHO MF-0-3) per image tile (512x512 pixels) (for colour key see right). **B:** Scatter plots to show the proportion of cell types per ITS (mean score across 512x512 pixel tiles per ITS) in MF samples (n=11, Pearson’s correlation). **C:** Images to show integration of: (left) CIF heatmap; (middle) cell type; (right) cell neighbourhoods (CNs) within an intertrabecular space (ITS). **D:** Boxplots to show the corresponding median CIF score of all cells within a CN per sample in: (left) ET and (right) MF. *Denotes p < 0.05 with a red star indicating that the median CIF was significantly different from CN2 (linear mixed effects model with Benjamini-Hochberg correction). **E:** Boxplot to show differential abundance (proportion of all cells within specified region) assigned to CN2 (stromal and immune enriched CN) across BM regions (endosteal, peritrabecular, intertrabecular) in: (top) normal (n = 5) and (bottom) MF (n = 11) (negative-binomial generalised linear model *(edgeR)* with Benjamini-Hochberg correction (*<0.05, **p<0.01). **F:** BMT heatmaps to show CIF score (see scale as per 7A) and the distribution of CN2 (green). The box highlighted with a dashed line corresponds to the image seen at higher power in 7C. CN2 is seen in the intertrabecular regions in MF. **G**: Scatter plot to show the mean CIF score per ITS plotted by distance from bone (μm) in (left) normal and (right) MF samples (Pearsons’s correlation).

**Figure 8:**
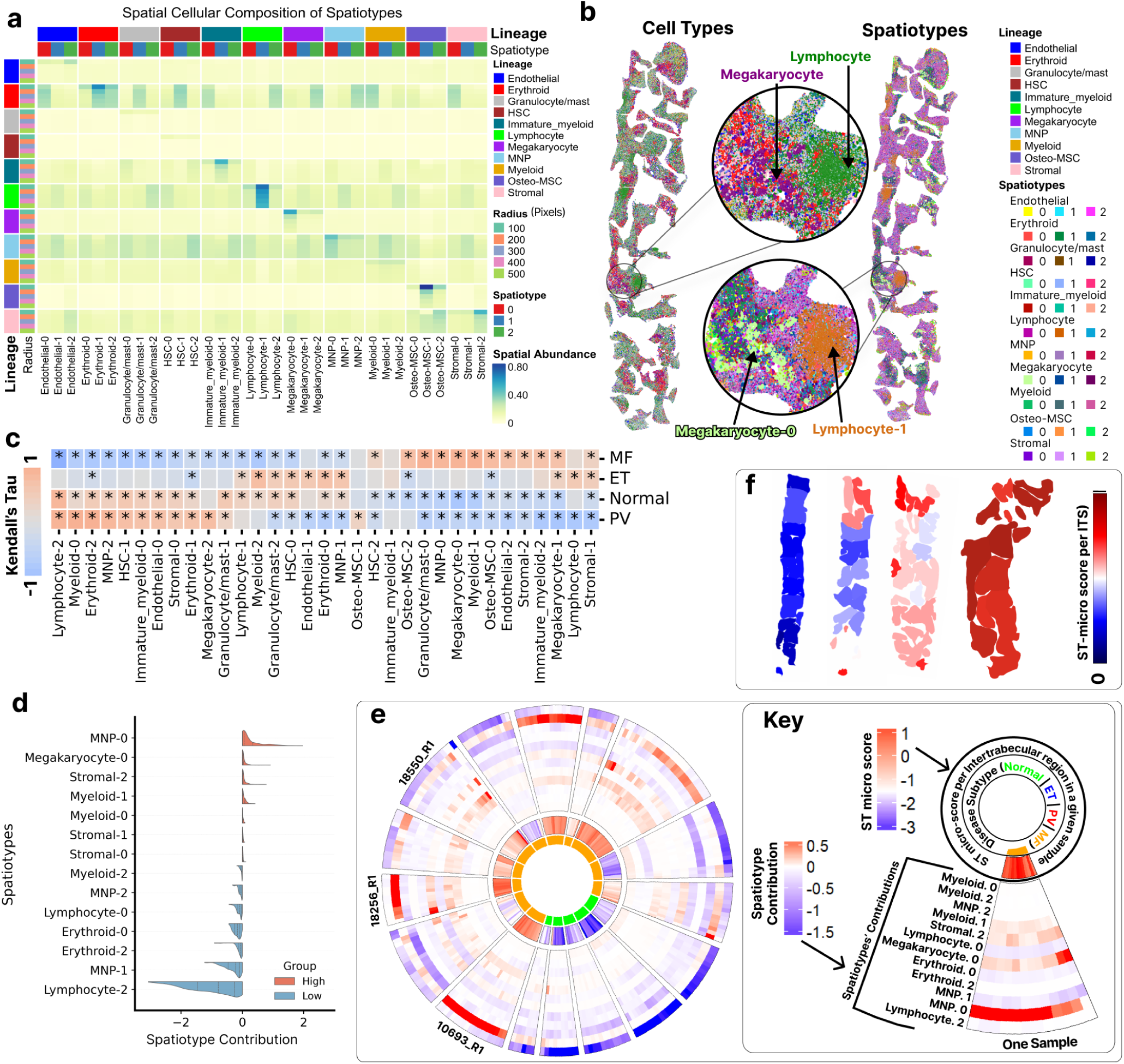
Identifying bone marrow spatial microenvironmental signatures in health and disease. **A:** Heatmap of cellular composition across spatiotypes. Cells are clustered by differential cellular abundance within radial windows (0-500 pixels). Columns are grouped by cell type, with three spatiotypes (0–2) per type (red=0, blue=1, green=2). Row-wise outer bars indicate cell type (lineage) and inner bars capture radius (100-500 pixels). Heatmap colours indicate relative abundance (blue=high, yellow=low). **B:** BMT images demonstrating cell type versus spatiotype assignment. Left BMT: cells coloured by cell type; Right BMT: cells coloured by spatiotype assignment. Inset images show the cell type and spatiotype annotation in the same region (see key). **C:** Hierarchically clustered heatmap of Kendall’s tau correlations between ITS-region-level spatiotype abundance (x-axis) and disease subtypes (y-axis). Red/blue indicates positive/negative correlations; asterisks (*) denote significant correlations (Benjamini–Hochberg FDR-adjusted *p* ≪ 0.05). **D:** Association between spatiotypes and ST-micro score. Violin plots summarise the average effect (*α*) of each spatiotype on the ST-micro score across test samples over 20 bootstrap runs. Red (mean *α* > 0) indicates contributions towards an MF-like microenvironment and blue (mean *α* < 0) towards a normal-like microenvironment; distance from the centre line (*α*=0) indicates effect size. **E:** Circle plot illustrating intra- and inter-patient heterogeneity. Each radial segment represents a patient and is subdivided into intertrabecular spaces (ITS). The inner ring shows disease subtype (normal or MF), the middle ring ITS-level ST-micro score, and the outer ring represents a heatmap of the 12 spatiotypes with the strongest positive or negative contributions to the ST-micro score. Colour gradients indicate contribution, with red reflecting MF-like and blue normal-like microenvironmental signatures. **F:** BMT heatmaps showing ST-micro scores across ITS regions. Dark red (+1) and dark blue (0) indicate MF-like and normal-like microenvironments, respectively. Scores are derived from an ML model trained to distinguish MF from normal based on spatiotype patterns.

To further interrogate cellular microenvironmental features associated with fibrosis, we explored the association between CNs and CIF (Fig. 7C). CN2, characterised by stromal (most significantly osteo-MSCs) and immune (monocytes, macrophages, DCs, granulocyte/mast cells, B-cells) co-enrichment (Fig. 3C) was associated with increased fibrosis (highest CIF score) in MF and ET (Fig. 7D). Notably, while CN2 was depleted in the intertrabecular space in normal samples, it was enriched in MF (Fig. 7E&F). This suggests that in MF there is a distinct pattern of stromal and immune co-enrichment that expands from the bone surface. This is in keeping with recent descriptions by Banjanin et al.^36^, who describe a pro-fibrotic osteolineage MSC population expanding from the bone surface in MF. Corroboratively, we found that in MF there was increasing fibrosis with distance from bone (Fig. 7G).

### Identifying bone marrow microenvironmental signatures to dissect sample and cohort-level heterogeneity

We then sought to more fully exploit the potential offered by ST for unsupervised quantitative interrogation of tissue microenvironments for the purpose of: 1) improving understanding of disease biology; 2) providing quantitative ways of capturing both cohort and sample level variation; and, 3) supporting hypothesis generation. We used our previously developed tissue-based ‘band descriptor’^37^ to identify spatial patterns of microenvironmental enrichment. In brief, using broader cell-level annotations (see Supplementary Fig. 20), for each cell we identified the proportion of cell types within a capture area of increasing diameter (see Methods, Extended Data Fig. 9A). We then performed k-means clustering according to cell proportions to identify cell groups defined by distinct local microenvironments. For example, we identified three megakaryocyte groups: megakaryocyte-0, characterised by local enrichment of megakaryocytes (i.e. megakaryocyte clusters); megakaryocyte-1, by local enrichment of MNPs; and megakaryocyte-2, by local enrichment of erythroid cells (Fig. 8A&B, Extended Data Fig. 9B). We termed these cell groups (and their associated microenvironmental patterns) ‘spatiotypes’. This analysis was performed across all cell types, generating a composite microenvironmental signature of the normal BM and BM in MPN (Fig. 8C).

This ST-derived data-driven approach revealed novel spatial microevironmental patterns in MPN. For example, while entirely in keeping with conventional pathological descriptions MF was enriched for megakaryocyte-0, (i.e. a megakaryocyte clustering signature) (Fig. 8A&C), it was also characterised by enrichment of novel spatial sigantures including spatiotypes associated with MNP cell enrichment (‘MNP-0’, ‘myeloid-1’ and ‘granulocyte/mast-0’) and stromal cells (‘stromal-2’ and ‘osteo-MSC-2’) (Fig. 8A&C). Lymphocyte-2 (lymphocyte-erythroid enrichment) was enriched in the normal BM, but depleted in MF (Fig. 8A&C). Notably, by comparing spatial patterns across MPN subtypes we demonstrated no concordance between spatiotypes enriched in MF and the normal BM, indicating a high degree of multi-lineage spatial perturbation in MF (Fig. 8C). In contrast, both ET and PV showed some concordance (shared spatiotypes) with the normal BM, with PV demonstrating greater commonality. However, we found that the spatiotypes in ET and PV that were divergent from the normal BM (not enriched in the normal BM) were discrete, indicating divergent patterns of microenvironmental deviation between these disease subtypes.

Finally, we then sought to employ these novel spatiotypes to more robustly interrogate both inter-and intra-individual microenvironmental BM heterogeneity. To do this, we developed a ranking-based multiple instance machine learning (ML) model trained on ST data to derive a single ST-microenvironment (‘ST-micro’) score indicating the extent to which the microenvironment (composite spatiotype signature) was akin to the normal BM, or to MF (see Methods). At a cohort level, the spatiotype most contributory to an MF-like ST-micro score was MNP-0 (MNP-MNP enrichment) (Fig. 8D). Megakaryocyte-0 (megakaryocyte-megakaryocyte enrichment), a known pathological feature of MF, was also highly contributory. Lymphocyte-2 (lymphocyte-erythroid enrichment) was the most prominent contributor to a normal ST-micro score (Fig. 8D). This approach allowed us to objectively capture inter-sample variability across the cohort. Importantly, we found that there was variation in the spatiotypes underpinning an MF-like ST-micro signature/score (Fig. 8E, Supplementary Fig. 21A). For example, the ‘MF-like’ ST-micro score for samples 10693_R1 and 18256_R1 was driven predominantly by MNP-0 (MNP-MNP co-enrichment), while the ST-micro score (again, ‘MF-like’) for sample 18550_R1 was underpinned by spatiotypes including MNP-0, Megakaryocyte-0, stromal-2 and myeloid-1. We then used this approach to explore intra-sample heterogeneity. In some samples, the ST-micro score was consistent across the trephine (‘MF-like’ towards +1; or akin to normal, towards 0), while some BMTs demonstrated marked intra-sample variation (Fig. 8F, Extended Data Fig. 9C). Critically, the output of this approach offers the benefit of explainability – one can identify the spatiotype driving a particular MF-like signature and characterise how this varies across a BMT sample (Extended Data Fig. 10). Notably, although there was correlation between overall ST-micro score and CIF (Supplementary Fig. 21B), there was discordance in some samples (e.g. high MF-like ST-micro scores and low fibrosis scores) suggesting that cell-cell microenvironments alone do not determine pathological marrow fibrosis in MPN patients.

Taken together, these findings indicate the clear potential of unsupervised microenvironmental quantitative approaches (leveraging ST-based approaches, ML and quantitative feature analysis) for the detailed characterisation of disease-associated perturbation in the human BM and identification of novel spatial signatures of both health and disease.

## DISCUSSION

Here, we describe an integrated workflow for the analysis of human BM ST data derived from routine diagnostic FFPE material, augmented by the application of AI-based image-analysis. In contrast to recent ST-based BM studies^7,8,38^, rigorous expert-led spatial morphology-driven QC removed areas of haemorrhage, crush and folded tissue from downstream analysis. We apply our recently developed AI-powered image-based algorithms to support BM feature detection, demonstrating how this can be integrated with ST within a single framework, both enabling characterisation of features not captured on ST alone and high-confidence topological mapping in architecturally preserved tissue regions. The result is a multi-layered, data-driven approach designed to quantitatively map the BM in MPN and fully leverage the unparalleled tissue-based insights offered by ST.

Our approach redefines our understanding of the normal BM niche. In contrast to a recent IF-based BM mapping study^9^ in which bone detachment compromised tissue spatial integrity, we successfully applied unsupervised cell neighbourhood analysis to the entire marrow space, including endosteal and peritrabecular niches. We find the human BM to be characterised by distinct endosteal and intertrabecular microenvironments. Peritrabecular regions are enriched for stromal cells and GMPs, while intertrabecular spaces are enriched for erythroid, mature myeloid groups (monocytes/macrophages), lymphocytes and megakaryocytes. This supports work by Lin et al.^38^, who used *Visium* to identify an osteogenic transcriptomic signature close to bone, and an intertrabecular immune-enriched signature. We find that this distinction is remarkably conserved across MPN subtypes, even in MF which is characterised by spatial perturbation. Likely reflective of the key role that adipocytes play in supporting haematopoiesis via production of cytokines including CXCL12, CSF3 and IL-8^39,40^ we demonstrate that the HSC niche is peri-adipocytic. However, given the scattered, dispersed distribution of HSCs, we consider the HSC niche to be best regarded a functional rather than spatially-restricted niche. Our data also supports and expands recent descriptions of spatially-restricted myelopoiesis^9^. In keeping with Bandyopadhyay et al.^9^ our data suggests that the myeloid differentiation trajectory is characterised by localisation of GMPs to the bone surface with migration to the central/intertrabecular space where mature MNP cell groups reside. However, unlike this group, we find that only macrophages are proximal to sinusoids, with no evidence of further localisation of other mature myeloid populations in sinusoidal or other vascular spaces.

Our study provides novel insights into MPN biology. Previous spatially-resolved descriptions of the BM in MPN have predominantly utilised murine models^36,41^. However, the murine BM is distinct in both cellular composition and topology^42^, and models of MPN do not faithfully recapitulate human disease trajectories. Our data provide the first whole-section quantitative description of the human BM in MPN at single cell resolution across a large cohort. We find that, in contrast to ET and PV, MF is characterised by a small number of large, expanded cell neighbourhoods. Such regional loss of local ecological diversity might, in part, suggest aberrancy in cell-cell localisation and cross-talk in MF. We hypothesise that this pattern of CN expansion might represent localised expansion of a mutation-bearing clone or a distinct clonal architecture.

By integrating ST data with AI-based image analysis, for the first time we can directly identify microenvironmental features associated with fibrosis across MPNs. Monocytes and myeloid-lineage cells are implicated in the development of BM fibrosis^14,43–45^ and we present corroborative spatial mapping, with fibrosis associated with enrichment of monocytes and macrophages. While existing studies describe the key role of MSCs in driving the development of BM fibrosis^13,36,44^, unexpectedly we found stromal cells to be depleted in regions of more severe fibrosis, suggesting that the contributory effect of MSCs may be most pertinent in the early stages of fibrosis (where collagen deposition is yet minimal), or is spatially distal from regions of fibrosis deposition. We identify a spatial neighbourhood (CN2), enriched for stromal cells (particularly osteo-MSCs) and a mixed immune population (DCs, monocytes and B-cells and granulocyte/mast cells) associated with fibrosis, which extends from the bone surface into the central space in MF. This is in keeping with recent work by Banjanin et al.^36^, who demonstrated that the development of BM fibrosis is driven by the expansion of an osteolineage stromal population from the bone surface in both the murine and human BM in MF. Our unsupervised analysis is supportive of a dual role for both osteogenic stromal and mixed immune populations in underpinning the development of fibrosis, and is corroborative of a spatially-restricted model of fibrotic development in MF.

We also present computational tools for capturing tissue microenvironmental heterogeneity. By employing our previously described approach for the identification of BM spatial microenvironmental signatures or ‘spatiotypes’^37^, we adopt a novel approach for leveraging ST-data, identifying novel spatial microenvironmental signatures that define the normal BM (erythroid-lymphocyte co-enrichment), and BM in MPN (local MNP enrichment). This orthogonal analytic approach validates our observations that mature MNP cell groups (monocyte and macrophages) are enriched in fibrosis. Moreover, this approach highlights the extent to which the marrow in health and MF are microenvironmentally distinct. Notably, we find that PV is more akin to the normal BM than ET, providing quantitative corroboration of conventional pathological descriptions of PV as a panmyelotic state with ‘balanced’ hyperplasia of all three haematopoietic lineages^46^ – we demonstrate that this likely occurs within the context of relative spatial preservation. The lack of shared spatiotypes between ET and PV was also striking, and demonstrates that these conditions are characterised by distinct patterns of microenvironmental deviation.

Finally, we use these spatiotype signatures to develop a ranking-based multiple instance learning (MIL) algorithm that quantitatively captures BM microenvironmental spatial heterogeneity at both a sample and cohort level. MF demonstrates marked intra- and inter-individual microenvironmental heterogeneity, suggesting that disease development and progression is a spatially and/or temporally highly heterogeneous and characterised by disparate disease trajectories and differential patterns of microenvironmental support and activation. It may be this previously unseen BM microenvironmental heterogeneity in MF patients contributes to prognosis discrepancy, variable clinical trajectories, and differential treatment response.

Our work demonstrates the power of a highly data-driven approach to ST analysis, with rigorous morphology-based QC, integration of computational AI-based tools for feature detection, both supervised and unsupervised spatial analysis providing a truly rich, ‘multi-lens’ perspective of human BM topology. Together, our observations redefine our understanding of human BM topology, provides a generalisable toolkit for spatial biomarker discovery in human disease.

## METHODS

### Samples and cohort

Human archival formalin-fixed paraffin-embedded (FFPE) bone marrow trephines (BMTs) were identified from the clinical archive at the John Radcliffe Hospital, Oxford University Hospitals NHS Foundation Trust. These BMTs had been fixed in 10% neutral buffered formalin followed by EDTA decalcification for 48 hours and paraffin embedding according to local standard clinical protocols. BMTs underwent pathological review to confirm adequacy (at least five intertrabecular spaces) and diagnosis. All samples were part of the INForMeD study (IRAS ID: 199833; REC reference: 16/LO/1376; PI AJ Mead). Myelofibrosis (‘MF’) samples were defined as those in fibrotic phase (WHO MF->2) according to standard diagnostic criteria.

### Sample preparation for Xenium

Samples were prepared as we have previously described^15^ and as per manufacturer’s standard published protocols. In brief, a H&E image of each BMT was reviewed to confirm sample adequacy/suitability for ST analysis. The region of interest/BMT in the FFPE block was gently scored with a scalpel. Prior to sectioning the environment was treated with RNaseZap. The FFPE block was placed on ice. A microtome was used to cut thin sections of tissue, initially discarding the first few sections. A full-face section was taken and manoeuvred onto the surface of a water bath (at 42°C) with tweezers. Excess paraffin wax was removed using tweezers, carefully avoiding the tissue region of interest. The process of excess paraffin removal was facilitated by scoring the block prior to cutting. This process was repeated until all required tissue for placement on a single Xenium slide was cut. The BMT sections were then manoeuvred onto the surface of the Xenium slide within the specified capture area (12 x 24 mm). The tissue was then dried according to standard protocol, and then stored in a desiccator prior to further processing.

### Xenium in-situ analysis

Following sectioning onto the 10x Xenium slides, deparaffinisation and permeabilisation was carried out as per manufacturer’s protocols. We used the Xenium multi-tissue and cancer probe panel comprising 377 probe markers. Following probe hybridisation, samples were run on a Xenium Analyzer as per manufacturer’s standard protocols.

### H&E staining post ST-analysis

Following data generation, samples (on the Xenium slides) were washed with MilliQ water and then stained with haematoxylin and eosin (H&E) using an automated wet tissue staining protocol. H&E-stained histological WSI images were generated using a Hamamatsu NanoZoomer S210 digital slide scanner.

## DATA PRE-PROCESSING AND QUALITY CONTROL

### Quality-control pre-processing and cell segmentation

Xenium explorer outputs a matrix of transcripts per cell, removing low quality transcripts (Q-score <20). To assign transcripts to the appropriate cell, we used the 10x Xenium In-situ Onboard Analysis software v1. This process segments nuclei according to the DAPI nuclear signal using a deep neural network approach which expands the boundaries of segmented nuclei by 15 μm, or until they encounter another cell boundary.

### Image registration

The H&E image (performed on the same section) was registered to the ST data (DAPI image) using the registration facility in Xenium Explorer v1. Corresponding histological landmarks were manually selected in H&E and DAPI images and a transformation matrix generated to align the H&E image to the DAPI. The registered images were reviewed using the Xenium Explorer v1 package to ensure accurate image registration.

### H&E-based annotation for QC and filtering

To remove areas of tissue unsuitable for downstream analysis (containing tissue or bone folding/detachment or crushed tissue) annotation of the H&E material was performed by a haematopathologist using the annotation tool in Xenium explorer (Extended Data Fig. 2). In brief, we first annotated and removed areas unsuitable for downstream analysis in which lack of cellular preservation might compromise cell-type assignment. This included areas of crushed tissue, haemorrhage, or tissue folding (Extended Data Fig. 1), termed ‘cytomorphologically unpreserved’. Cells within these areas were removed from any further downstream analysis. Across all 30 samples, 464,793 cells were within regions of poor cellular preservation and not taken forward for any further downstream analysis (e.g. cell typing). This included 87,268 cells with a low transcript number (<4 transcripts/cell), however following this annotation step, we also removed all remaining cells with a low transcript number (<4 transcripts/cell, 180,784 cells). Together, a total of 672,360 cells were removed from downstream analysis (cell typing or spatial analyses).

We also found that some areas of tissue demonstrated good cellular cytomorphological preservation, but poor architectural/topological preservation. While these areas were suitable for cell typing and BM composition quantitation, they were unsuitable for downstream spatial analysis as the tissue in these areas did not reliably represent tissue architecture in vivo. We performed a second round of haematopathologist-led annotation of the H&E image, identifying architecturally preserved regions. These annotations broadly aligned to an intertrabecular space (ITS) – a region between bony trabeculae (Extended Data Fig. 2). Of the 5,285,236 cells retained after removal of those in regions of poor cytomorphological preservation, 4,422,020 cells (84%) lay within spatially preserved regions.

### AI-enhanced megakaryocyte segmentation

We noticed that large, polylobated cells such as megakaryocytes were segmented into numerous cells (or ‘multipolygons’) by the Xenium segmentation tool v1 (Extended Data Fig. 3C). We therefore applied our previously developed H&E-based AI-based megakaryocyte detection and segmentation algorithm to the corresponding H&E image as previously described^16^. The coordinates of segmented megakaryocytes were then registered to the DAPI image using the transformation matrix computed for the H&E-to-DAPI image registration.

To consolidate Xenium cell segmentation with H&E-based megakaryocyte masks, we first brought Xenium cell and nucleus boundaries and H&E-derived megakaryocyte segmentations into a common pixel space and coordinate frame. We then used the GeoPandas overlay function (how = “difference”) to subtract H&E-derived megakaryocyte polygons from the Xenium cell boundaries. This operation both identified cell boundaries that intersected megakaryocytes and returned only the residual parts of those boundaries, while leaving non-overlapping cell boundaries unchanged. The majority of megakaryocyte cell boundaries remained unchanged and were placed back into the original segmentation. For the cells that were altered by the megakaryocyte overlay, multipolygons were exploded into separate cells. Those cells with boundaries that contained a nucleus centroid were placed back into the original segmentation. Cells with boundaries that did not contain a nucleus centroid were divided into two groups according to a size threshold. The threshold was set at half the median area of all cells in the original Xenium segmentation. Cells below the threshold were merged into the nearest non-megakaryocyte cell. Cells above the threshold without a nucleus were labelled as ‘artefacts’ and were removed from downstream analysis. Finally, we computed the transcript per cell based on the updated cell boundaries using the Sopa package^47^. AI-enhanced segmentation identified 31,583 cells and we removed 27,449 (0.005%) ‘artefacts’ (polygons without a nucleus) generated after AI-detected megakaryocyte segmentation and insertion.

### Cell-type annotation

We observed little variation across six Xenium runs (batches of x2 slides) (Extended Data Fig. 3F), however, to mitigate for any batch effect across the six runs the gene expression matrix was normalised using *sctransform* from the *Seurat v5*^48^ package. We used *Harmony*^49^ for batch correction using 50 principal components.

We utilised both gene expression data and spatial information for cell-type assignment. First, we identified HSCs as those cells co-expressing *CD34* and *KIT* (total of 27,059 cells). These cells were removed from downstream unsupervised clustering. We performed dimensionality reduction, clustering and visualisation using *Seurat v5*. We constructed the KNN graph with *FindNeighbours* from the first 20 principal components from *Harmony*. We explored cluster stability across resolutions of 0.2, 0.4, 0.6 and 1. We performed Louvain clustering with *FindClusters* at a resolution of 1, returning 50 cell clusters. We then merged the HSC group back into the dataset and performed differential expression analysis with *FindAllMarkers* (log2FC > 1, adj.p-val < 0.05 and min.pct1 > 0.25). We used Uniform Manifold Approximation and Projection (UMAP) to visualise cell clusters.

To confirm cluster assignment, we also annotated our clusters using *SingleR*^50^ using two reference bone marrow scRNA-seq datasets; the HuBMAP Azimuth Bone Marrow reference dataset^17^ and a stromal-enriched scRNA-sequence dataset (Bandyopadhyay et al^9^). We subsetted each reference dataset to the common genes (375 and 365 respectively) and used the second annotation level in each to transfer labels to each cell. Informed by both supervised annotation (with *SingleR*) and the results from differential expression analysis, we then merged clusters of the same cell-type annotation.

To identify stromal sub-populations we performed supervised annotation with the Bandyopadhyay et al. scRNA-seq dataset which is enriched for stromal cells. We used *SingleR* to transfer labels from this stromal-enriched bone marrow reference scRNA-seq dataset to cells that had been assigned as stromal as per lineage marker gene expression following unsupervised clustering.

Any cell not annotated as either osteo-MSC, osteoblast, adipo-MSC, vascular smooth muscle cell (VSMC) or smooth muscle cell (SMC) by *SingleR* was labelled ‘stromal’. For the remaining cells we transferred the *SingleR* annotations, merging osteo-MSC and osteoblast into an ‘osteo-MSC’ group and VSMC and SMC into SMC.

Finally, if a cell was segmented by the H&E-based AI algorithm as a megakaryocyte, we relabelled that cell as a megakaryocyte. If a cell was identified as a megakaryocyte on unsupervised clustering but not identified by the H&E-based computational algorithm, it was deemed a megakaryocyte.

### Annotation of H&E-based features

#### Adipocyte Segmentation

To identify adipocytes we used *MarrowQuant* 2.0^19^ with QuPath version 0.5^51^. To ensure accurate segmentation, we also then manually inspected the output of the automated segmentation for each sample and manually annotated any adipocytes not captured by automated segmentation.

#### Bone segmentation

To segment bones, we used our previously described bone segmentation algorithm, which uses a UNet46 architecture^18^. The bone structure was segmented at 5x magnification (Supplementary Fig. 6) and tiny bone fragments were excluded from downstream analysis. We then visualised the bone segmentation overlying the H&E-images to confirm segmentation accuracy (Supplementary Fig. 6).

#### Vessel annotation

We performed manual pathologist-led annotation of arterioles and large sinusoids on the H&E image (Fig. 3A). Using the previously described image registration anchors, the corresponding coordinates in the ST-data were identified. The annotation was external to the vessel wall i.e. external to endothelial cells for sinusoids and tunica externa for arterioles.

#### Automated fibrosis detection

Fibrosis was quantified using a reticulin-free H&E-based AI algorithm as previously described^30^. In brief, the model was trained using pairs of 512 x 512 (113 µ m x 113 µ m) H&E tiles at 40× magnification, ranked by fibrosis severity score based on the Continuous Indexing of Fibrosis (CIF) score. These were also aligned to WHO fibrosis grades (0-3) as previously described^18^. This reticulin-free model was then applied to non-overlapping tiles of size 512 x 512 pixels at 40x magnification. The tile-level fibrosis scores were visualised as a colour map overlaid onto the WSI to visualise overall fibrosis severity (Extended Data Fig. 8A). Correlation analyses involving the CIF score were also performed. Specifically, we evaluated: (1) the correlation between CIF and cell abundance, and (2) the correlation between CIF and the distance from bone and, (3) the correlation between CIF and the ST-micro score. The correlation between CIF and cell abundance was evaluated by calculating the mean CIF score across tiles in each intertrabecular space (ITS) and the proportion of each cell type. For the correlation between CIF and distance from bone we measured, within each intertrabecular space, the distance from bone to the centre of each CIF-predicted tile. Distances were then binned using the tile size (512 pixels), and the average CIF score within each bin was computed. Using these bin-wise average CIF scores and distances, we performed correlation analysis. Correlation between fibrosis and ST-micro score was performed by calculating the mean CIF score per intertrabecular space and then correlating this with the cellular proportion. All correlation analyses were performed using Pearson’s correlation.

#### Immunohistochemical cell detection and quantitation

To validate ST cell type annotation and quantitation, we compared the ST cell quantitation to that of immunohistochemistry (IHC)-based detection. Whole slide images (WSI) of key IHC markers (CD3, CD20, Ret40f) were obtained using a Hamamatsu NanoZoomer S210 digital slide scanner. The lymphoid markers (CD3, CD20) were analysed in QuPath (v0.5.1)^51^, using the ‘positive cell detection’ function with default parameters. The detections were manually inspected by a haematopathologist to confirm adequate performance. The cell detection output was exported for further analysis.

To quantitate erythroid populations we used Ret40f (glycophorin C) IHC. As positive staining is seen in both the nucleated erythroid precursors (the bone-marrow resident population of interest) and in red blood cells, we developed a custom image processing tool to detect only nucleated cells with Ret40f expression. This method uses classical (non-AI) image processing approaches to detect cells with a brown ring of 3,3-diaminobenzidine (DAB) positivity surrounding a central blue haematoxylin (H) stained nucleus. First, WSI were tiled and only tissue-containing tiles analysed. Each individual tile was split into H and DAB images using colour deconvolution, as described by Ruifrok^52^ and implemented in rgb2hed function from the color module in scikit-image^53^. The subsequent framework largely used utilities within OpenCV library^54^. The H and DAB images were thresholded using Otsu’s method^55^ and individual object contours were extracted. The DAB image contours were filled and the filled image was subtracted from the original thresholded DAB image. The resultant image was compared with the thresholded H image and only contours shared between H and filled DAB image were retained. These contours were further filtered based on non-restrictive shape and size criteria to remove artefactual detections well outside of the ranges expected for erythroid cells. The resultant contours were recorded and counted as ‘positive’ cells.

The WSI-level total positive cell detection counts for each sample and IHC marker were compared to their respective ST-based cell type assignment counts (i.e. CD3 vs T-cells, CD20 vs B-cells, Ret40f vs erythroids). To account for the differences in tissue area between ST data and IHC image that occurred due to upstream tissue microdissection, or post-hoc morphology driven-QC, we applied tissue area normalisation. The ST tissue area was calculated as the total area of the cytomorphologically preserved tissue (i.e. total tissue mask area minus the total area of regions without cytomorphological preservation). The IHC tissue area was calculated as the pixel count of the thresholded IHC image (on the same image magnification). The IHC counts were area normalised via multiplication with a ratio of ST area to IHC area. The area normalised IHC counts were correlated with the ST-based cell counts (Pearson’s correlation).

#### Differential abundance analysis

A negative-binomial generalised linear model *edgeR*^56^ was used to perform global differential abundance analysis of cell types across conditions using Benjamini and Hochberg procedure for adjustment for multiple testing (FDR < 0.05).

## SPATIAL ANALYSIS

### Regional enrichment analysis

To explore the differential abundance of BM cell populations across the marrow we defined supervised spatial regions defined by distance from bony trabeculae. Informed by conventional histological descriptions of the BM, we defined the endosteal space as that containing cells touching bone, peritrabecular space as the region immediately distal to the endosteal space (within 300 pixels of bone – approximately two fat spaces, which typically are seen in the normal BM) and intertrabecular (or central) space as that lying between peritrabecular spaces (Fig. 3G). To perform differential abundance analysis of cell types BM regions (endosteal/peritrabecular/intertrabecular) we used a negative-binomial generalised linear model *edgeR*, with models fitted with *glmFit* and hypothesis testing with *glmLRT*, with adjustment for multiple testing with Benjamini & Hochberg procedure to identify significant comparisons (FDR < 0.05). Sample ID was accounted for as a factor.

### Point pattern proximity analysis

We assessed the proximity of cell types and cell neighbourhoods (CNs) to microanatomical structures (bone, adipocytes, arterioles and sinusoids), as well as between different cell types, using a Poisson point process (PPP) model to model random distributions of points in space as reported by Bandyopadhyay et al.^9^. A key characteristic of PPP is the independence of events (in this case, occurrence of cell) in non-overlapping regions, with a Poisson distribution which is the probability distribution that describes the likelihood of a given number of events occurring within a fixed interval of space. PPP model is fitting the data based on:

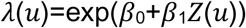

where u is the x-y coordinates of a cell, λ(u) is likelihood of occurrence of a cell, β_0_ is the baseline cell population where there is no covariate (in this case, distance to the structure or cell type of interest), Z(u) is distance to structure or cell type of interest, β_1_ is decay rate of Z(u) which represents proximity. As β_1_ value gets smaller, the cells are more likely to be found close to the structure.

To assess proximity of architectural features, we used only a subregion of the whole tissue. Unlike cells, arterioles and sinusoids are not uniformly distributed across the tissue. If proximity were computed based on the entire tissue, it could lead to inaccurate measurements. Therefore, we focused on only the regions local to arterioles and sinusoids rather than using the whole tissue. To extract the subregion, we first identified, for each arteriole or sinusoid, the closest structure among bone, adipocytes, arterioles and sinusoids. We then computed the distances from the arteriole or sinusoid to each of these nearest structures and selected the largest value among them. This maximum distance served as the half-length of a square subregion. Starting from the centre of the arteriole or sinusoid, we expanded outward by this distance in all directions, and the resulting minimum and maximum x–y coordinates defined the upper-left and lower-right corners of the region of interest. By doing so, we ensured that the resulting subregion included all types of structures. Since we use a subregion-based approach, proximity between structures is assessed only in the samples that contain all structures (17 out of 32 samples).

Since PPP requires a complete view of the spatial domain, we first generated the distance map for the structure or cell type of interest using the annotation mask for each sample. For each cell type, we then computed PPP using the x-y coordinates of the centres of the observed cells. Once β_1_ values were obtained for each cell type, we ranked these values for the cells in each sample and normalised them to a range of 0-1. To compare across different MPN subtypes, we used the median normalised proximity rank for each MPN subtype.

Additionally, we employed a permutation test to ensure that the spatial arrangement of cell types relative to the target structure or cell type was biologically meaningful and not random. We performed 100 permutations of the cell type labels and computed the median distance for each permutation. The p-value was calculated as the proportion of distances smaller than the observed cell type distance. To aggregate p-values across cell types, we applied Stouffer’s method to combine the permutation test p-values. The PPP model was implemented in the *spatstat R* package, and statistical analysis carried out in Python.

### Cell neighbourhood analysis

#### Graph construction

Prior to unsupervised cell neighbourhood analysis with *CellCharter*^20^, we performed graph construction. We constructed a graph representation of each tissue section by defining edges between cells whose Xenium-segmented cell-boundary polygons touch or overlap after a small expansion. Specifically, we expand (buffer) each cell boundary polygon outwards by 2 µm to identify, for every cell, all other cells whose expanded boundary intersect its own. Each intersecting pair (i.e. touching polygon) was treated as a direct cell–cell contact and recorded as an undirected edge between the corresponding cell centroids in a sparse adjacency matrix. To manage irregularly shaped cells the graph was constructed between centroids of all cells in the intertrabecular space (ITS) regions (i.e. areas with good architectural preservation). We refer to the resulting graph as a topological adjacency graph. Compared with Delaunay-based graphs (with and without distance thresholds), this boundary-based graph more closely matched qualitative expectations of spatial proximity and neighborhood relationships in the tissue (Supplementary Fig. 7&8).

#### Unsupervised neighbourhood analysis

We used *CellCharter*^20^ to identify unsupervised spatial cell neighbourhoods (CNs). *CellCharter* is a graph-based analysis and is scalable across mulitple samples. We used the topological adjacency graph (see above) to define cell connectivity. We performed log transformation of gene expression data, and performed dimentionality reduction using the cVI model (scvi.model.SCVI). The model was trained with sample ID and batch as variables and cell annotations as labels, outputting a 128-dimensional latent representation per cell. *CellCharter* concatenates each cell’s latent vector from gene expression and cell annotations, and those of its neighbours to up k hops. For k = 3 this results in four vectors: the cell in question, and the averaged vectors of first, second and third hop neighbours. These combined vectors were then clustered into a user-defined number of spatial domains using a Gaussian Mixture model. We selected n = 10 CNs, informed by both cluster stability metrics (calculated with average Fowlkes-Mallows Index between n–1, n, and n+1) and prior knowledge of bone marrow biology.

We compared the size of CNs across conditions (normal, PV, ET, MF). To do this we first identified connected components in the topological adjacency graph using the function cc.gr.connected_components(), with a minimum cell threshold of 5, thereby retaining only CNs containing at least five cells. For each sample, counts were normalised by the total number of cells in that sample to obtain proportions, thereby accounting for differences in overall cellularity across samples. We then compared the median CN component size per sample across conditions with a Mann-Whitney U test. We also compared the median CN size of the ten largest CN components (Mann-Whitney U), and then compared the median size of the largest CN components across CN types within each condition (Wilcoxon signed-rank test).

#### Cell neighbourhoods versus Continuous Index of Fibrosis score

We quantified the fibrosis (Continuous Index of Fibrosis (CIF)) within different CNs. To do this we calculated the corresponding median CIF of CN regions by condition, applying a linear mixed effects model with Benamini-Hochberg correction for comparison between CNs.

### Spatial pattern analysis

To characterise the patterns of the spatial landscape of cells within the BM samples, we used spatial autocorrelation analysis. To run these methods we utilised the Squidpy^57^ toolkit. We first constructed a graph representation of a tissue section (whole trephine or intertrabecular region), wherein nodes represent individual cells and edges encode spatial connectivity or adjacency. Following graph construction spatial statistics could be computed as outlined below. To define spatial relationships among cells within a given ITS of a tissue section, we constructed its graph representation using Squidpy’s *sq.gr.spatial_neighbors* function. The function outputs a cell-cell adjacency matrix (spatial_connectivities) and pairwise euclidean distances between cells (spatial_distances), providing a foundation for downstream spatial analyses.

#### Spatial autocorrelation

To quantify the spatial organisation of cell types, we computed Moran’s I, a measure of spatial autocorrelation that compares the value of a feature at one location to values at neighboring locations. Moran’s I is defined as follow:

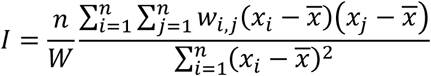

where *n* denotes the total number of spatial points under consideration (in our case cell *x, y* location across a tissue section), *w_i,j_* is the connectivity/adjacency between two cells *i* and *j*, and *W* = ∑*_i_* ∑*_j_ w_ij_* is the normalisation factor, calculated as the sum of all spatial weights. *x_i_* represents the feature values at location *i* (e.g., presence or absence of a given cell type), and *x̅* is the mean values of *x* across the tissue section. The value of *I* ranges from -1 (perfect dispersion) to 1 (clustering), with 0 suggesting nearly random spatial arrangement.

We performed this spatial autocorrelation analysis on binary vectors (presence =1 and absence = 0) of each individual cell types cell types at resolution 3. Moran’s I was computed using the sq.gr.spatial_autocorr() function from Squidpy with parameters mode=“moran” and n_perm = 1000. The computed Moran’s I statistics and the multiple-hypothesis corrected *p*-values across 1,000 permutations were extracted from the *moranI* field of the resulting AnnData object.

### Data-driven identification of microenvironmental spatiotypes

To capture microenvironmental patterns for the purpose of exploring disease biology, for hypothesis generation and to capture intra-and inter-sample heterogeneity we used our recently proposed ‘band descriptor’ approach^37^. The method characterises the microenvironment of each cell using a set of concentric bands, with number of bands and band width user-defined parameters. In our analysis, we used five concentric bands (100, 200, 300, 400, 500 pixels). Within each band, we quantified the relative abundance of neighboring cell types, resulting in a multi-scale, spatially-aware descriptor for each cell. We then applied K-means clustering (k=3) to these descriptors to identify distinct spatiotypes representing recurring patterns of local microenvironmental enrichment across the entire cohort. Clustering identifies three spatiotypes per cell type, effectively capturing representative spatial diversity of cells across multiple tissue sections while potentially reducing the bias introduced either by cellularity or spatial scale selection.

#### Characterising spatiotypes in MPN

We assessed the enrichment of spatiotypes across disease subtypes by computing the correlation between spatiotype abundance at the ITS-region level and corresponding patient level disease labels. Specifically, we computed the abundance of each spatiotype within individual ITS regions and assigned the corresponding patient-level disease label to each region. For example, if an ITS region originated from a normal sample, we assigned it the label ‘normal’. We then one-hot encoded the ITS-region-level disease labels for each disease subtype, indicating whether a region belonged to a specific subtype (1) or not (0). Kendall’s rank correlation coefficient was then computed between each spatiotype and disease subtype to assess monotonic relationships between spatial patterns and disease states. For each spatiotype we reported the correlation values along with Benjamini–Hochberg-adjusted p-values. This allowed us to identify spatiotypes that are significantly enriched or depleted within a specific disease subtype.

#### ML model for linking spatial cellular microenvironmental patterns to disease status

To capture microenvironmental heterogeneity within a sample and across the cohort, we developed a weakly supervised multiple instance learning (MIL)-based framework that learns a mapping between spatial cell-arrangements (spatiotypes) and disease status using only biopsy-level labels. Inspired by previous aggregation-based MIL approaches^58^, each BMT was represented as a bag *B* = {*x*_1_, *x*_2_, …, *x_n_*}, where each *x_i_* ∈ ℝ*^d^* represents a *d*-dimensional spatiotype abundance vector for the *i*-th ITS-region, and *n* is the total number of ITS regions in the biopsy. The model architecture consists of an instance-scoring module *f*(*x*) = *w T x* + *b*, followed by a top-k MIL aggregation. The output of the scoring module *f*(*x*) reflects the contribution of an ITS-region to the biopsy-level prediction. To learn the parameters ***w*** and ***b***, we use a top-k ranking loss. For each positive bag and its paired negative bags, the model *f*(∴) predicts instance-level scores and then selects the *k* highest scoring regions from each. Let ***s***^⁺^ denote the low-scoring regions from the positive bag, ***s***^⁻^ denotes the high-scoring regions from the negative bag; then the loss is computed as follows:

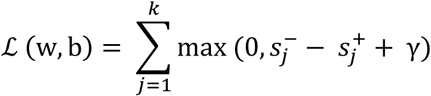

The loss function enforces the model to rank ITS regions in the positive bag higher than in the negative bag by a margin *γ*. Additionally, we applied L1 regularisation (*λ*|*w*| with *λ* = 0.001) to promote sparsity in the learned weight vector. The final objective is minimised using stochastic gradient descent, encouraging the model to focus on spatial features most predictive of disease.

#### Model Training and Evaluation

To evaluate model performance, we used patient-level stratified bootstrapping strategy to ensure unbiased training and testing. As some patients contributed multiple tissue sections, sampling was performed at the patient level to avoid data leakage across splits. In each bootstrap iteration, patients we randomly sampled with replacement, using 67% of the data for training and 33% for testing. The model training was perfomed only on normal (n = 5) and MF (n = 11) cases. The performance of the trained model was evaluated in two ways:

1. In-sample evaluation on the held-out normal and MF cases in each run.
2. Out-of-sample evaluation on additional diagnostic groups not seen during training, specifically ET and PV cases.

We perform *n* = 20 bootstrap runs, training the model for 200 epochs using an Adaptive momentum-based (Adam)^59^ optimiser with a learning rate of 0.01 and weight decay of 0.01. Model predictive performance was quantified using the Area Under the Receiver Operating Characteristic Curve (AUROC), and mean value across all runs was reported.

To further assess the discriminative ability of the trained models in inferring disease trajectory (normal to MF), we generated ITS-level prediction for all samples in the cohort and aggregated the predictions across all bootstrap model using mean, yielding a single score per ITS region. This ensemble prediction strategy enabled us to position patients along a disease continuum, using normal and MF samples as anchors - normal cases at the lower end and MF cases at the upper extreme of the trajectory.

#### Model Explainability

Given the linear nature of our model, interpretability is intrinsic to its design. Each trained model produces a weight vector that can be directly interpreted as the contribution of each spatiotype feature to the prediction. To assess cohort-level (global) explanations, we recorded the learned weight vector from 20 bootstrap runs. This allowed us to examine the median, variability, and directionality of each feature’s weight — indicating whether a given spatiotype consistently contributed positively or negatively to the models’ predictions across the cohort. For ITS-region-level (local) explanations, the aggregated weight vector from different ensemble model can be multiplied with the spatiotype abundance vector. This yields both a scalar contribution score for that region’s influence on the slide-level prediction, as well as feature-level effect values, computed via element-wise multiplication of the regional feature vector and the global weight vector. Together, these global and local perspectives enable interpretation of which spatial features drive disease predictions across the cohort and within specific tissue regions.

## Supporting information

Table of Cell type DEGs

## Acknowledgments

This work was funded by Blood Cancer UK (Grant/Award Number: 23012); Cancer Research UK (Grant/Award Number: EDDPJT-May23/100034); a Wellcome Trust Bioimaging Technology Award (313477/Z/24/Z); a EPSRC-funded Seebibyte programme (EP/M013774/1); the MPN Research Foundation^TM^ and the Ludwig Institute for Cancer Research, Oxford Branch. RAC, AS and MG are funded by the Jean Shanks Foundation / Pathological Society of Great Britain and Ireland. ET was supported by a EPSRC Grant (number EP/S024093/1). JR is an adjunct professor of the Ludwig Oxford Branch. BP is a a Senior Fellowship funded by Cancer Research UK in partnership with the Rosetrees Trust (to BP) and the Ludwig Institute for Cancer Research (Associate Member Program Grant to BP). FI and OM are supported by the Chinese Academy of Medical Sciences (CAMS) Innovation Fund for Medical Science (CIFMS), China (grant number: 2024-I2M-2-001-1). OM was supported by a Medical Research Council Clinical Research Training Fellowship, award reference: MR/V000942/1 and Royal College of Surgeons of England Research Fellowship.

The authors thank those patients who gave permission for their samples to be used in this study. We would like to thank Alan King Lun Liu for valuable discussion and advice.

## Author Contributions

RAC and DJR conceived the project. RAC, DJR, JR and DW designed the study. DJR, RAC and JR organised and oversaw the study. ET, MD, HR, AS, CP, RAC, DW and MG analysed the data and interpreted results. ET, MD, HR, AS, CP, MG and RC generated figures. OM and RT generated data. DJR and CP performed pathological review and annotation. JR, DJR, BP, AM, FI and JH provided resources. DJR, RAC and JR provided funding support. RAC, ET, MD and DJR wrote the paper. ET, MD, HR, RC, DJR and JR developed AI-based analysis method, with MD generating the model architecture/code. RAC, JR, DJR, DW, BP and AM made a substantial contribution to the organisation and conduct of the study and critiqued the output for important intellectual content. All authors reviewed and approved the final copy of the manuscript.

## Data availability & Code availability

Code to support this paper is available at: https://github.com/imuhdawood/MarrowMap Data is available on request from the corresponding author.

## References

1. Jacobsen, S. E. W. & Nerlov, C. Haematopoiesis in the era of advanced single-cell technologies. Nature Cell Biology 2019 21:1 21, 2–8 (2019).

2. Tikhonova, A. N. et al. The bone marrow microenvironment at single-cell resolution. Nature 2019 569:7755 569, 222–228 (2019).

3. O’Sullivan, J. M., Mead, A. J. & Psaila, B. Single-cell methods in myeloproliferative neoplasms: old questions, new technologies. Blood 141, 380–390 (2023).

4. Janesick, A. et al. High resolution mapping of the tumor microenvironment using integrated single-cell, spatial and in situ analysis. Nature Communications 2023 14:1 14, 1–15 (2023).

5. Baccin, C. et al. Combined single-cell and spatial transcriptomics reveal the molecular, cellular and spatial bone marrow niche organization. Nature Cell Biology 2019 22:1 22, 38–48 (2019).

6. Sarachakov, A. et al. Spatial mapping of human hematopoiesis at single-cell resolution reveals aging-associated topographic remodeling. Blood 142, 2282–2295 (2023).

7. Muiños-Lopez, E. et al. Characterization of the bone marrow architecture of multiple myeloma using spatial transcriptomics. Communications Biology 2025 8:1 8, 1620- (2025).

8. Yip, R. K. H. et al. Profiling the spatial architecture of multiple myeloma in human bone marrow trephine biopsy specimens with spatial transcriptomics. Blood 146, 1837–1849 (2025).

9. Bandyopadhyay, S. et al. Mapping the cellular biogeography of human bone marrow niches using single-cell transcriptomics and proteomic imaging. Cell 187, 3120–3140.e29 (2024).

10. Mead, A. J. & Mullally, A. Myeloproliferative neoplasm stem cells. Blood 129, 1607–1616 (2017).

11. Tefferi, A. et al. Long-term survival and blast transformation in molecularly annotated essential thrombocythemia, polycythemia vera, and myelofibrosis. Blood 124, 2507 (2014).

12. Li, R. et al. A proinflammatory stem cell niche drives myelofibrosis through a targetable galectin-1 axis. Sci Transl Med 16, (2024).

13. Gleitz, H. L. F. E. et al. Increased CXCL4 expression in hematopoietic cells links inflammation and progression of bone marrow fibrosis in MPN. Blood 136, 2051 (2020).

14. Leimkühler, N. B., Costa, I. G. & Schneider, R. K. From cell to cell: Identification of actionable targets in bone marrow fibrosis using single-cell technologies. Exp Hematol 104, 48–54 (2021).

15. Cooper, R. A., Thomas, E., Sozanska, A. M., Pescia, C. & Royston, D. J. Spatial transcriptomic approaches for characterising the bone marrow landscape: pitfalls and potential. Leukemia 2024 1–5 (2024) doi:10.1038/s41375-024-02480-8.

16. Sirinukunwattana, K. et al. Artificial intelligence–based morphological fingerprinting of megakaryocytes: a new tool for assessing disease in MPN patients. Blood Adv 4, 3284–3294 (2020).

17. Azimuth. https://azimuth.hubmapconsortium.org/.

18. Ryou, H. et al. Continuous Indexing of Fibrosis (CIF): improving the assessment and classification of MPN patients. Leukemia 2022 5, 1–11 (2022).

19. Sarkis, R. et al. MarrowQuant 2.0: A Digital Pathology Workflow Assisting Bone Marrow Evaluation in Experimental and Clinical Hematology. Modern Pathology 36, 100088 (2023).

20. Varrone, M., Tavernari, D., Santamaria-Martínez, A., Walsh, L. A. & Ciriello, G. CellCharter reveals spatial cell niches associated with tissue remodeling and cell plasticity. Nature Genetics 2023 56:1 56, 74–84 (2023).

21. Psaila, B., Lyden, D. & Roberts, I. Megakaryocytes, Malignancy and Bone Marrow Vascular Niches. Journal of Thrombosis and Haemostasis 10, 177 (2012).

22. Spatial Point Patterns: Methodology and Applications with R - Adrian Baddeley, Ege Rubak, Rolf Turner - Google Books. https://books.google.co.uk/books?hl=en&lr=&id=rGbmCgAAQBAJ&oi=fnd&pg=PP1&dq=A.BaddeleyE.RubakR.Turner+Spatial+point+patterns:+methodology+and+applications+with+RFirst+edition+2015+Routledge&ots=2zXPpOGbwy&sig=mImRVW7lzEhMRCSCgTfLaTQNzic&redir_esc=y#v=onepage&q&f=false.

23. Méndez-Ferrer, S. et al. Bone marrow niches in haematological malignancies. Nat Rev Cancer 20, 285 (2020).

24. Morrison, S. J. & Scadden, D. T. The bone marrow niche for haematopoietic stem cells. Nature 2014 505:7483 505, 327–334 (2014).

25. Furuhashi, K. et al. Bone marrow niches orchestrate stem-cell hierarchy and immune tolerance. Nature 2025 638:8049 638, 206–215 (2025).

26. Zhou, B. O. et al. Bone marrow adipocytes promote the regeneration of stem cells and hematopoiesis by secreting SCF. Nat Cell Biol 19, 891 (2017).

27. Chen, J., Hendriks, M., Chatzis, A., Ramasamy, S. K. & Kusumbe, A. P. Bone Vasculature and Bone Marrow Vascular Niches in Health and Disease. Journal of Bone and Mineral Research 35, 2103–2120 (2020).

28. Owen-Woods, C. & Kusumbe, A. Fundamentals of bone vasculature: Specialization, interactions and functions. Semin Cell Dev Biol 123, 36–47 (2022).

29. Passamonti, F. & Mora, B. Myelofibrosis. Blood 141, 1954–1970 (2023).

30. Ryou, H., et al. Reticulin-Free Quantitation of Bone Marrow Fibrosis in MPNs: Utility and Applications. EJHaem 6, e70005 (2025).

31. Cooper, R. et al. Mapping the Human Bone Marrow in Myeloproliferative Neoplasia Using Spatial Transcriptomics. Blood 144, 876–876 (2024).

32. Yao, J. C. et al. TGF-β signaling in myeloproliferative neoplasms contributes to myelofibrosis without disrupting the hematopoietic niche. J Clin Invest 132, e154092 (2022).

33. Vukotić, M. et al. Inhibition of proinflammatory signaling impairs fibrosis of bone marrow mesenchymal stromal cells in myeloproliferative neoplasms. Experimental & Molecular Medicine 2022 54:3 54, 273–284 (2022).

34. Gianelli, U. et al. The European Consensus on grading of bone marrow fibrosis allows a better prognostication of patients with primary myelofibrosis. Modern Pathology 2012 25:9 25, 1193–1202 (2012).

35. WHO Classification of Tumours Online. https://tumourclassification.iarc.who.int/welcome/.

36. Banjanin, B. et al. Wnt-dependent spatiotemporal reprogramming of bone marrow niches drives fibrosis. bioRxiv 2025.02.12.637594 (2025) doi:10.1101/2025.02.12.637594.

37. Dawood, M. et al. Unsupervised Discovery of Spatiotypes and Context-Aware Graph Neural Networks for Modeling Clinical Endpoints. 603–612 (2026) doi:10.1007/978-3-032-05141-7_58.

38. Lin, W. et al. Mapping the spatial atlas of the human bone tissue integrating spatial and single-cell transcriptomics. Nucleic Acids Res 53, (2025).

39. Mattiucci, D. et al. Bone marrow adipocytes support hematopoietic stem cell survival. J Cell Physiol 233, 1500–1511 (2018).

40. Zhou, B. O. et al. Bone marrow adipocytes promote the regeneration of stem cells and haematopoiesis by secreting SCF. Nat Cell Biol 19, 891–903 (2017).

41. Grockowiak, E. et al. Different niches for stem cells carrying the same oncogenic driver affect pathogenesis and therapy response in myeloproliferative neoplasms. Nature Cancer 2023 4:8 4, 1193–1209 (2023).

42. Nombela-Arrieta, C. & Manz, M. G. Quantification and three-dimensional microanatomical organization of the bone marrow. Blood Adv 1, 407 (2017).

43. Gleitz, H. F. E. et al. Inhibiting the alarmin-driven hematopoiesis-stromal cell crosstalk in primary myelofibrosis ameliorates bone marrow fibrosis. Hemasphere 9, e70179 (2025).

44. Leimkühler, N. B. et al. Heterogeneous bone-marrow stromal progenitors drive myelofibrosis via a druggable alarmin axis. Cell Stem Cell 28, 637–652.e8 (2021).

45. Fan, W. et al. Contributions of bone marrow monocytes/macrophages in myeloproliferative neoplasms with JAK2V617F mutation. Ann Hematol 102, 1745 (2023).

46. Silver, R. T., Chow, W., Orazi, A., Arles, S. P. & Goldsmith, S. J. Evaluation of WHO criteria for diagnosis of polycythemia vera: a prospective analysis. Blood 122, 1881–1886 (2013).

47. Blampey, Q. et al. Sopa: a technology-invariant pipeline for analyses of image-based spatial omics. Nature Communications 2024 15:1 15, 4981- (2024).

48. Butler, A., Hoffman, P., Smibert, P., Papalexi, E. & Satija, R. Integrating single-cell transcriptomic data across different conditions, technologies, and species. Nat Biotechnol 36, 411 (2018).

49. Korsunsky, I. et al. Fast, sensitive and accurate integration of single-cell data with Harmony. Nature Methods 2019 16:12 16, 1289–1296 (2019).

50. Aran, D. et al. Reference-based analysis of lung single-cell sequencing reveals a transitional profibrotic macrophage. Nat Immunol 20, 163–172 (2019).

51. Bankhead, P. et al. QuPath: Open source software for digital pathology image analysis. Scientific Reports 2017 7:1 7, 1–7 (2017).

52. Ruifrok, A. C. & Johnston, D. A. Quantification of histochemical staining by color deconvolution. Anal Quant Cytol Histol 23, 291–299 (2001).

53. Van Der Walt, S. et al. Scikit-image: Image processing in python. PeerJ 2014, e453 (2014).

54. Bradski, G. The OpenCV Library. Dr. Dobb’s Journal of Software Tools 120, 122–125 (2000).

55. Otsu, N. A Selection Threshold Method from Gray-Level Histograms. IEEE TRANSACTIONS ON SYSTREMS, MAN, AND CYBERNETICS, 9, 62–66 (1979).

56. Chen, Y., Chen, L., Lun, A. T. L., Baldoni, P. L. & Smyth, G. K. edgeR v4: powerful differential analysis of sequencing data with expanded functionality and improved support for small counts and larger datasets. Nucleic Acids Res 53, (2025).

57. Palla, G. et al. Squidpy: a scalable framework for spatial omics analysis. Nature Methods 2022 19:2 19, 171–178 (2022).

58. Asif, A. & Minhas, F. ul A. A. An embarrassingly simple approach to neural multiple instance classification. Pattern Recognit Lett 128, 474–479 (2019).

59. Kingma, D. P. & Ba, J. L. Adam: A Method for Stochastic Optimization. 3rd International Conference on Learning Representations, ICLR 2015 - Conference Track Proceedings https://arxiv.org/pdf/1412.6980 (2014).

