## Extended Data Figures for "Redefining the topology of the human bone marrow using augmented spatial transcriptomic analysis"

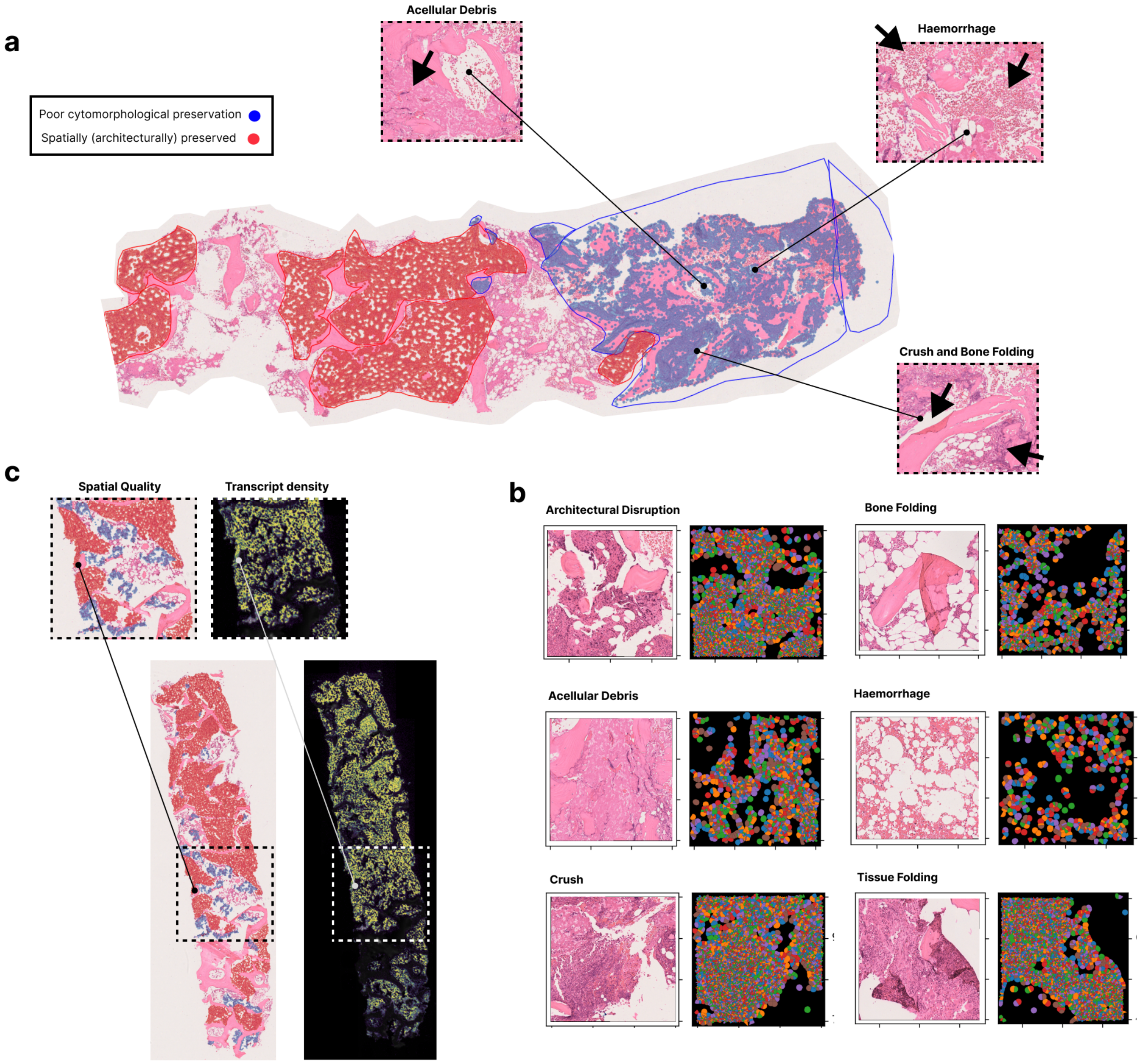


**Extended Data Figure 1: Visual inspection of ST data does not detect poorly preserved tissue regions unsuitable for downstream analysis**

**A:** Bone marrow trephine (BMT) with inset images demonstrating regions of acellular debris, haemorrhage, tissue crush and folding. Line annotations were drawn by a pathologist, identifying regions in which cells showed poor cytomorphological preservation (blue) and regions of good architectural tissue preservation (red). The latter broadly align to an intertrabecular space (ITS) (region between bone trabeculae). Areas of tissue outside of both of these annotated areas were preserved cytomorphologically (i.e. preserved cell shape and size) but lacked appropriate architectural preservation (i.e. normal tissue structure), therefore were suitable for analysis of cellular microenvironmental composition but excluded from spatial analyses. **B:** Tile H&E images (left) to show key examples of tissue features associated with lack of cytomorphological preservation (e.g. crush, tissue folding) or architectural disruption and the corresponding Xenium ST output (right) for the same region. **C:** BMT to show comparison of transcript density and tissue preservation. Image left shows a BMT with cells removed from analysis due to poor cytomorphological preservation highlighted in blue, and those in architecturally preserved regions in red. Image right shows the Xenium transcript density heatmap in the same region.

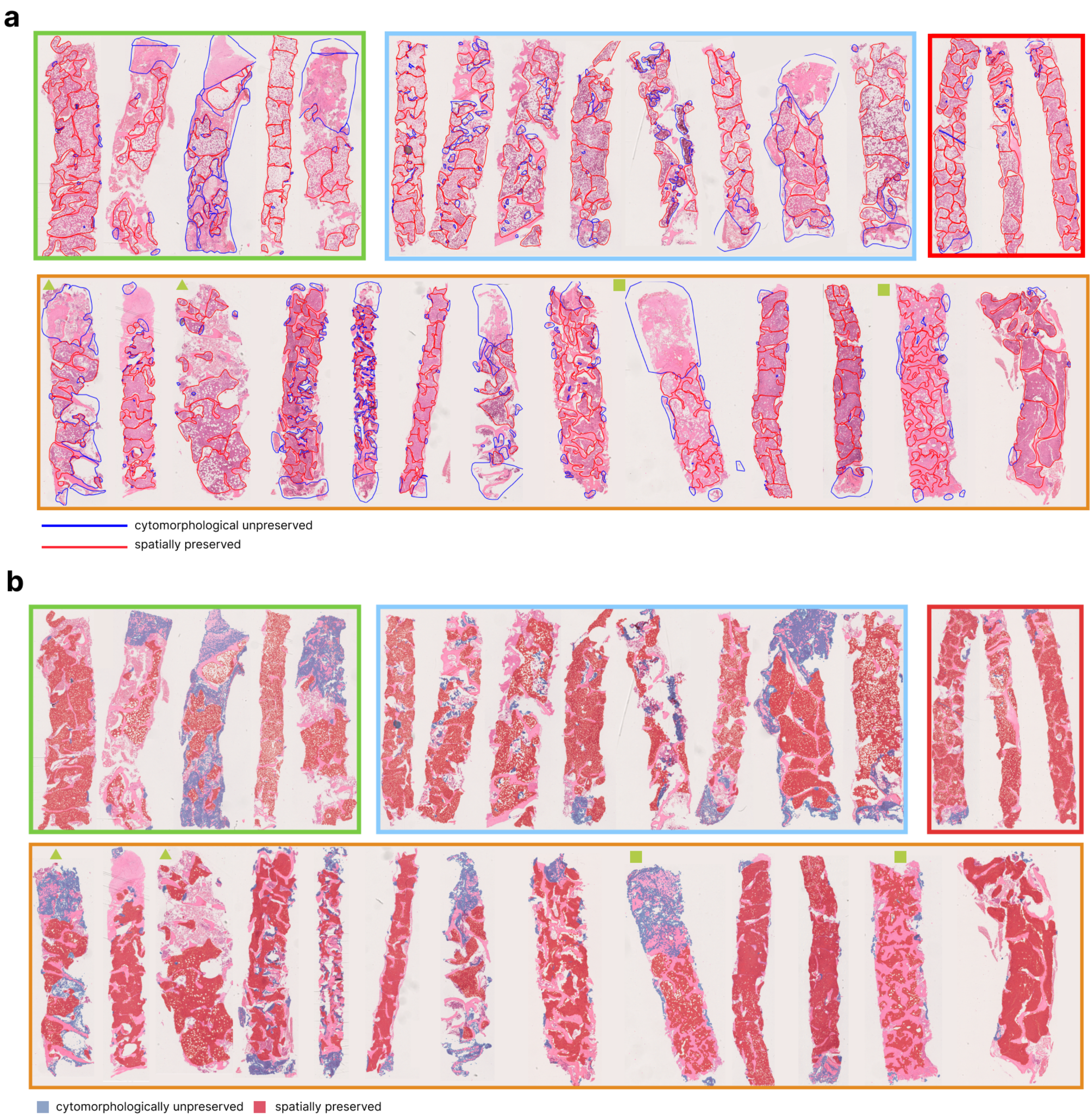


**Extended Data Figure 2: H&E-based annotation for spatial QC**

**A:** Bone marrow trephine (BMT) H&E images and haematopathologist annotations. Red line annotations indicate regions of good architectural preservation, while areas annotated in blue are regions of poor cellular preservation to be removed from any further downstream analysis. The trephines are grouped by condition (green – normal; blue – ET; red – PV and orange – MF). The images include 27 samples, with 18612_R1 and 18612_R4 (labelled with a yellow square); and 18612_R2 and 18612_R3 (yellow triangle) BMTs from the same BMT sample and bisected to facilitate tissue placement on the Xenium slide during processing. PrePMF samples (n=3) were annotated but are not shown as they were not taken forward for spatial analysis downstream. **B:** Bone marrow trephines to show ST data overlying the H&E image (as per Extended Data Fig. 2A) with cells coloured according to location within spatially preserved regions (red), or cytomorphological unpreserved regions to be removed (blue). The images include 27 samples, with 18612_R1 and 18612_R4; and 18612_R2 and 18612_R3 from the same BMT sample and bisected.

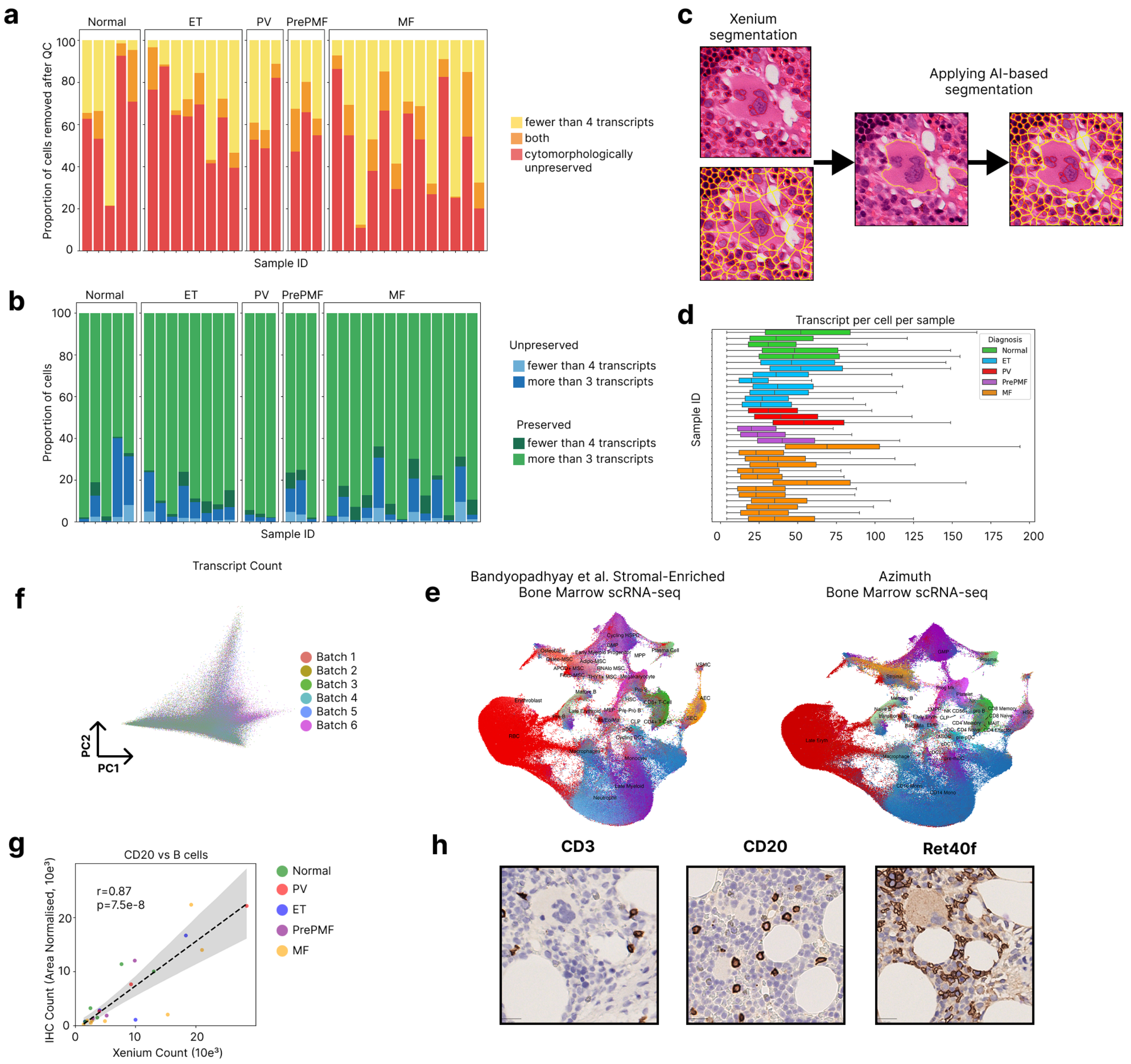


### Extended Data Figure 3: Data QC and cell type annotation

**A:** Stacked bar plot to show proportion of cells removed within a region of poor cytomorphological preservation (red), those with fewer than four transcripts (yellow), and those cells satisfying both criteria (orange). **B:** Stacked bar plot to show proportion of cells within cytomorphological unpreserved regions with more than three transcripts (dark blue) or fewer than four transcripts (light blue); and cells in cytomorphologically preserved regions with more than four transcripts (light green) or fewer than four transcripts (dark green) per sample. **C:** Tile images showing improved segmentation following insertion of AI-detected megakaryocytes into the original ST-based cell segmentation. Tile images show: Left, top – megakaryocyte nuclear segmentation with Xenium; Left, bottom – megakaryocyte segmentation with Xenium; middle – application of AI-based megakaryocyte detection algorithm; right: AI-detected megakaryocyte segmentation output. The red line indicates the nuclear boundary and yellow the cell boundary. **D:** Boxplot to show transcript counts per cell per sample and by condition (post-QC). **E:** UMAP plots to show supervised annotation of BM ST data with Azimuth[17] and Bandyopadhyay et al.[9] external reference datasets. All cells were annotated using *SingleR*. **F:** PCA plot to show dimensionality reduction across the ST cohort coloured by batch (1-6). **G:** Scatterplot to show correlation between proportion of cells in ST data and quantification of immunohistochemistry (IHC)-detected cells. Plot show correlation between B-cells as detected on ST data and CD20 IHC (n=23 BMTs) (Pearson's correlation). **H:** Images to show CD3, CD20 and Ret40f immunostains. For the Ret40F (glycophorin C) we developed an approach to quantify only nucleated cells (see Methods), although the stain also detects erythrocytes (e.g. regions of haemorrhage/blood).

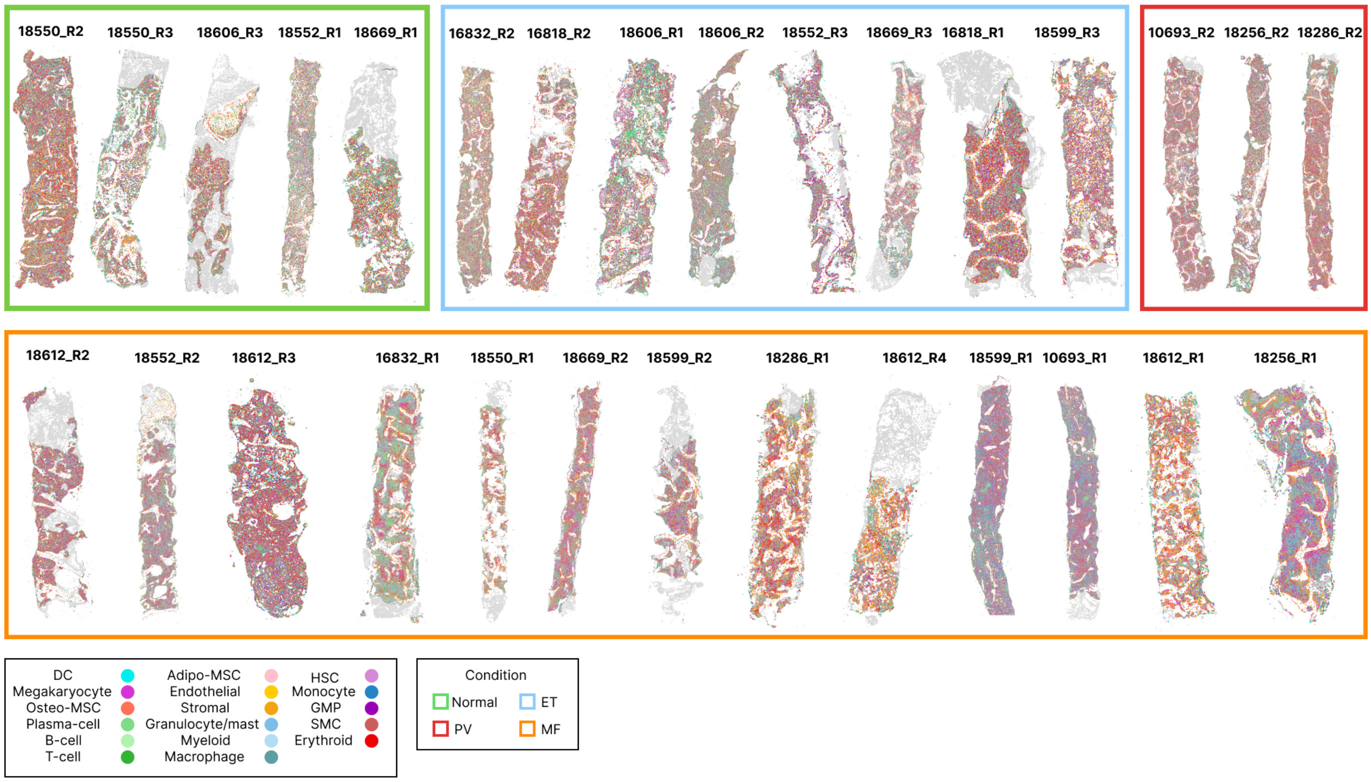


**Extended Data Figure 4: Bone marrow trephines and cell type annotation**
Bone marrow trephine (BMT) samples with cells coloured by type. BMTs are grouped by condition (green – normal, blue – ET, red – PV, orange – MF). Figure includes 27 samples; however, note that 18612_R1 and 18612_R4, and 18612_R2 and 18612_R3 are from the same BMT sample and bisected to facilitate tissue placement during processing. PrePMF samples are not visualised.

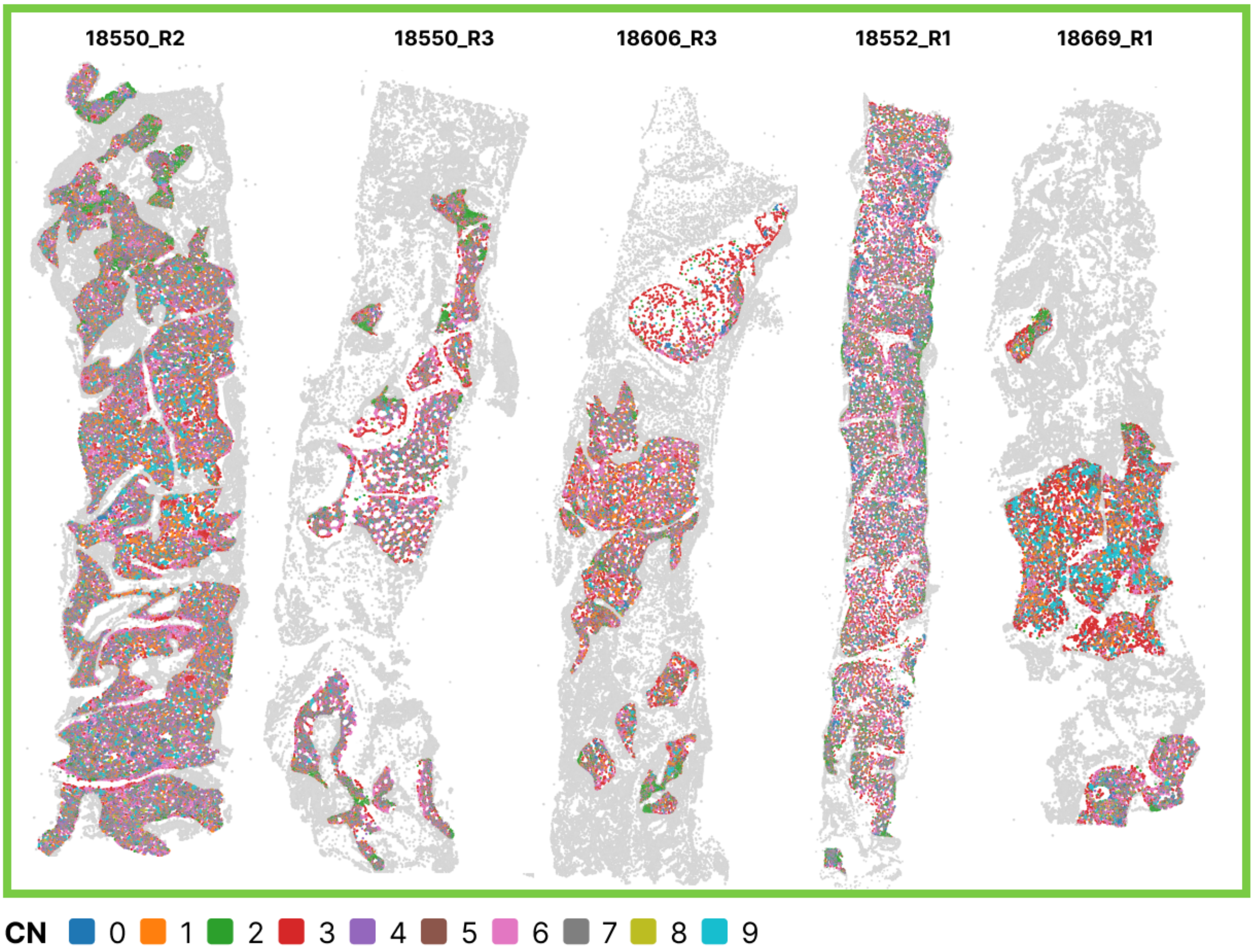


**Extended Data Figure 5: Bone marrow spatial cellular neighbourhoods**
Bone marrow trephine plots to show *CellCharter* derived cell neighbourhoods (CNs) across normal samples (n=5). Cells are coloured by CN (see key at bottom). Cells coloured in light grey are those lying outside of spatially preserved regions and not included in this analysis.

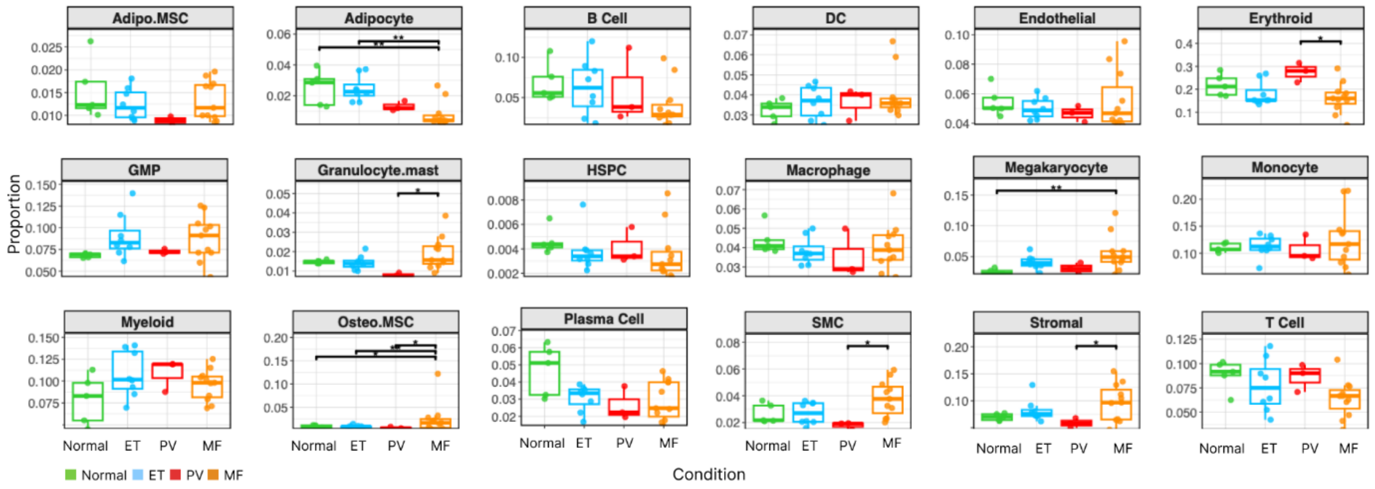


**Extended Data Figure 6: Differential cellular abundance in MPN**

Boxplot to show the differential abundance of cell types across conditions (n = 27, including five normal samples, three PV, eight ET and eleven MF BMTs) (negative-binomial generalised linear model with *edgeR* and Benjamini-Hochberg correction). Adipocyte category denotes segmented fat spaces detected by *MarrowQuant2.*

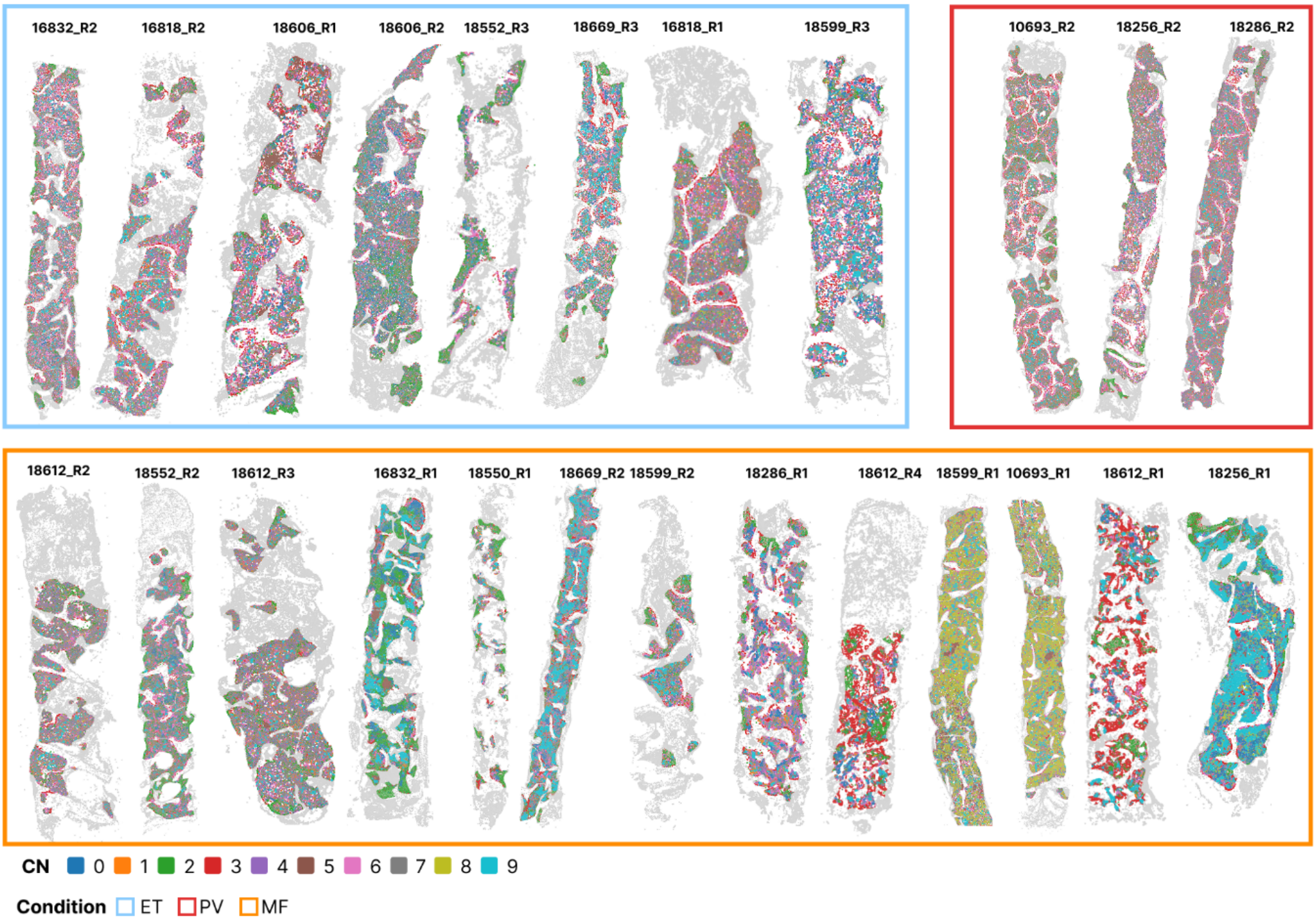


**Extended Data Figure 7: Cell neighbourhood analysis in MPN**

Bone marrow trephines (BMTs) to show cells coloured according to *CellCharter* cell neighbourhood (CN) assignment (CN0-9), grouped by condition (blue border – ET; red border – PV; orange border – MF). Cells coloured in grey lie outside of spatially preserved regions.

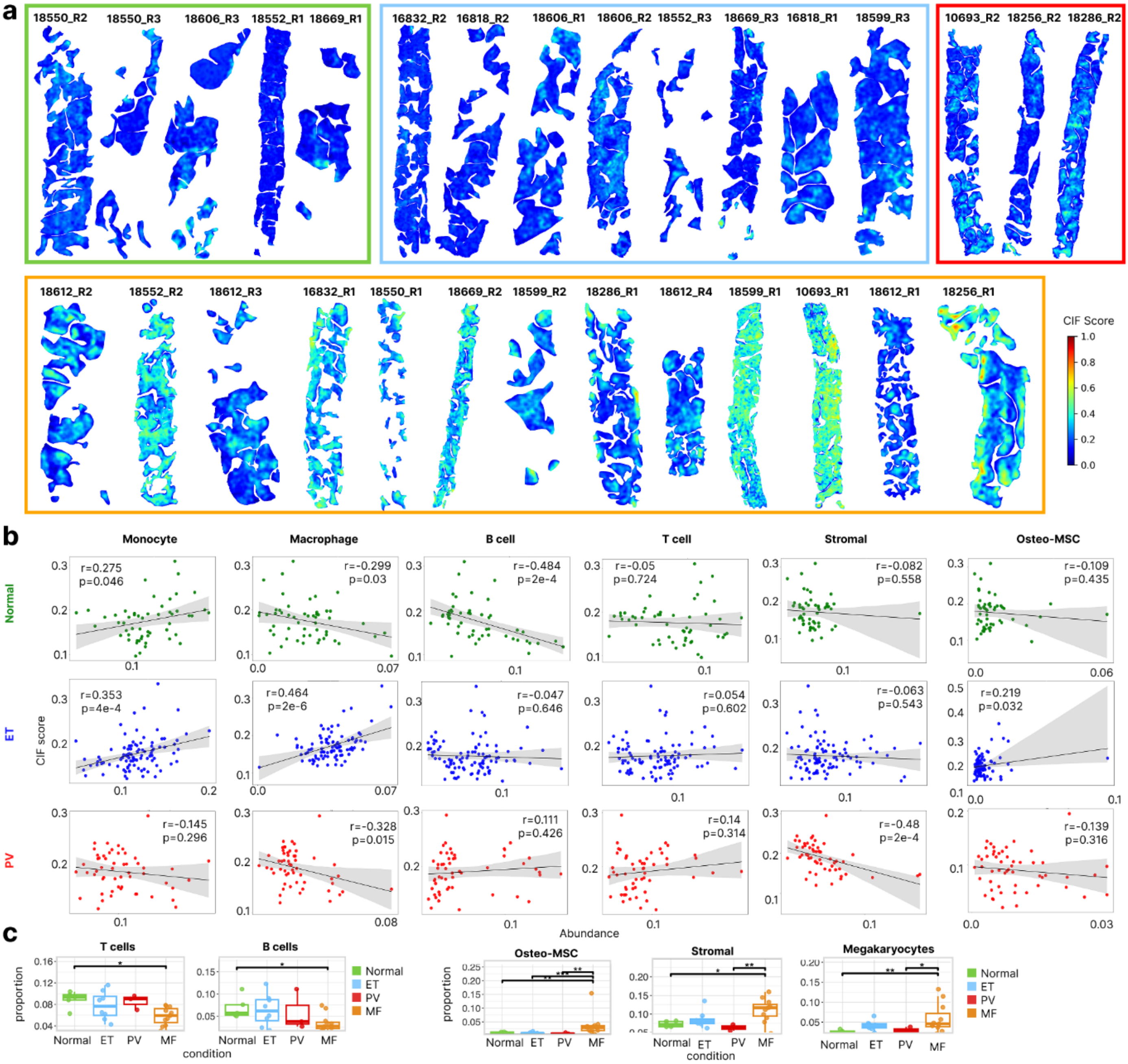

**Extended Data Figure 8: Microenvironmental correlates of bone marrow fibrosis**

**A:** Heatmaps to show the Continuous Index of Fibrosis (CIF) score per sample in intertrabecular space (ITS) regions. Areas of higher fibrosis are highlighted in yellow/orange/red, and regions of lower fibrosis are highlighted in blue/green (see colour scale right). **B:** Scatterplots to show correlation between CIF and cellular type proportion in normal, ET and PV (Pearson's correlation). The mean CIF score per ITS region was correlated with proportion of each cell type per ITS. **C:** Boxplots to show differential abundance of cell types in regions without increased fibrosis (WHO MF-0) across conditions (normal, PV, ET and MF). *p<0.05, **p<0.01, ***p<0.001.

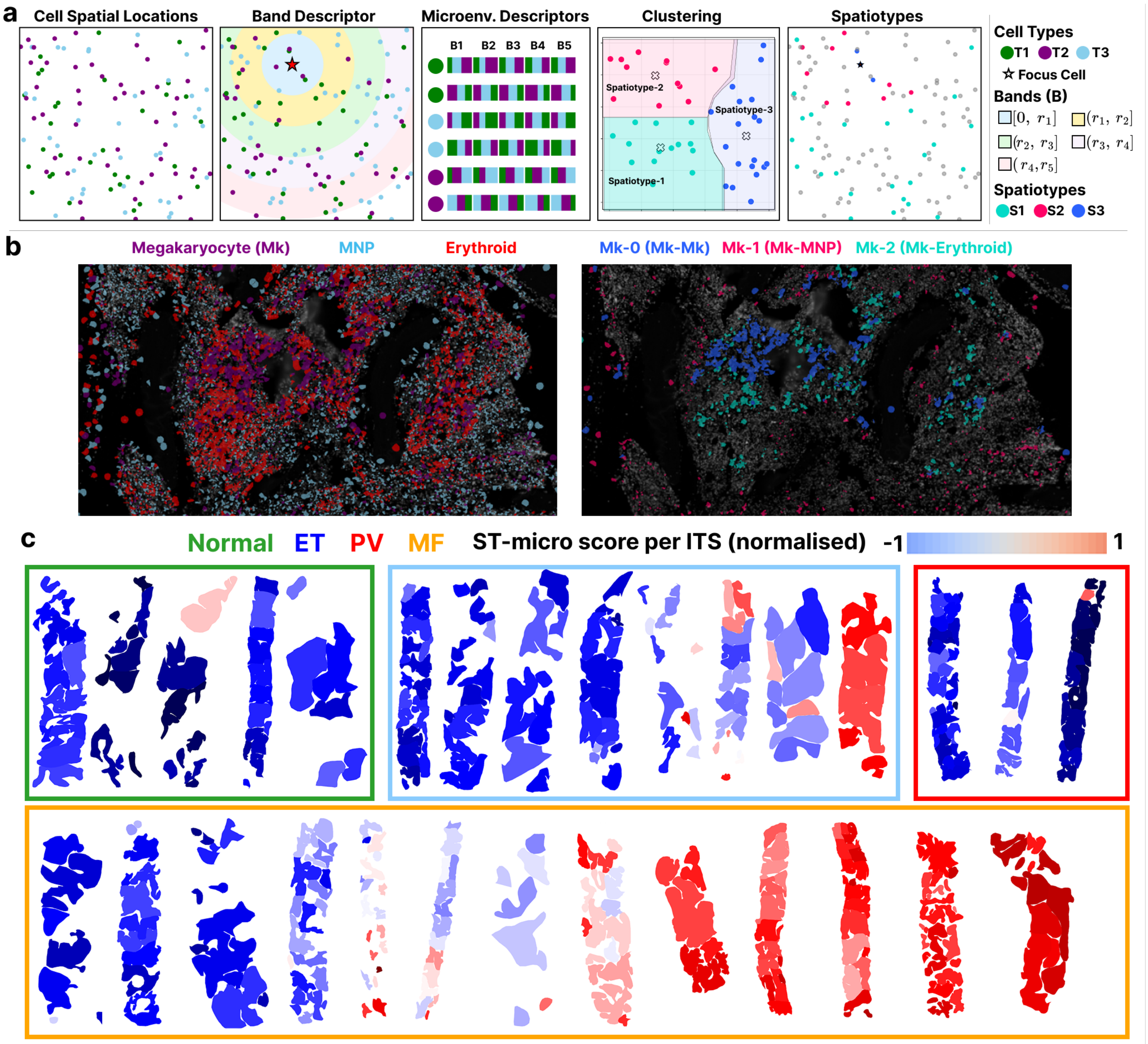


### Extended Data Figure 9: Identifying bone marrow microenvironmental signatures and characterising microenvironmental variation

**A:** Workflow to demonstrate characterisation of BM spatiotypes. The workflow takes cell (x, y) coordinates as input and computes the spatial cellular abundance surrounding each focus cell using five concentric bands covering a total radius of [0, 500] pixels. For each cell, a single vector is computed to represent the local spatial cellular composition. K-means clustering (k = 3) is then applied to these descriptors for each cell type to identify representative microenvironmental signatures, or spatiotypes. Each cell is subsequently assigned a refined label based on its local microenvironment or 'spatiotype'. For example, a megakaryocyte assigned a megakaryocyte-0, megakaryocyte-1, or megakaryocyte-2 spatiotype assignment, representing the distinct patterns of spatial abundance of cells surrounding megakaryocytes. **B:** Xenium ST DAPI images to demonstrate megakaryocyte cellular spatiotypes. To the right, the three megakaryocyte spatiotypes are highlighted (megakaryocyte-0 in blue, megakaryocyte-1 in red and megakaryocyte-2 in aquamarine). Megakaryocyte-0 is characterised by local enrichment of megakaryocytes, shown in the left image (purple). Megakaryocyte-2 (aquamarine) is characterised by local enrichment of erythroid cells, demonstrated in the left image (erythroids are highlighted in red). **C:** Bone marrow trephine heatmaps to show the overall ST-micro score per intertrabecular space (ITS) across the cohort, including normal samples (green border), ET samples (blue border), PV samples (red border) and MF samples (orange border). Regions highlighted in dark red (towards 1) infer a high ST-micro score and MF-like microenvironment, while those in dark blue (towards -1) are enriched for normal-like microenvironment. Scores were derived from a ranking-based multiple-instance-learning (MIL) model trained to rank instances from MF samples higher than those from normal samples (see Methods).

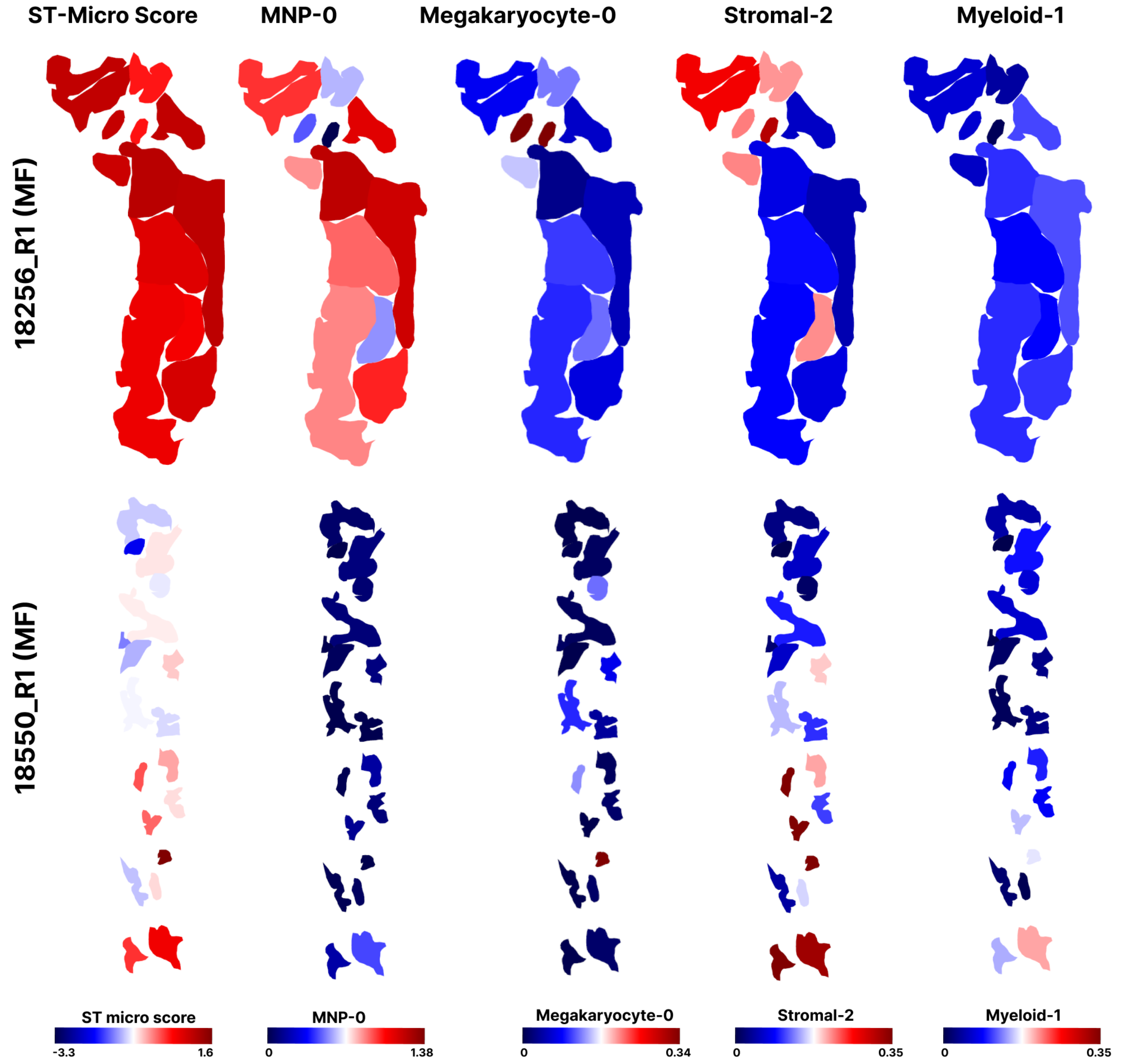


**Extended Data Figure 10: Machine learning (ML)-based detection of microenvironmental heterogeneity**

Bone marrow trephine (BMT) heatmaps to show: (left) overall mean ST-micro score per intertrabecular space (ITS) and the enrichment of MNP-0, megakaryocyte-0, stromal-2 and myeloid-1 spatiotypes seen to be driving the overall MF-like ST-micro score in Figure 8E. Samples 18256_R1 and 18550_R1 are both BMTs from patients with MF.
