## Supplementary Figures for "Redefining the topology of the human bone marrow using augmented spatial transcriptomic analysis"

| Sample ID | Age | Sex | Diagnosis | Mutation Status |
|---|---|---|---|---|
| 16818_R1 | 40-49 | F | ET | JAK2 |
| 16818_R2 | 30-39 | M | ET | JAK2 |
| 16832_R2 | 60-69 | F | ET | JAK2 |
| 18552_R3 | 40-49 | M | ET | JAK2 |
| 18599_R3 | 20-29 | M | ET | CALR |
| 18606_R1 | 30-39 | F | ET | CALR |
| 18606_R2 | 40-49 | M | ET | JAK2 |
| 18669_R3 | 40-49 | M | ET | TN |
| 10693_R2 | 50-59 | F | PV | JAK2 |
| 18256_R2 | 60-63 | M | PV | JAK2 |
| 18286_R2 | 60-67 | F | PV | JAK2 |
| 18256_R3 | 60-69 | F | PrePMF | TN |
| 18286_R3 | 40-49 | M | PrePMF | TN |
| 16818_R3 | 50-59 | F | PrePMF | JAK2 |
| 10693_R1 | 70-79 | F | MF | JAK2 |
| 16832_R1 | 50-59 | F | MF | JAK2 |
| 18256_R1 | 70-79 | F | MF | JAK2 |
| 18286_R1 | 49-49 | M | MF | JAK2 |
| 18550_R1 | 49-49 | M | MF | JAK2 |
| 18552_R2 | 30-39 | F | MF | JAK2 |
| 18599_R1 | 70-79 | M | MF | JAK2 |
| 18599_R2 | 70-79 | M | MF | CALR |
| 18612_R2 | 60-69 | F | MF | JAK2 |
| 18612_R3 | * | * | * | * |
| 18612_R1 | 50-59 | M | MF | CALR |
| 18612_R4 | * | * | * | * |
| 18669_R2 | 70-79 | M | MF | JAK2 |
| 18550_R2 | 40-49 | M | Normal | NA |
| 18550_R3 | 60-62 | M | Normal | NA |
| 18552_R1 | 50-59 | F | Normal | NA |
| 18606_R3 | 70-79 | F | Normal | NA |
| 18669_R1 | 30-39 | F | Normal | NA |

**Supplementary Figure 1: Cohort metadata**

Details of the study cohort, including age, sex, diagnosis and driver mutation. ET: essential thrombocythaemia; PV: polycythaemia vera; MF: myelofibrosis. TN indicates triple negative (i.e. no mutation detected in *JAK2, CALR* or *MPL*). *18612_R2 and 18612_R3 are from the same BMT bisected to facilitate tissue placement on the Xenium slide. 18612_R1 and 18612_R4 are from the same BMT (bisected).

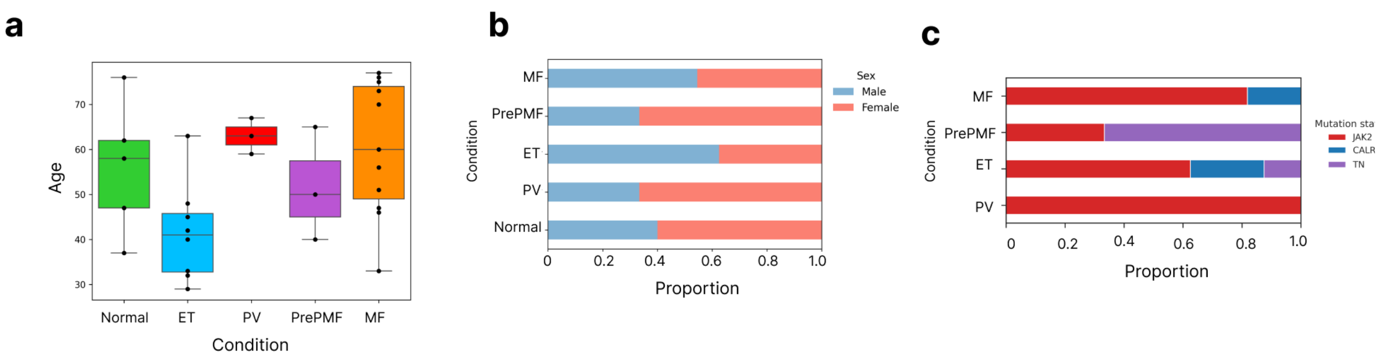


### Supplementary Figure 2: Description of cohort

**A:** Boxplot to show age distribution of the cohort by condition (normal, ET, PV, PrePMF and MF). **B:** Stacked bar plot to show the proportion of samples from each sex by condition. **C:** Stacked bar plot to show the proportion of samples with a key MPN-related driver mutation by condition.

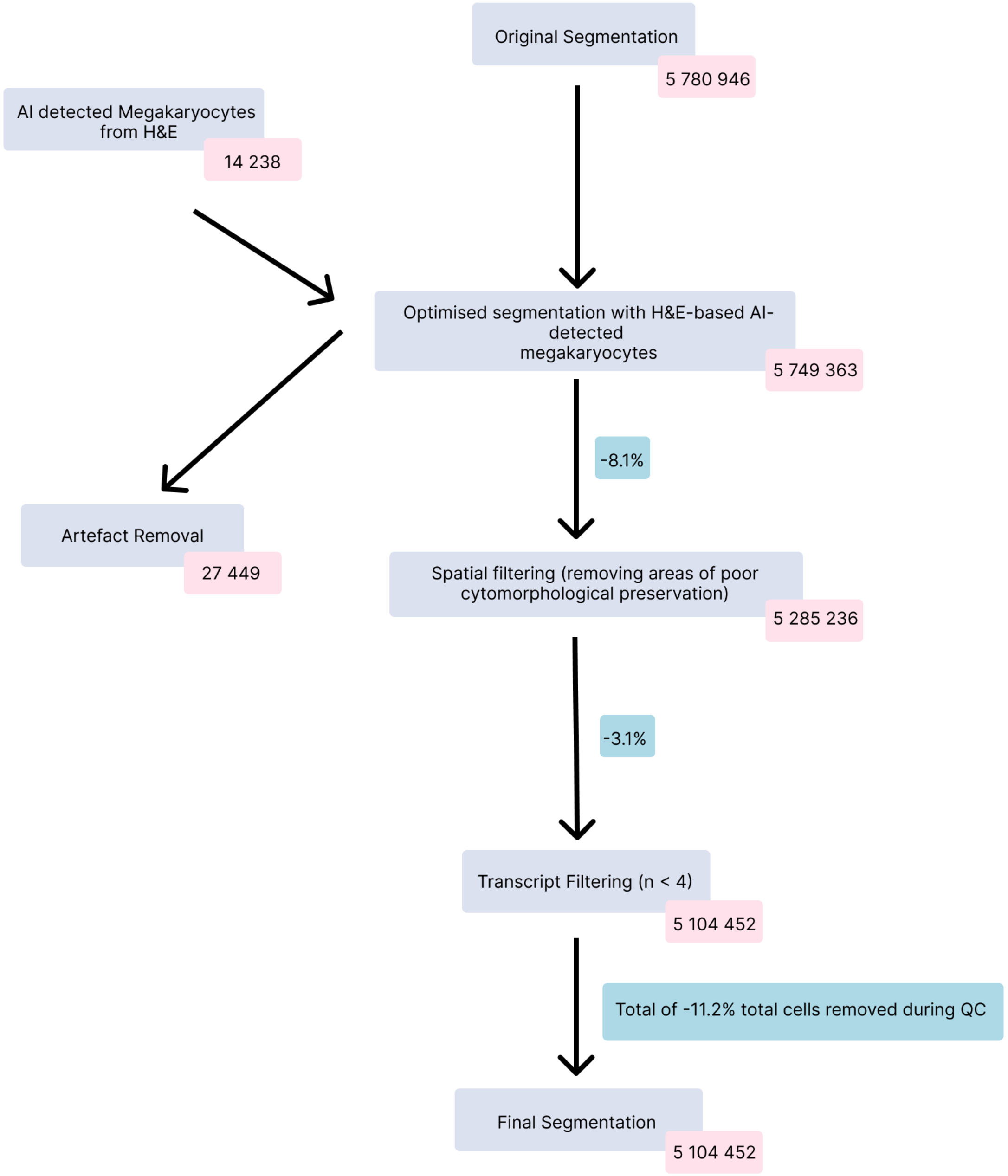


**Supplementary Figure 3: Data QC workflow**

Flowchart to show segmentation and filtering steps for quality control (QC) of Xenium bone marrow spatial transcriptomic (ST) data. Following Xenium segmentation, 5,780,946 cells were obtained from 30 BMT samples (from 30 different individuals). Following the application of our AI-based megakaryocyte detection algorithm, 14,238 megakaryocytes were identified and when these were overlayed (see Methods) onto the ST data 31,583 cells were removed (through, for example cell merging). After pathologist-led annotation, cells outside of regions of cytomorphological preservation were then removed (n=464,127 cells, 8.1%). After this, remaining cells with fewer than 4 transcripts were removed (n=180,784 cells). A total of 5,104,452 cells were taken forward for cell typing and downstream analysis.

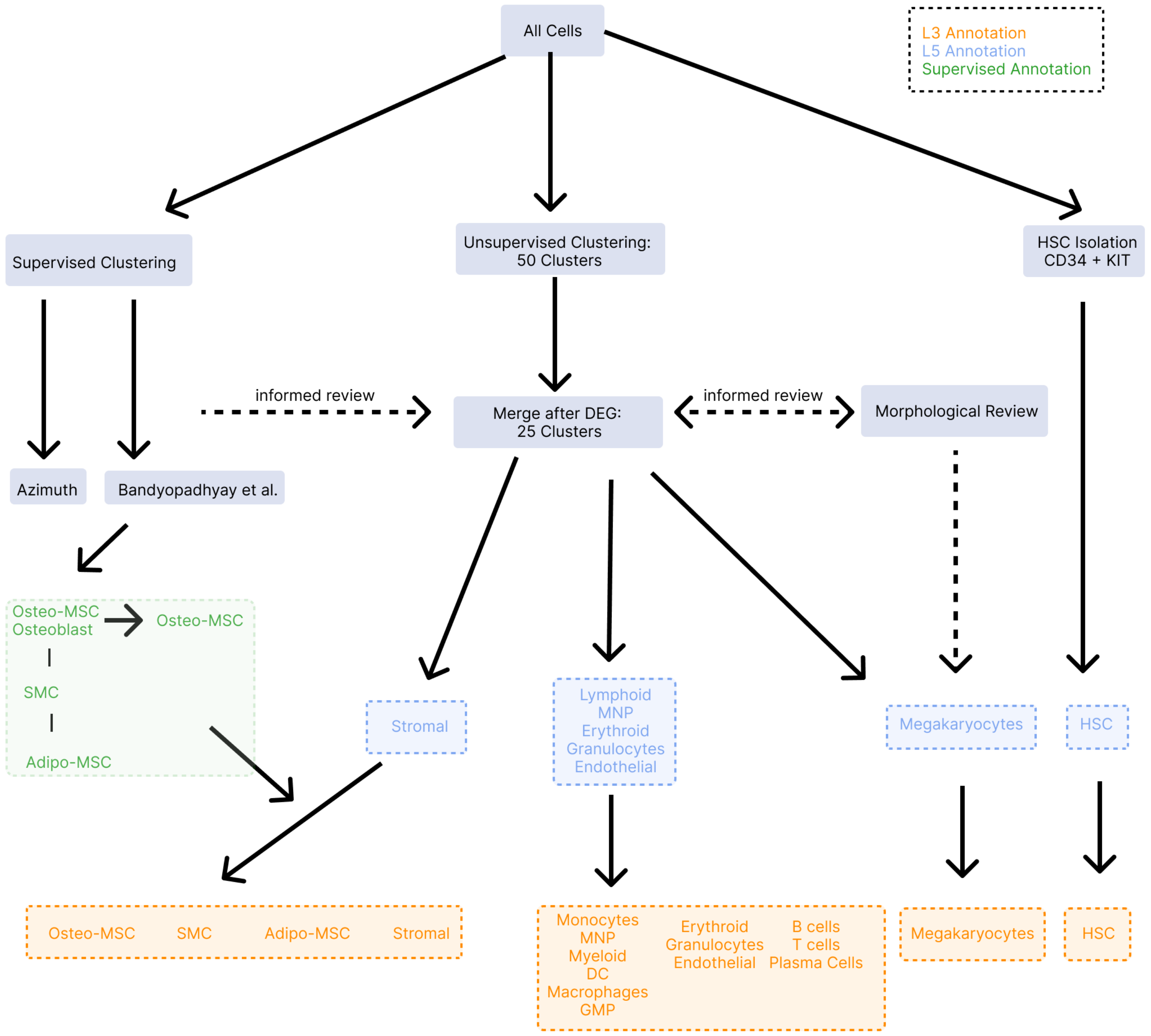


**Supplementary Figure 4: Cell type annotation workflow**
Following upstream QC, 5,104,442 cells were taken forwards for cell type annotation. We manually filtered HSCs as those co-expressing CD34 and c-kit (n = 21,680 cells, 0.4%). With the remaining cells we then performed unsupervised clustering with 20 principal components. With clustering-detected cell groups we performed differential expression analysis (using *Seurat FindAllMarkers*) to identify cluster-defining markers, and merged cells with the same assignment. We then reviewed the cell assignment by overlying the ST-detected cells onto the H&E-stained tissue. We performed supervised annotation using *SingleR* with external scRNA-seq reference datasets (Extended Data Fig. 3E) to support and confirm lineage assignment[1,2]. We also used supervised annotation with the stromal-enriched external reference scRNA-seq dataset (Bandyopadhyay et al.[2]) to identify stromal subtypes (osteo-MSCs, adipo-MSCs, stromal and smooth muscle cells (SMCs). We also identified a cell group only defined by expression of CD69 comprising only 0.09% (n=4,524) cells of all cells, which we removed from all downstream analysis due to lack of confidence in lineage assignment.

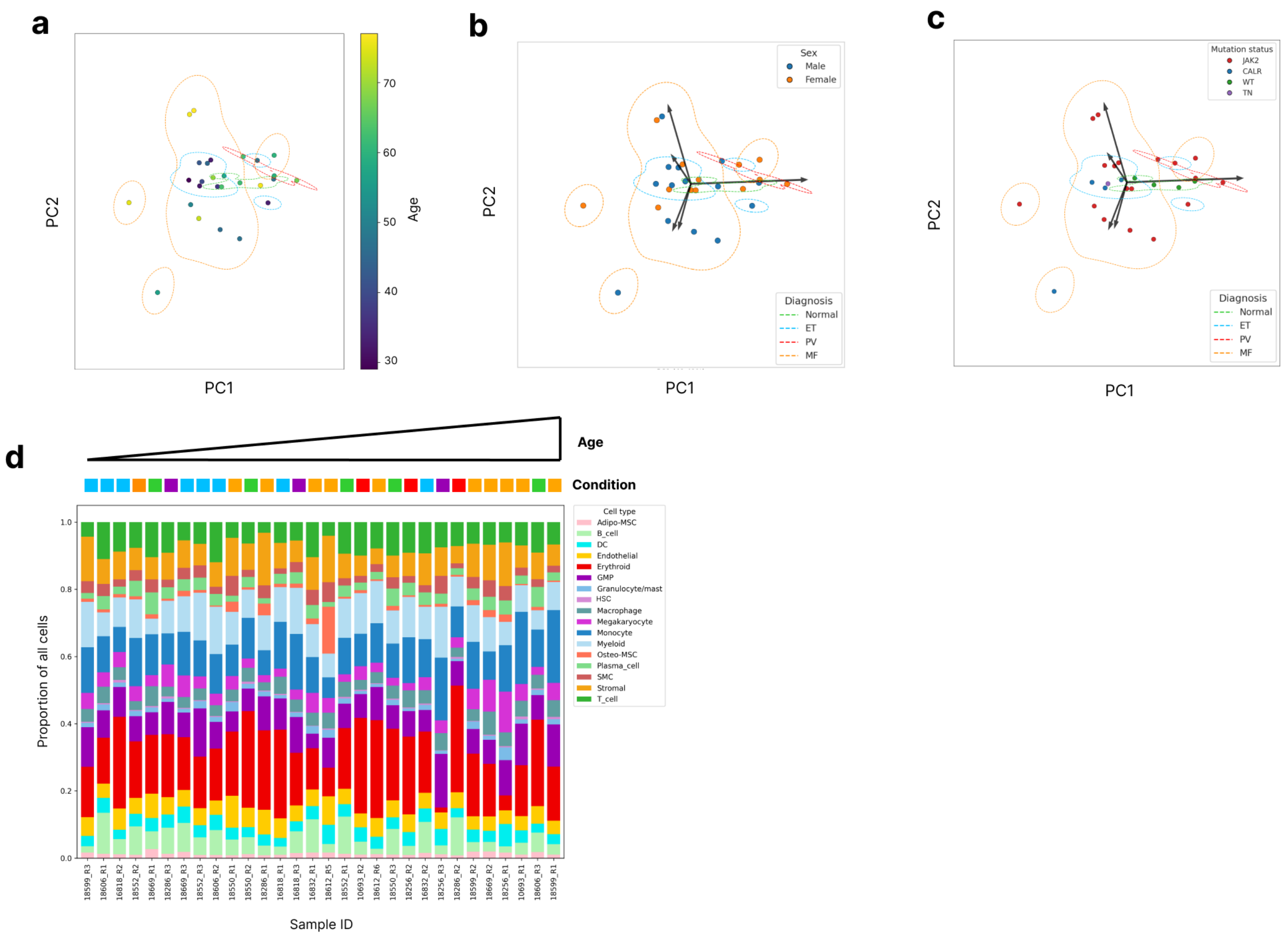


### Supplementary Figure 5: Cohort details

**A:** PCA plot to show the variance (PC1 & PC2) of differential abundance of BM cells types. Points represent a sample coloured by age (see scale right). **B:** PCA plot to show the variance of differential cellular abundance. Points represent a sample coloured by sex. **C:** PCA plot to show the variance of differential abundance of all cells. Points represent a sample coloured by mutation status. **D:** Stacked bar plot to show proportions of each cell type per sample ordered from youngest (left) to oldest (right). Colour key at the top indicates condition (normal – green; ET – blue; PV – red; MF - orange).

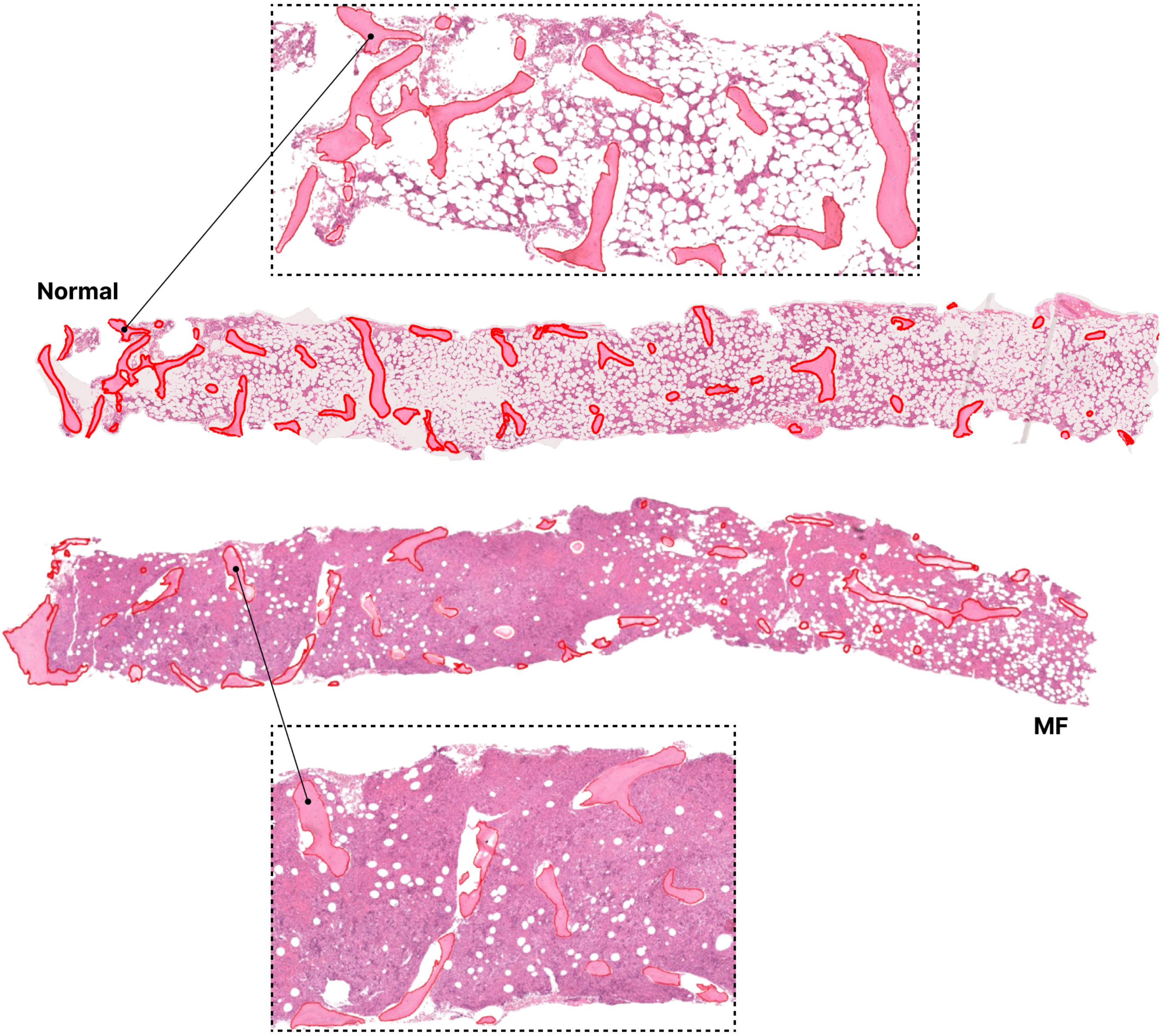


**Supplementary Figure 6: Bone segmentation**
Images show the output of the bone segmentation AI-based algorithm applied as previously described by Ryou *et al*.[3]. This algorithm detects bones in normal samples (top) and below (MF). The red line indicates the bone surface as detected by the AI-based algorithm.

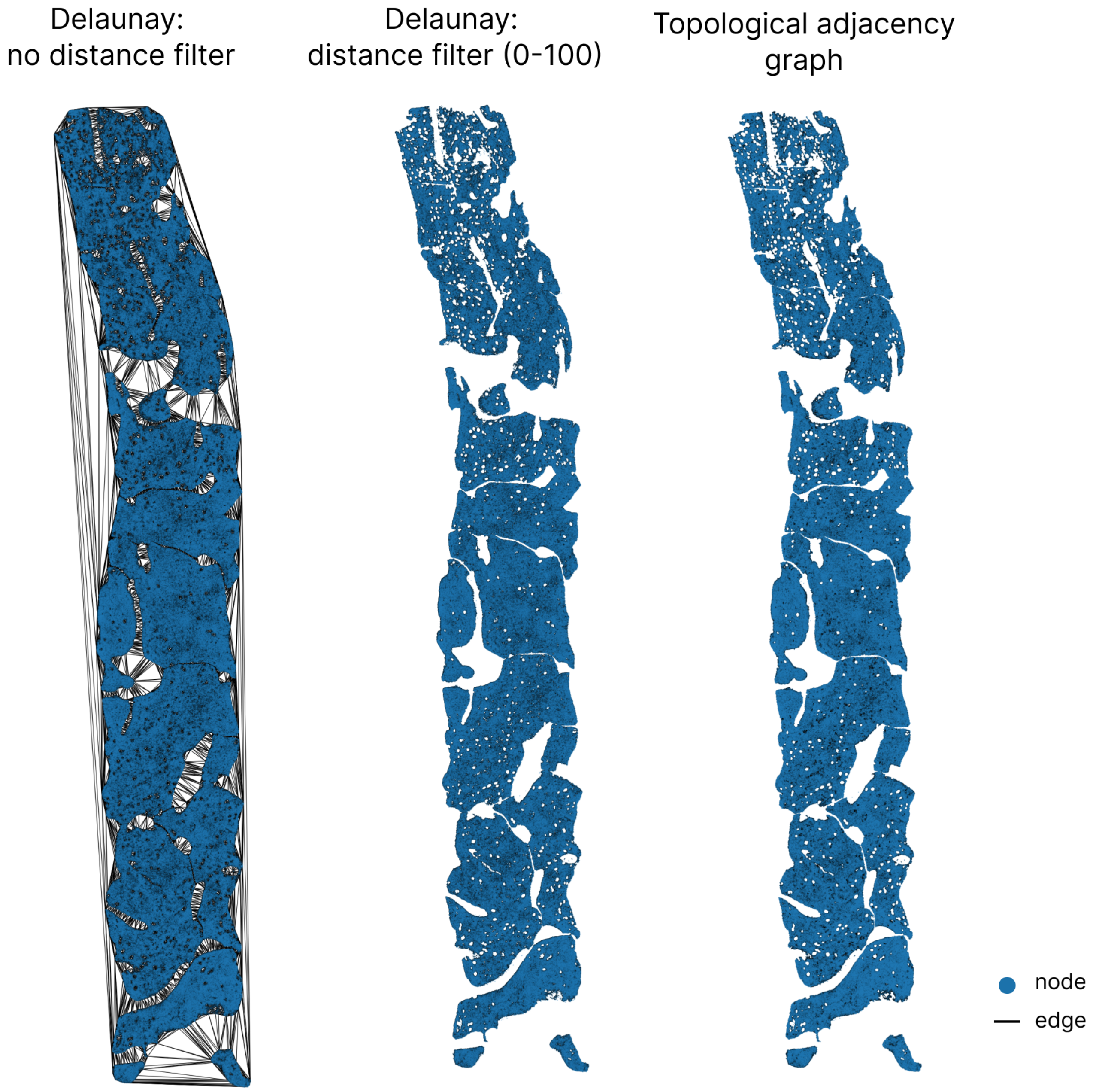


**Supplementary Figure 7: Spatial graph construction**

Whole BMT images to show spatial graph construction using both Delaunay (both with and without distance filter) and topological adjacency graph-based method. (Left) BMT to show Delaunay spatial graph without a distance filter. Without the distance filter, spurious connections (edges) are created between cells across spaces (bone and adipose tissue). (Middle) Delaunay graph construction with a 100-pixel distance filter. (Right). Topological adjacency graph construction. The Delaunay graph construction using a distance filter and the topological adjacency graph construction approach both limit inappropriate edge connections between cells not spatially local to each other (e.g. across bone, end-to-end connections).

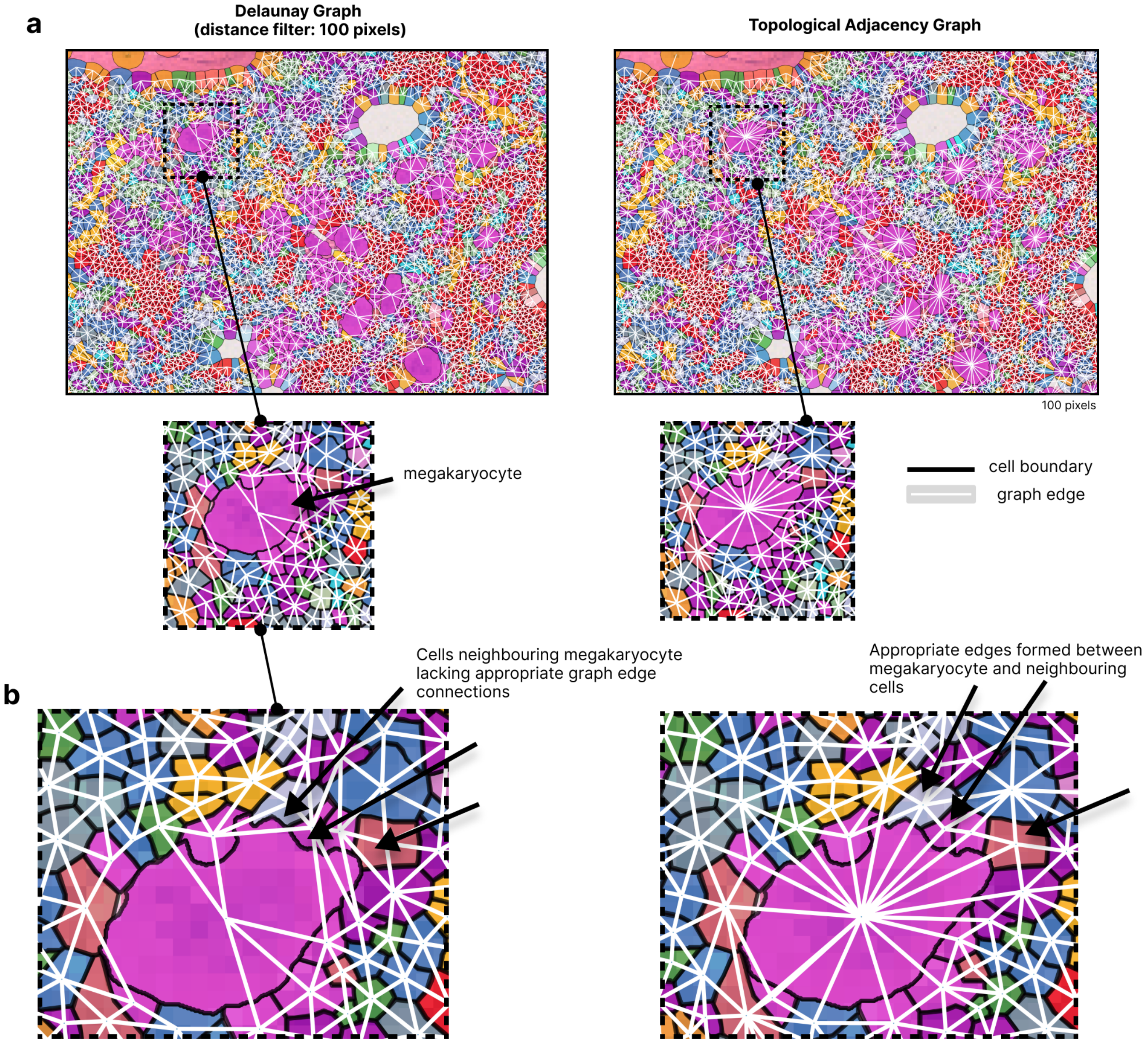


**Extended Figure 8: Spatial graph construction**

**A:** Image to show spatial graph construction using a Delaunay-based method (left, with a 100 pixel distance filter) and a topological adjacency-based method (right). Delaunay spatial graph construction performs triangulation that forbids the intersection of triangles. This means that cells neighbouring irregularly-shaped cells (e.g. megakaryocytes) are not consistently captured. Topological adjacency-based graph construction allows for the intersection of edges and does not require a distance threshold. This means that cells neighbouring those of varying sizes are more consistently captured. Topological adjacency-based graph construction also prevents graph construction between cells that are not touching across large spaces (e.g. across bony trabeculae). This also preserves fat spaces, preventing graph edges across mature adipocytes. The inset images at higher power demonstrate the lack of appropriate edges between the megakaryocytes and neighbouring cells with the Delaunay graph construction. **B:** (Left) Tile image to show Delauny spatial graph construction around a large, irregularly shaped cell (megakaryocyte) with a 100 pixel distance filter. Here, edges (connections) are lacking between the megakaryocyte and some neighbouring cells. With the topological adjacency-based graph construction method (right) appropriate edges are created between cell neighbours.

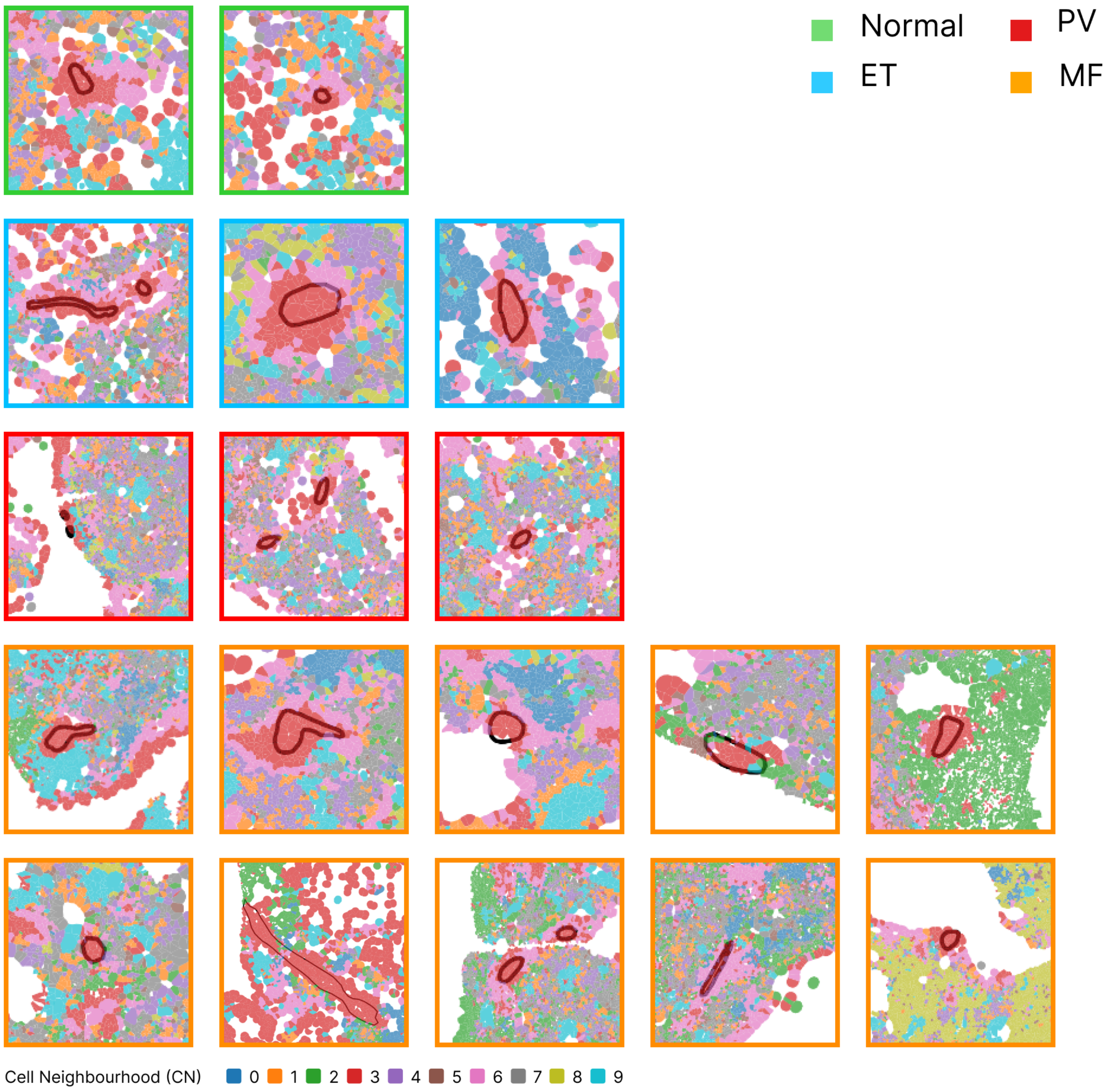


**Supplementary Data Figure 9: Arteriole location**
Tile images to show the location of arterioles (annotated) in regions of cell neighbourhood 3 (CN3) (red). Cells are coloured by CN assignment. Tile images are bounded by border colours to indicate condition with green = normal, blue = ET, red = PV and orange = MF.

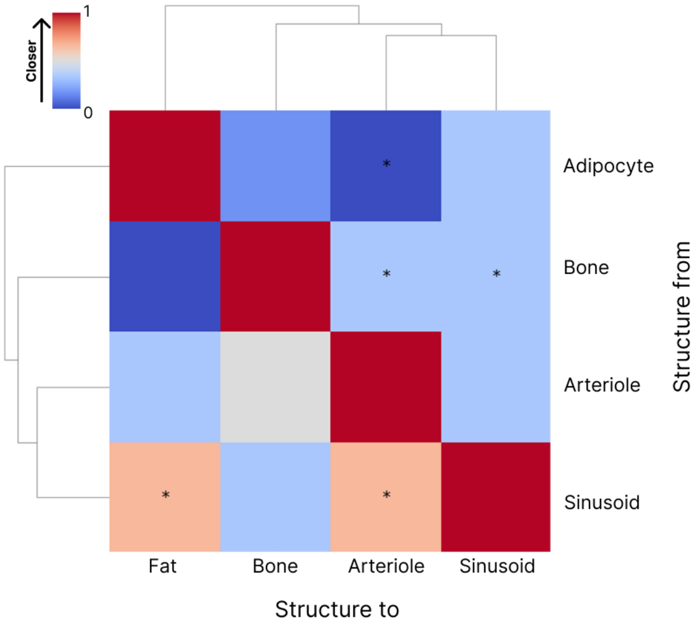


**Supplementary Figure 10: Structure-structure proximity analysis**
Heatmap to show the difference (delta) in rank proximity of architectural features (bone, adipocytes (fat), arterioles, sinusoids) to each other in normal samples. Towards red infers proximity, while towards blue indicates a more distal distribution (*denotes p-value <0.05, Wilcoxon rank sum test).

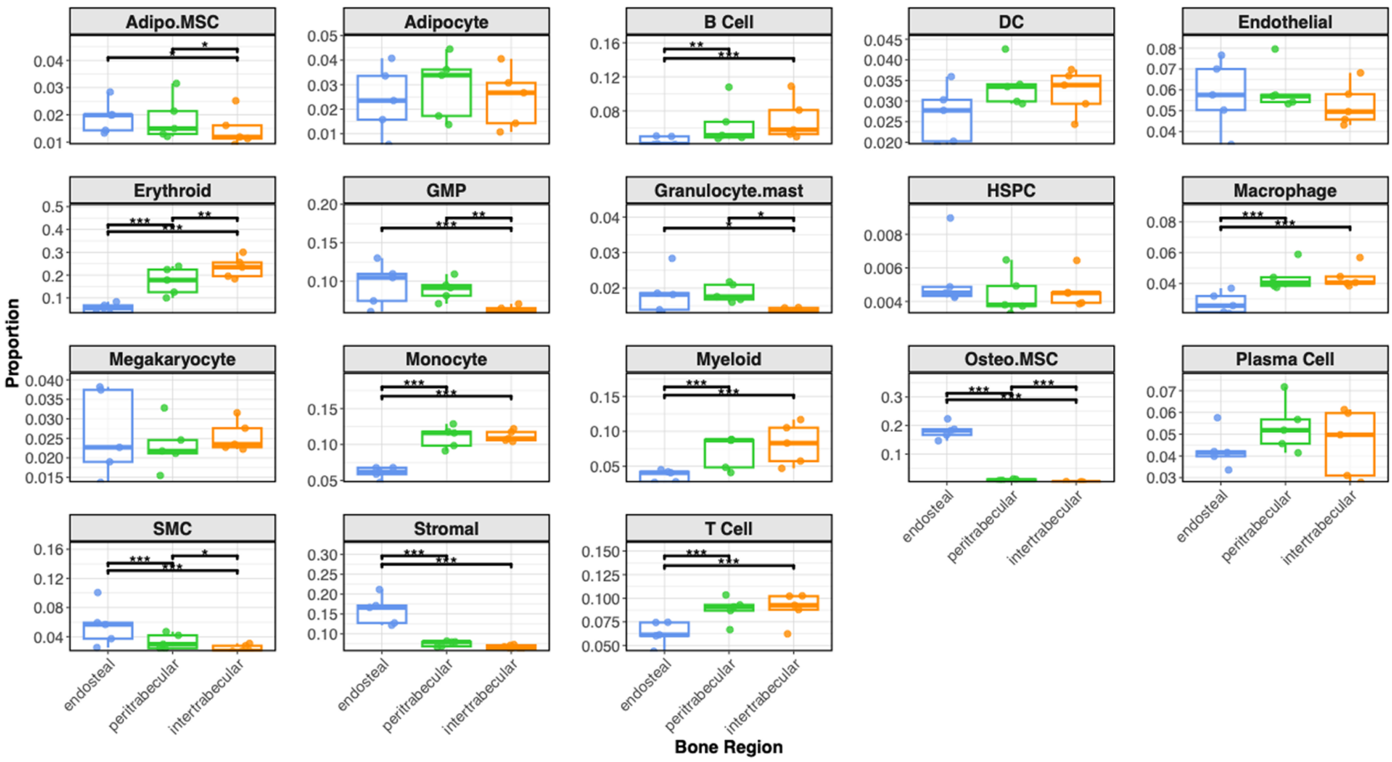


**Supplementary Data Figure 11: Differential cellular abundance across bone marrow regions**
Boxplots to show the differential abundance of cell types across supervised bone marrow niche spaces (endosteal/peritrabecular/intertrabecular – as per Figure 3G) in normal samples (n=5) (generalized linear model, *edgeR*).

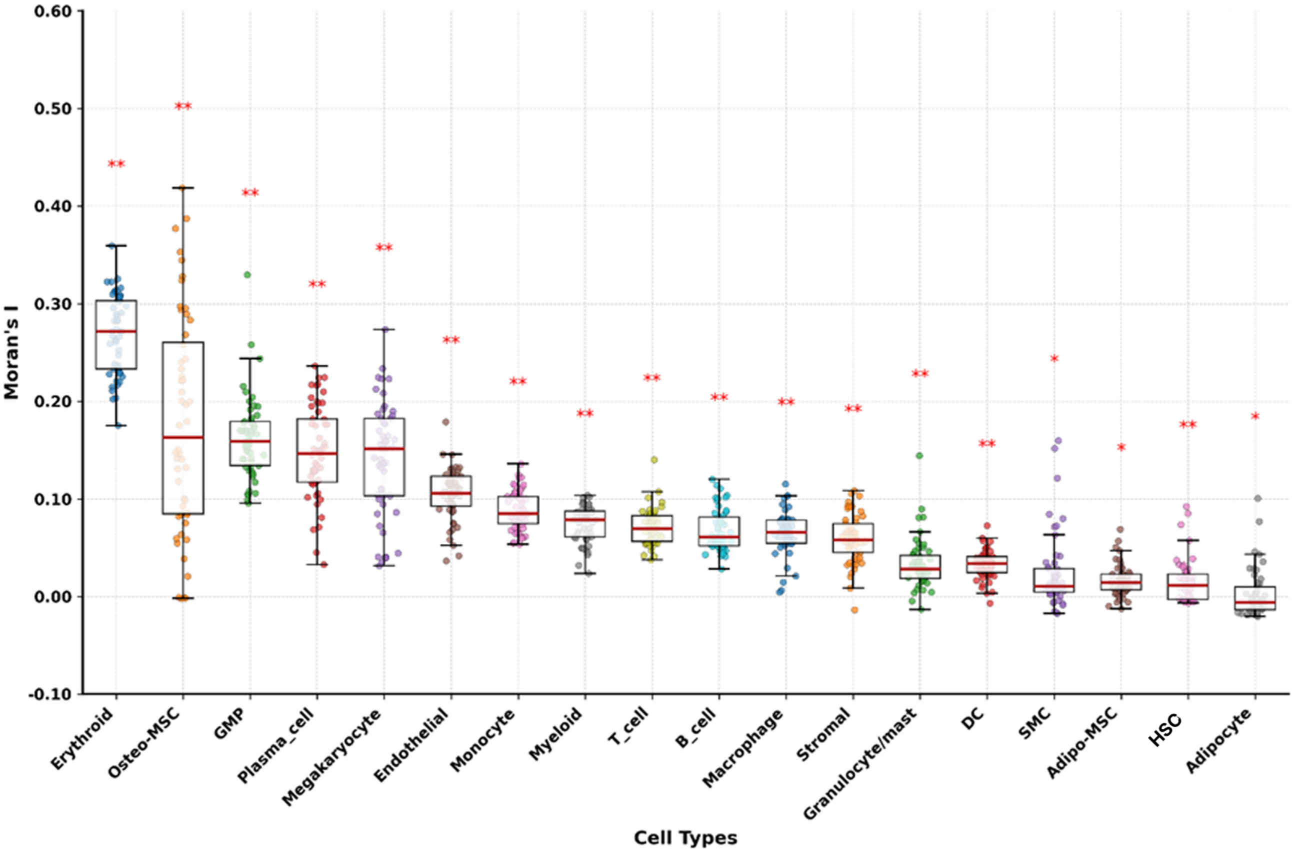


**Supplementary Figure 12: Cellular spatial autocorrelation**

Each boxplot shows the Moran's I values per cell type per intertrabecular space (ITS) region in normal samples (n = 5). Cell types are ordered by decreasing mean Moran's I. Red asterisks indicate statistical significance estimated using 2* (median p-value) as conservative estimates. (* $p<0.05$, ** $p<0.01$, *** $p<0.001$).

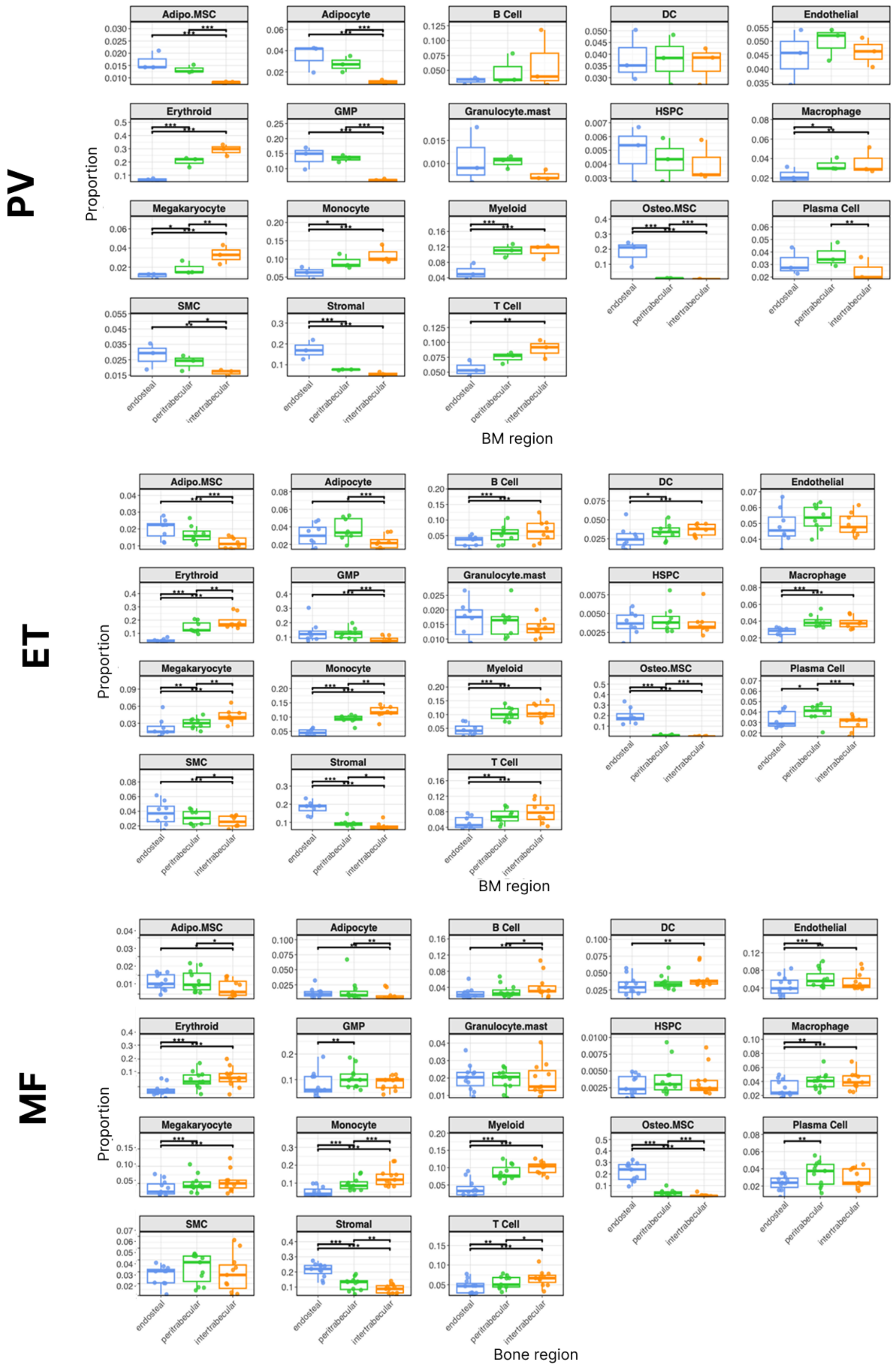


**Supplementary Figure 13: Cellular abundance across bone marrow regions in MPN**

Boxplots to show the differential abundance of cell types across bone marrow regions (endosteal space, peritrabecular space and intertrabecular/central space as per 3G) in PV, ET and MF BMTs (negative generalised linear model, *edgeR*).

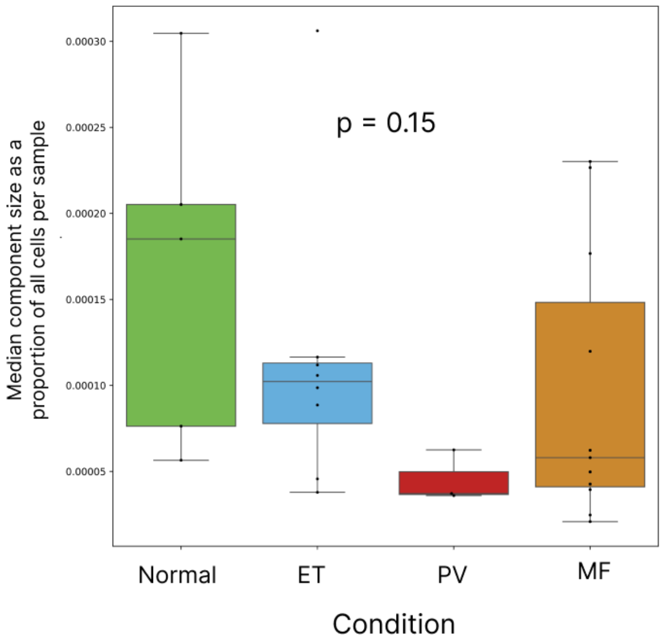


**Supplementary Figure 14: Cell neighbourhoods**

Boxplot to show the median cell neighbourhood (CN) component size as a proportion of all cells per sample across conditions (Mann-Whitney U test).

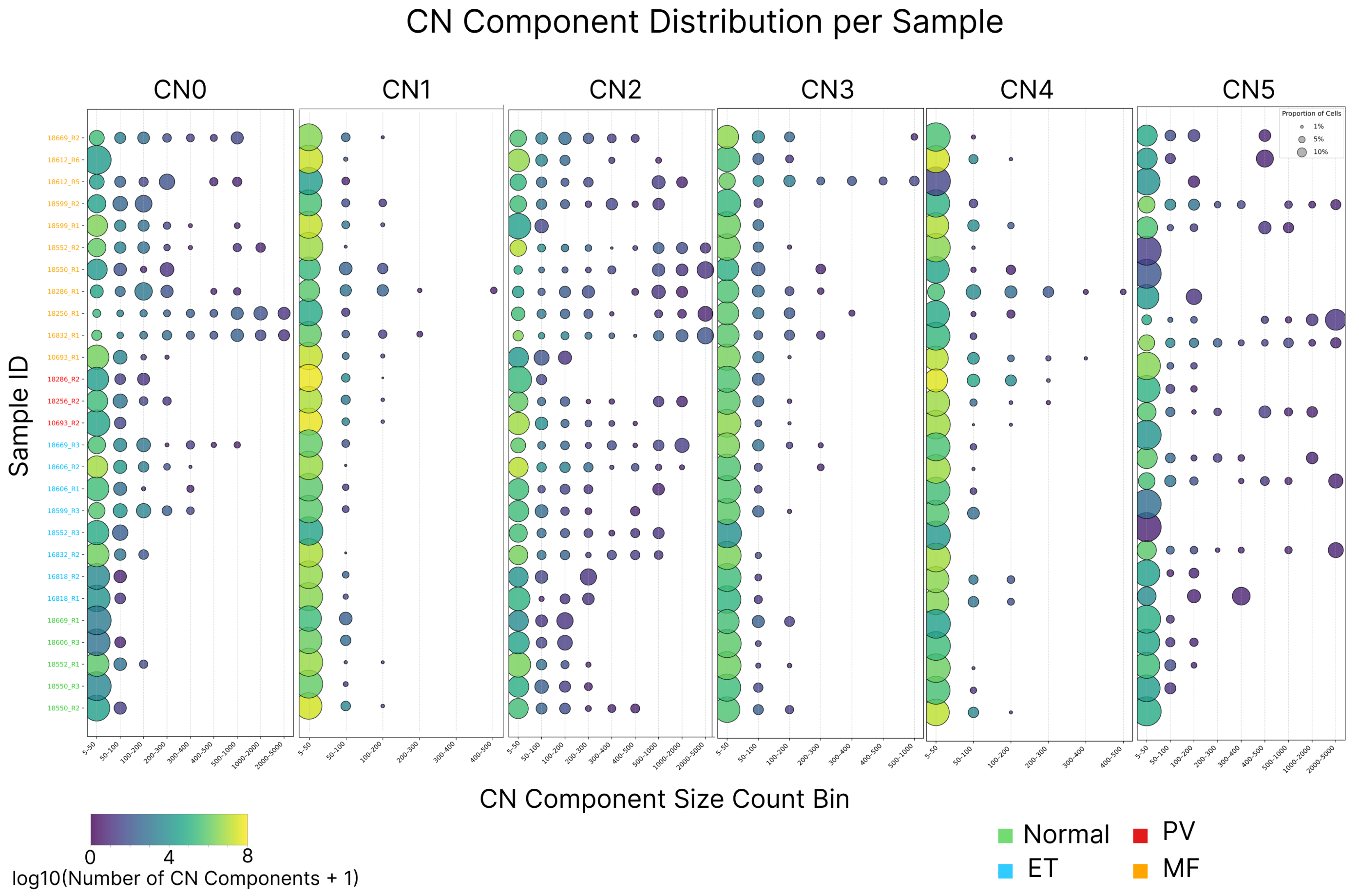


**Supplementary Figure 15A: Cell neighbourhood component size (CN0-CN5)**

Dot plots to show cell neighbourhood (CN) component size and size distribution per sample. The y-axis shows samples with sample ID coloured/grouped by condition. The x-axis shows the component bin size (where size is number of cells in each component). Dot size indicates the proportion of all cells in that sample within that particular bin size. Dot colour indicates the number (log10+1) of discrete CN components in per size 'bin' (low – purple, high – yellow, see colour scale bottom left).

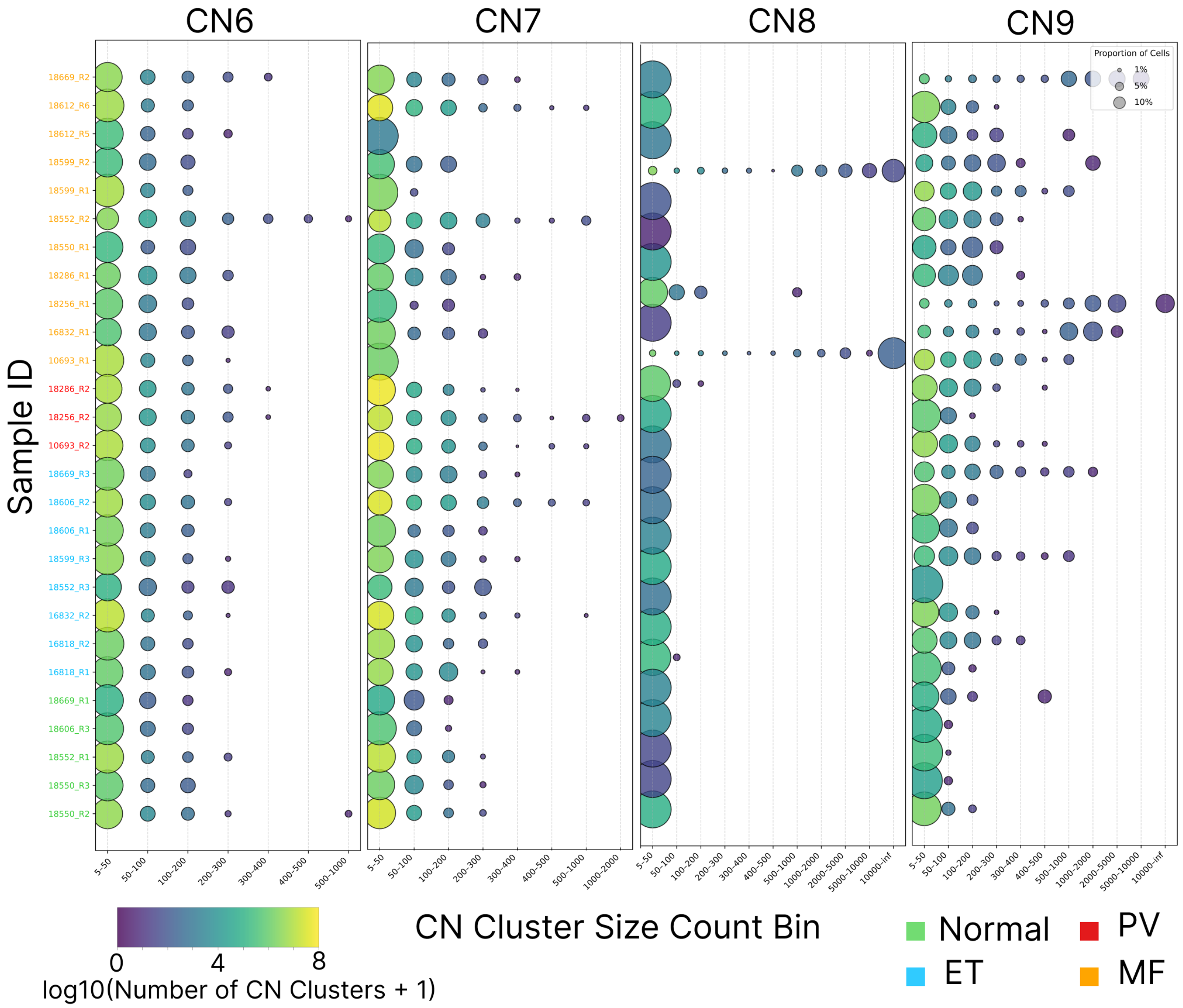


**Supplementary Figure 15B: Cell neighbourhood component size (CN6-9)**

Dot plots to show cell neighbourhood (CN) component size and size distribution per sample. The y-axis shows samples with sample ID coloured/grouped by condition. The x-axis shows the component bin size (where size is number of cells in each component). Dot size indicates the proportion of all cells in that sample within that particular bin size. Dot colour indicates the number (log10+1) of discrete CN components in per size 'bin' (low – purple, high – yellow, see colour scale bottom left).

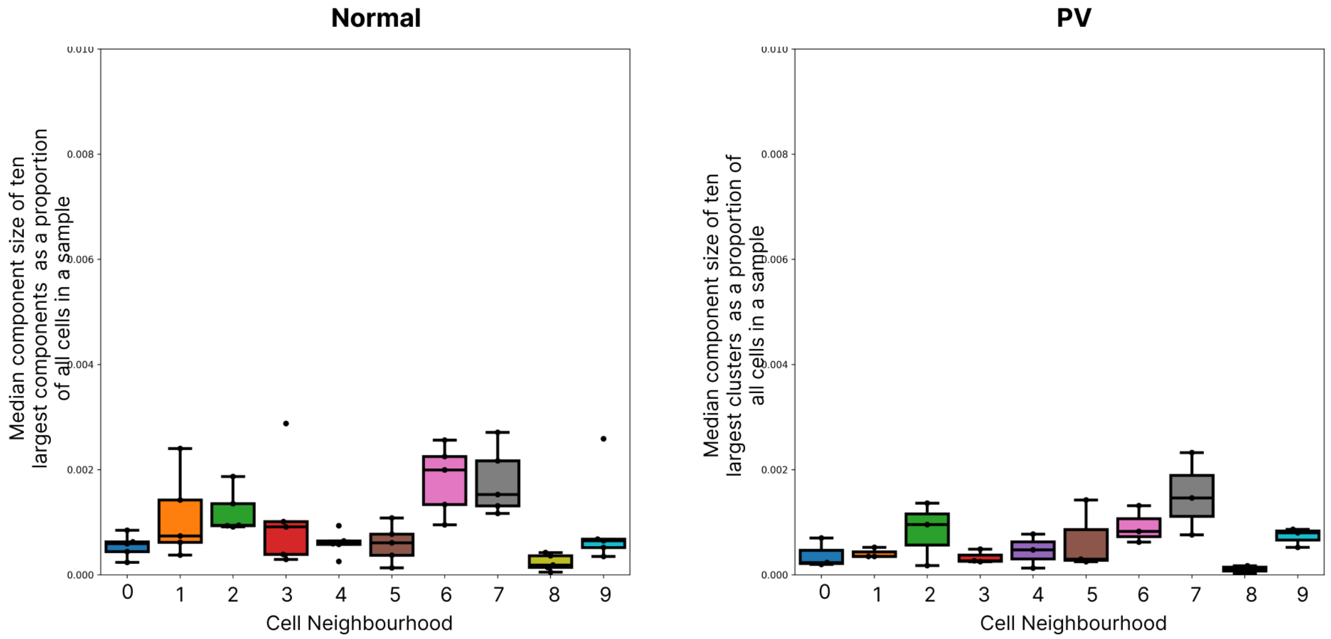


**Supplementary Figure 16: Cell neighbourhood component size in normal samples and PV**
Boxplot to show the median cell neighbourhood (CN) component size as a proportion of all cells in normal and PV samples (Wilcoxon signed-ranked test, no significant differences between any CNs).

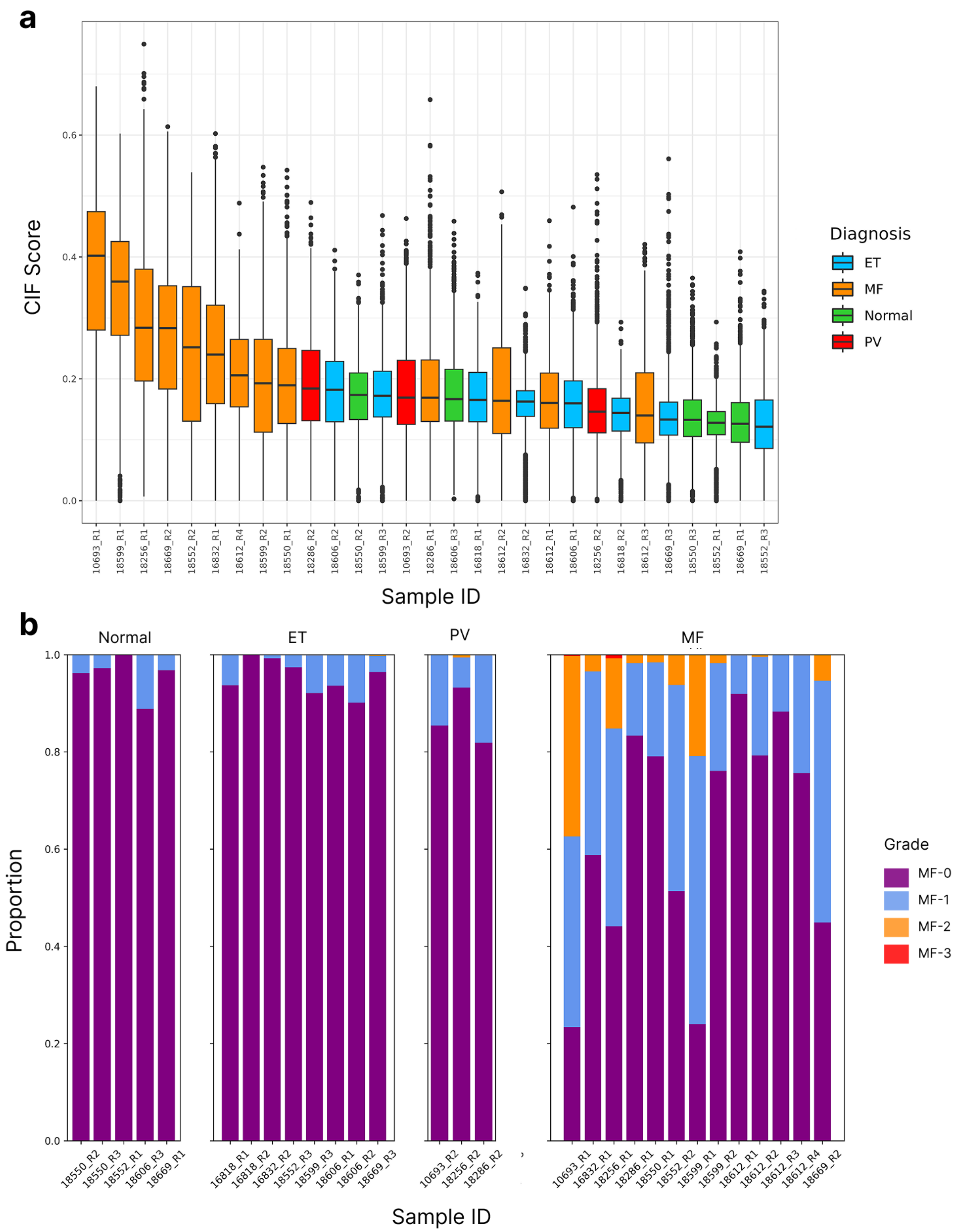


**Supplementary Figure 17: Continuous index of Fibrosis (CIF)**
**A:** Boxplot to show the CIF scores per sample by condition. The CIF score per 512x512 image was plotted per sample. Boxplots are coloured by condition. **B:** Stacked barplot to show the distribution of fibrosis scores per sample aligned to WHO fibrosis grades (WHO grade 0-3).

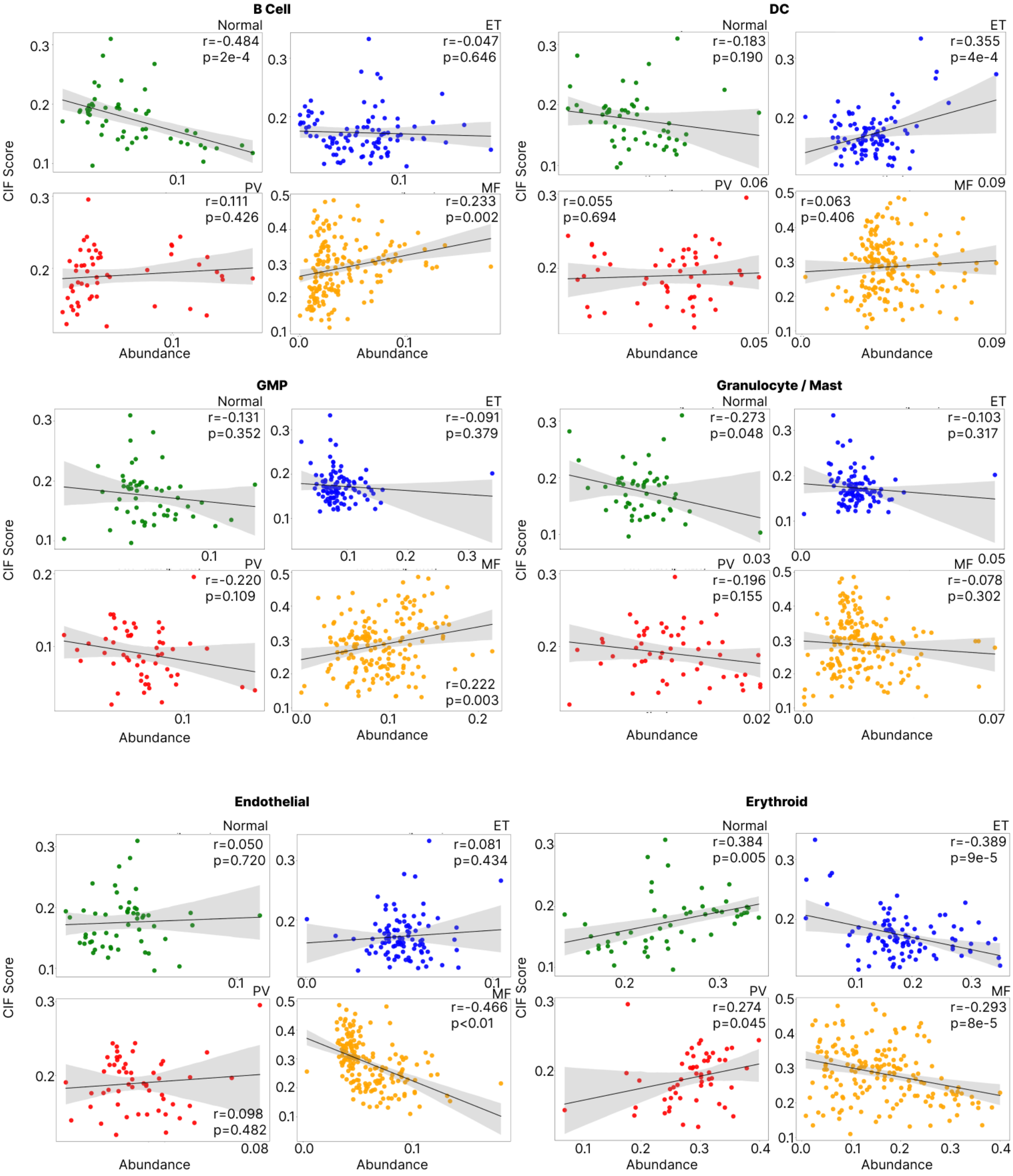


**Supplementary Figure 18A: Microenvironmental cellular abundance and fibrosis (CIF)**

Scatter plots to show correlation between the differential abundance of cell types (proportion of cell types per intertrabecular space, ITS) versus CIF (mean tile-level value per ITS) in normal, ET, PV and MF samples (Pearson's correlation).

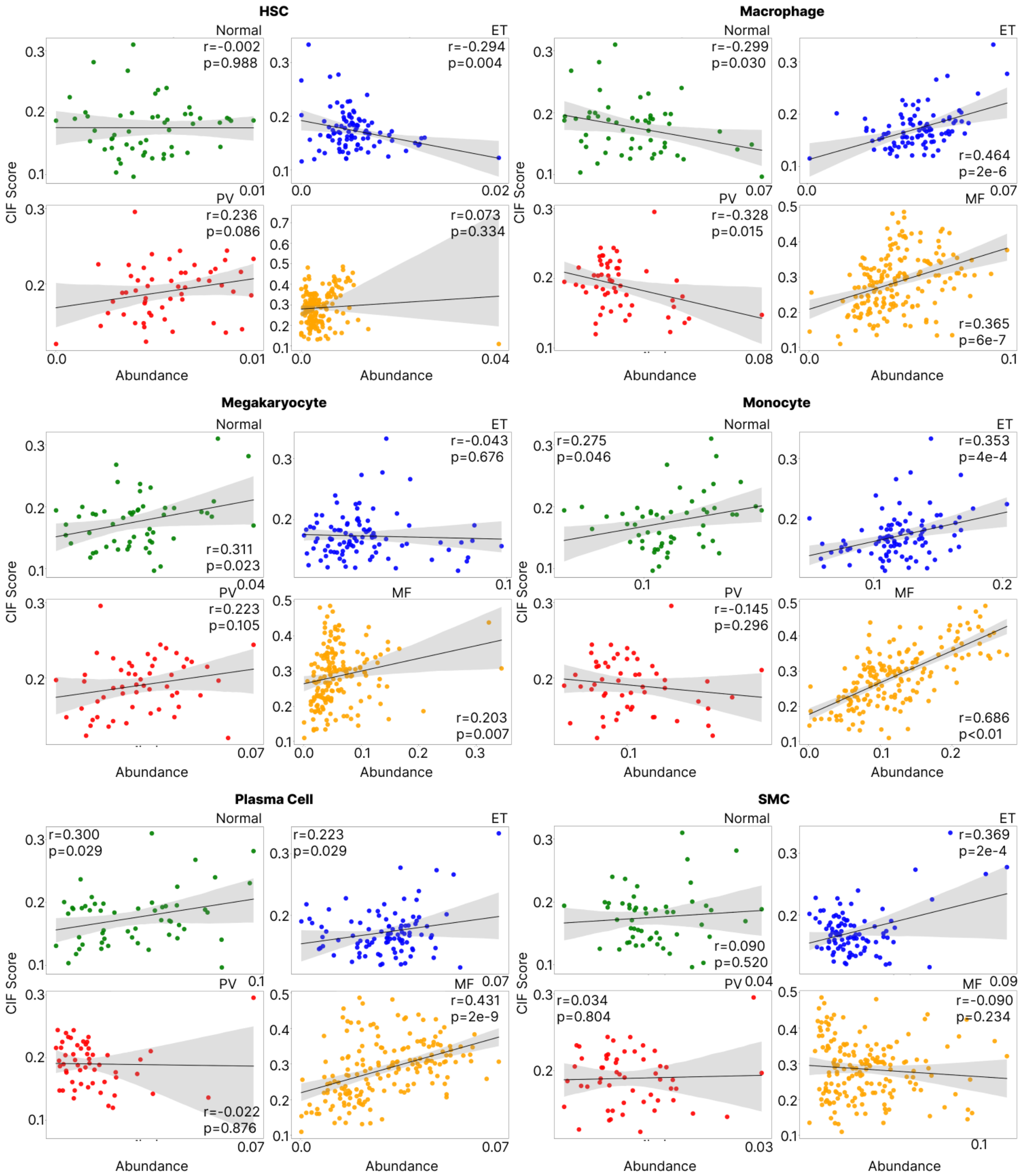


**Supplementary Figure 18B: Microenvironmental cellular abundance and fibrosis (CIF)**

Scatter plots to show correlation between the differential abundance of cell types (proportion of cell types per intertrabecular space, ITS) versus CIF (mean tile-level value per ITS) in normal, ET, PV and MF samples (Pearson's correlation).

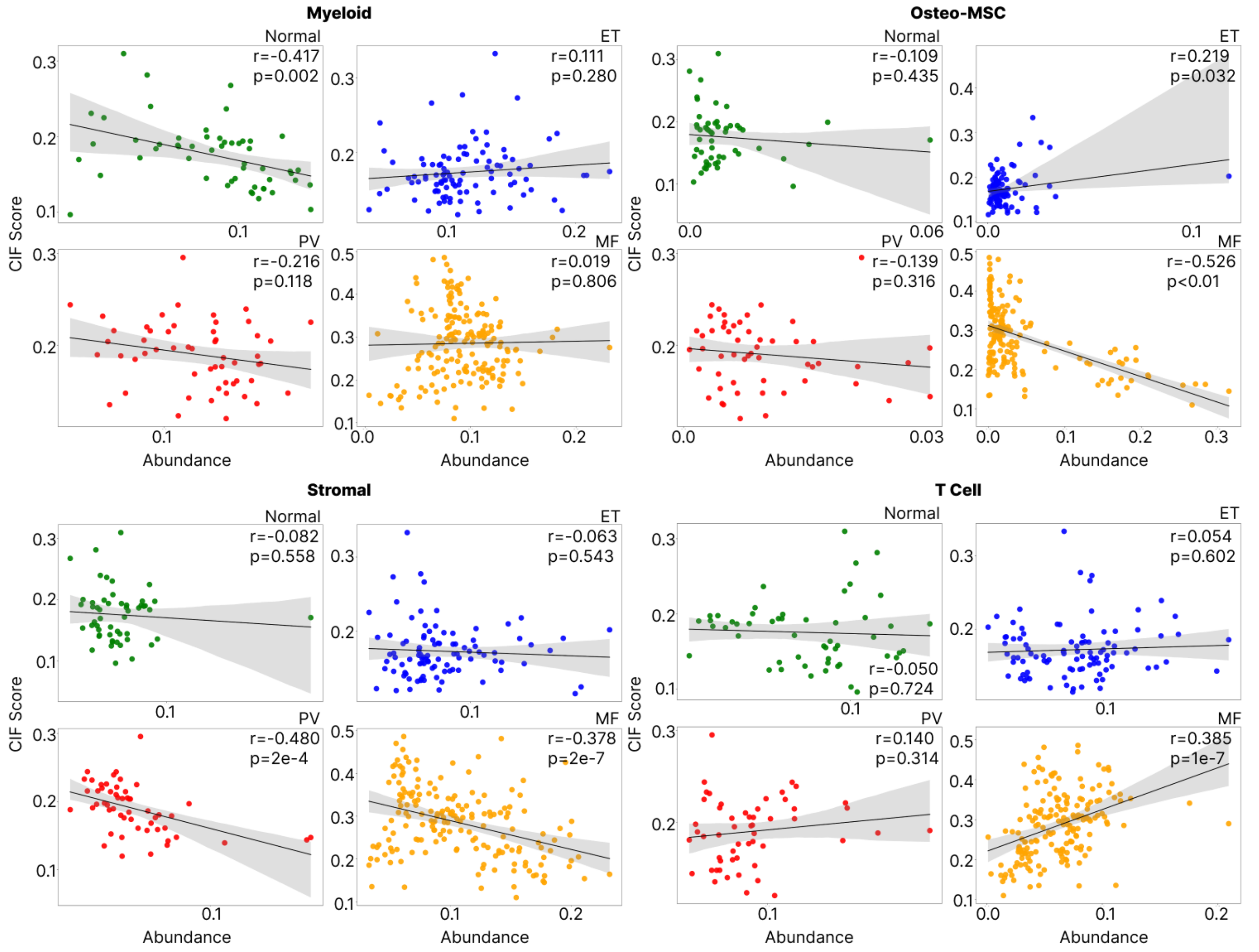


**Supplementary Figure 18C: Microenvironmental cellular abundance and fibrosis (CIF)**

Scatter plots to show correlation between the differential abundance of cell types (proportion of cell types per intertrabecular space, ITS) versus CIF (mean tile-level value per ITS) in normal, ET, PV and MF samples (Pearson's correlation).

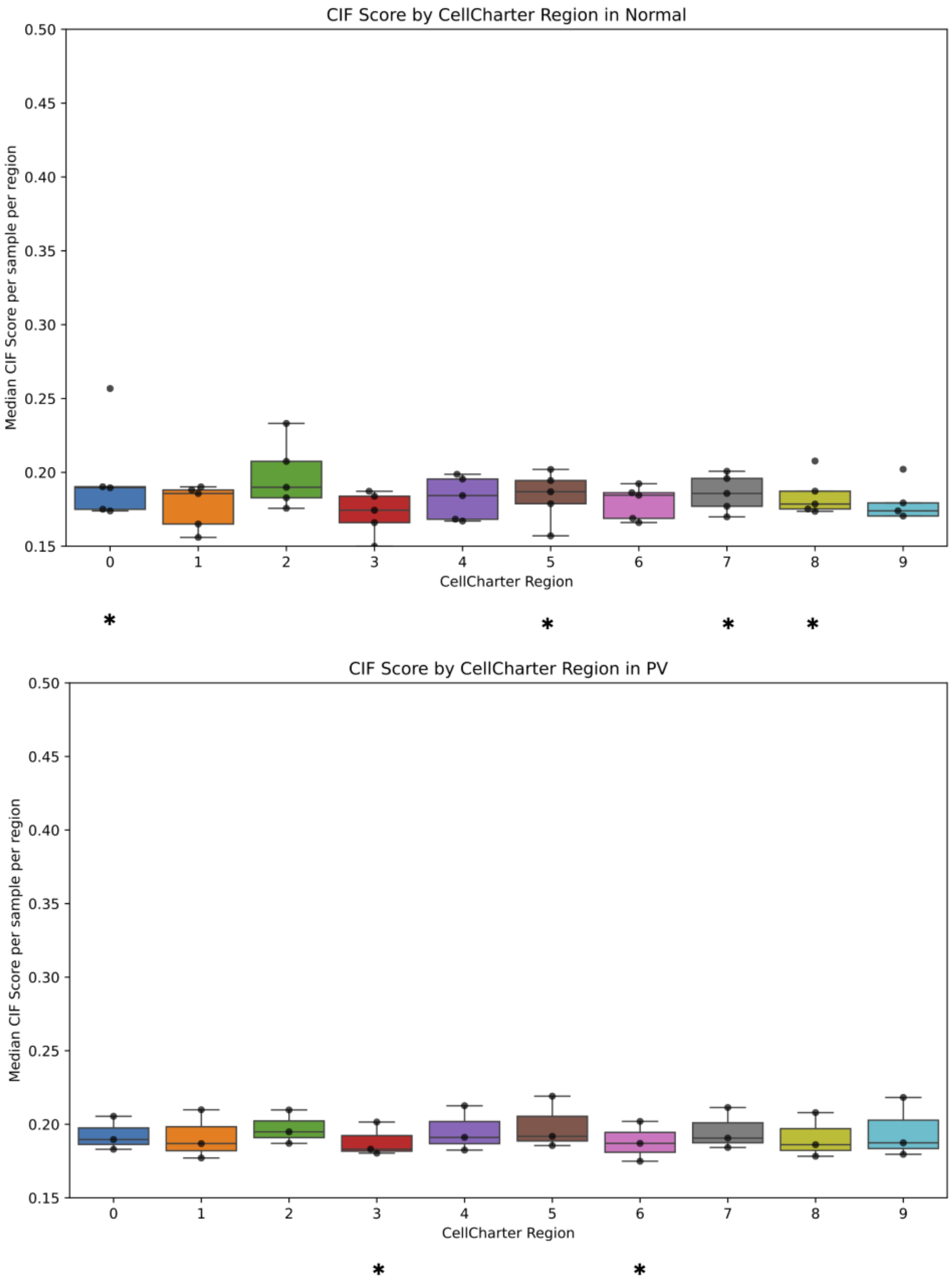


**Supplementary Figure 19: CIF score and cell neighbourhoods**
Boxplots to show the median corresponding CIF score for each CN per sample in (bottom) PV and (top) normal samples. *denotes that the median CIF score per CN was significantly different from CN2 (linear mixed effects model with Benjamini-Hochberg correction).

| Cell types | Spatiotypes cell type assignment |
|---|---|
| DC | MNP |
| Erythroid | Erythroid |
| Myeloid | Myeloid |
| Monocyte | MNP |
| B_cell | Lymphocyte |
| T_cell | Lymphocyte |
| Megakaryocyte | Megakaryocyte |
| Endothelial | Endothelial |
| Osteo-MSC | Osteo-MSC |
| HSC | HSC |
| Macrophage | MNP |
| Plasma_cell | Lymphocyte |
| GMP | Immature_myeloid |
| SMC | Stromal |
| Stromal | Stromal |
| Adipo-MSC | Stromal |
| Granulocyte/mast | Granulocyte/mast |

**Supplementary Data Figure 20: Table to show cell groups used for spatiotype analysis**
Cell groups were merged into broader lineage appropriate groups for the 'band descriptor' microenvironmental signature / spatiotype analysis. For example, monocytes, macrophages and dendritic cells were merged into a group termed mononuclear phagocytic (MNP) cells.

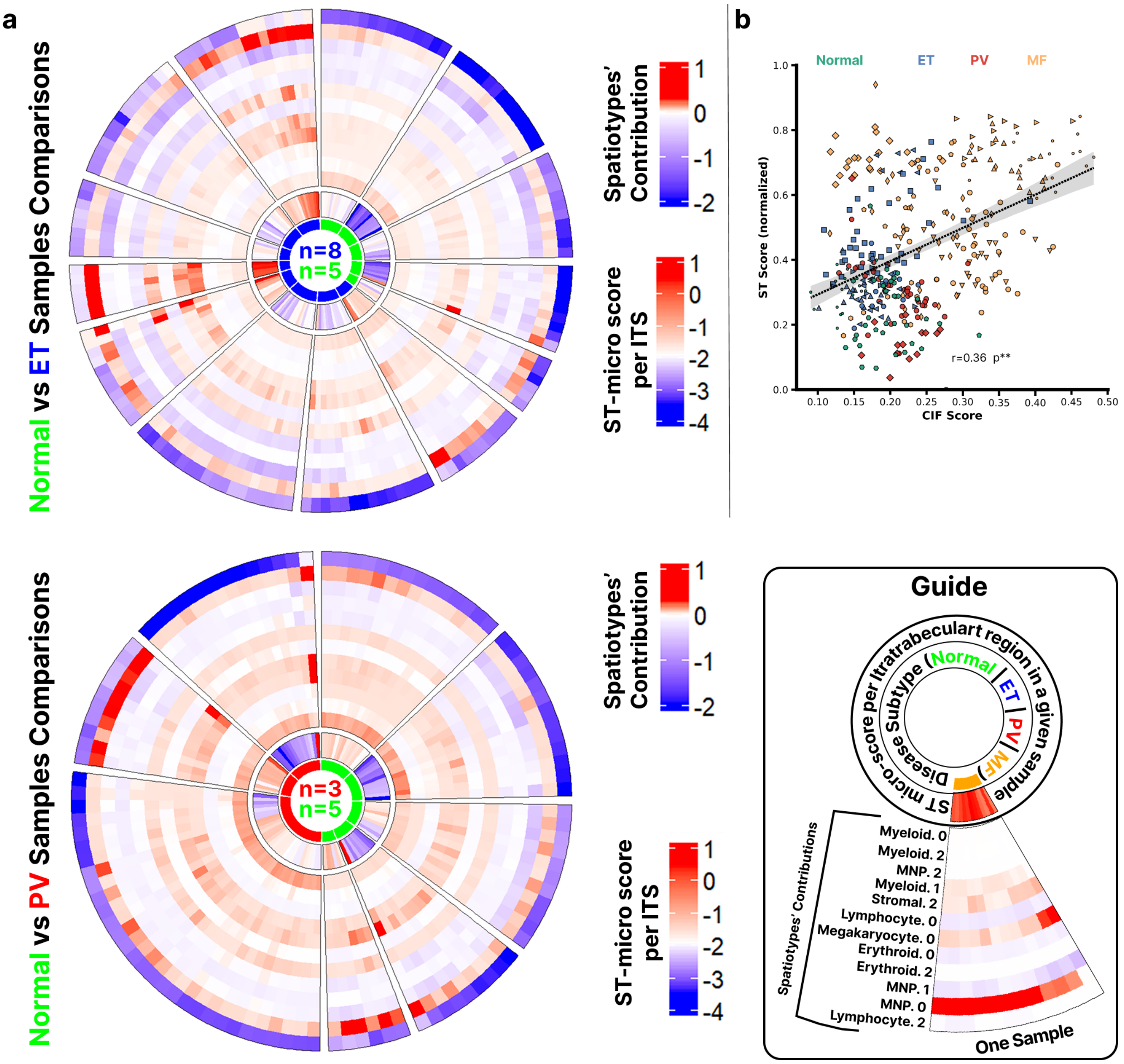


**Supplementary Data Figure 21: Microenvironmental spatiotypes capture microenvironmental heterogeneity**

**A:** Circle plots as per Figure 8F to demonstrate the spatiotypes per sample driving the overall ST-micro signature (center). The top circle plot represents the ET samples, and bottom circle plot PV with both including normal samples to allow for comparison. For a more detailed explanation, see Figure 8E and methods. **B:** Scatterplot to show the mean ST-micro score versus the CIF score per intertrabecular space. The points are coloured by condition, and shapes indicate sample (Pearson's correlation).
